# Cotranslational membrane insertion of the voltage-sensitive K^+^ channel KvAP

**DOI:** 10.1101/2024.05.28.596144

**Authors:** Justin M. Westerfield, Petra Kozojedová, Cara Juli, Ane Metola, Gunnar von Heijne

## Abstract

Voltage-sensor domains (VSDs), found in many voltage-sensitive ion channels and enzymes, are composed of four transmembrane helices (TMHs), including the atypical, highly positively charged S4 helix. VSDs are cotranslationally inserted into the membrane, raising the question of how the highly charged S4 helix is integrated into the lipid bilayer as it exits the ribosome. Here, we have used Force Profile Analysis to follow the cotranslational insertion of the six-TMH KvAP voltage-sensitive ion channel into the *E. coli* inner membrane. We find that the insertion process proceeds through three semi-independent steps: (*i*) insertion of the S1-S2 helix hairpin, (*ii*) insertion of the S3-S5 helices, and (*iii*) insertion of the Pore and S6 helices. Our analysis highlights the importance of the concerted insertion of helical hairpins, the dramatic influence of the positively charged residues in S4, and the unexpectedly strong forces and effects on downstream TMHs elicited by amphipathic and re-entrant helices.

## Introduction

The great majority of α-helical integral membrane proteins are cotranslationally inserted into their target membrane with the aid of a translocon. In the bacterium *Escherichia coli*, the translocon function is carried out by the SecYEG and YidC translocons located in the inner membrane (1), while in eukaryotic cells this function is carried out by the Sec61 and EMC complexes and accessory proteins in the rough endoplasmic reticulum (ER) (2). During the cotranslational insertion process, transmembrane α-helices (TMHs) partition into the membrane due to their overall hydrophobic character (3), while their in-out orientation relative to the membrane is largely dictated by flanking positively charged residues according to the “positive-inside” rule (4, 5).

While these basic features of membrane protein biogenesis have long been relatively well understood, it is only recently that methods have become available to follow the cotranslational membrane insertion process on a residue-by-residue level. Chief among those methods is Force Profile Analysis (FPA). FPA was first introduced as a method to follow the cotranslational insertion of model TMHs into the inner membrane of *E. coli* and, analogously, into ER-derived rough microsomes (RMs) (6). Subsequently, it has been applied to study the cotranslational folding of globular proteins (7, 8), cotranslational membrane translocation of proteins into the periplasmic space (9, 10), and cotranslational insertion of multispanning membrane proteins (11–13).

From the perspective of cotranslational membrane insertion, the so-called voltage-sensor domains (VSDs), first identified in voltage-sensitive ion channels (14), are of particular interest. VSDs are membrane-embedded domains composed of four TMHs, named S1-S4, and sense changes in the membrane potential via S4. Unique among known TMHs, S4 TMHs contain four to six conserved, positively charged residues that translate changes in the membrane potential into up-down movements of the S3-S4 “paddle” in the membrane (15). In voltage-sensitive ion channels, the movement of the paddle controls the opening and closing of an attached pore domain comprised of S5, a re-entrant helix named the Pore helix, and S6 (16). Cotranslational membrane insertion of VSDs poses a distinct problem due to the high charge density of S4, which has to eventually become embedded in the hydrophobic membrane environment (17).

In order to gain a deeper understanding of the cotranslational membrane insertion of VSDs, we have applied FPA to the archaeal voltage-sensitive K^+^ (Kv) channel KvAP from *Aeropyrum pernix*. KvAP was chosen because it is a prototypic tetrameric Kv channel of known structure (18) and the overall membrane insertion propensities of the S1-S6 TMHs have been measured (19). Our results unveil a complex insertion process, strongly influenced by the positively charged residues in S4.

## Results

### Force-profile analysis of cotranslational membrane insertion

Briefly, FPA measures forces on the nascent chain (NC) generated by the cotranslational membrane insertion and folding of TMHs, using so-called translational arrest peptides (APs) engineered into the NC as force-sensors (6, 20–25). The most commonly used AP for this application is from the *E. coli* SecM protein (Uniprot ID P62395, residues 150 to 166; hereafter called the SecM(*Ec*) AP). Immediately upon synthesis, the AP binds in the upper reaches of the ribosome exit tunnel and induces an altered active-site geometry in the peptidyl transferase center (PTC), preventing chain elongation (26). By application of an external force on the AP, however, it can be detached from its binding site, restoring the correct active-site geometry and allowing elongation to continue (6, 8, 26), Fig. 1a. *In vivo* expression, [^35^S]-Met pulse-labeling, immunoprecipitation, and separation by SDS-PAGE of constructs containing the SecM(*Ec*) AP thus result in two radioactive bands, corresponding to the arrested (A) and the full-length protein (FL), Fig. 1b. The higher the external pulling force applied on the NC at the point when the ribosome reaches the last codon of the AP, the larger will be the fraction of full-length protein, calculated as *f_FL_* = *I_FL_*/(*I_A_* + *I_FL_*), where *I_A_* and *I_FL_* are the integrated intensities of the arrested and full-length bands, respectively (6, 8).

**Figure 1.**
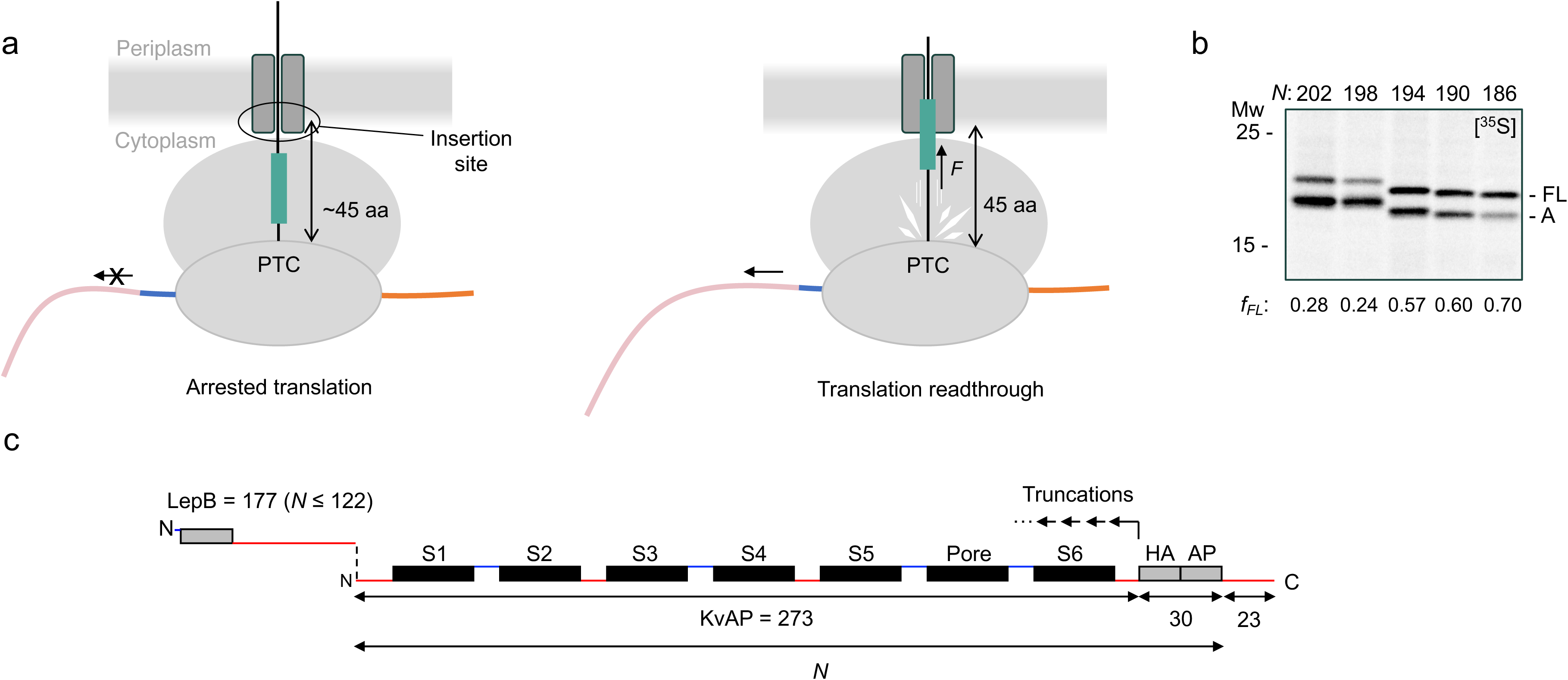
Force profile analysis. (a) When translating a construct harboring a target protein (pink), an arrest peptide (AP) (blue) and a C-terminal extension (orange), the ribosome will stall with the C-terminal residue of the AP in the PTC (left). When a sufficiently hydrophobic segment (green) reaches the insertion site ∼45 residues from the PTC, the stall is released by the force *F* generated by the segment partitioning into the membrane (right). (b) SDS-PAGE analysis of immunoprecipitated, [^35^S]-Met labeled constructs (identified by their *N* values, *c.f.* panel *c* and Fig. 2a). Stalling efficiency is quantified as the fraction of full-length protein (*f_FL_*), calculated from the integrated intensities of the full-length (FL) and arrested (A) bands. A 2’ labeling time was used throughout this work to optimize the dynamic range of the full FP (c.f., Fig. 2a). (c) The basic construct design, comprising full-length KvAP, an HA tag, the SecM(*Ec*) AP, and a 23-residue C-terminal extension. A series of constructs were generated by truncating four residues at a time from the C-terminal end of the KvAP segment; each construct corresponds to one data point on the FP (see Supplementary Materials for sequences; note that the C-terminal 53-residues are common to all constructs). For constructs with *N* ≤ 122, a 177-residue segment derived from the N-terminal part of the *E. coli* LepB protein (including its N-terminal TMH, in gray) was fused to the N-terminus of KvAP in order to ensure sufficient mass for separation in SDS-PAGE and proper membrane topology, c.f., Supplemental Fig. S1. Cytoplasmic and periplasmic loops are indicated by red and blue lines, respectively.

By analyzing a series of constructs where the SecM(*Ec*) AP is fused to progressively longer truncations of the protein of interest, a force profile (FP) can be generated by plotting *f_FL_* against the number of residues, *N*, measured from the construct’s N-terminus to the end of the AP, Fig. 1c. The FP thus represents the variation in the instantaneous force exerted on the AP during translation. High pulling forces are generated during the cotranslational partitioning of TMHs into the membrane, and the FP thereby provides a detailed picture of how different parts of the protein interact cotranslationally with the translocon/membrane system as they emerge from the ribosome (13).

### The KvAP force profile

We determined the full FP for KvAP at four-residue resolution (except for the region *N* = 150-166, which was mapped at one-to-two-residue resolution), Fig. 2a, using a series of constructs schematically represented in Fig. 1c (see Supplemental Table S1 for sequences of all constructs). Note that the 30-residue HA/AP part has the same sequence in all constructs and fills up most of the exit tunnel, as judged from both biochemical data (27), FPA data on cotranslational folding of water-soluble proteins (28), and the structure of a SecM-stalled ribosome-nascent chain complex (29) (Supplemental Fig. S1). Thus, because the sequence present inside the exit tunnel is the same in all constructs, the variation in pulling force recorded in the FP mainly reflects interactions between the NC and the translocon/membrane environment. In order to be able to visualize the shortest constructs (*N* ≤ 122 residues) on the SDS-PAGE gels, a 177-residue part of the *E. coli* inner membrane protein LepB was fused to their N-terminus, as indicated in Fig. 1c (see Materials and Methods); as shown in Supplemental Fig. S2, the FP obtained with the LepB fusion tends to give slightly higher *f_FL_* values (the average difference is 0.06) but does not differ significantly in shape over the tested region *N* = 110-166 residues. Similar control experiments in an earlier study of the GlpG and BtuC inner membrane proteins (13) also showed not significant differences ±LepB, and the replacement of the H2 transmembrane helix in LepB with a 19-residue segment composed of 8 Leu and 11 Ala residues likewise was found to have no effect on *f_FL_* for a model membrane protein construct (6).

**Figure 2.**
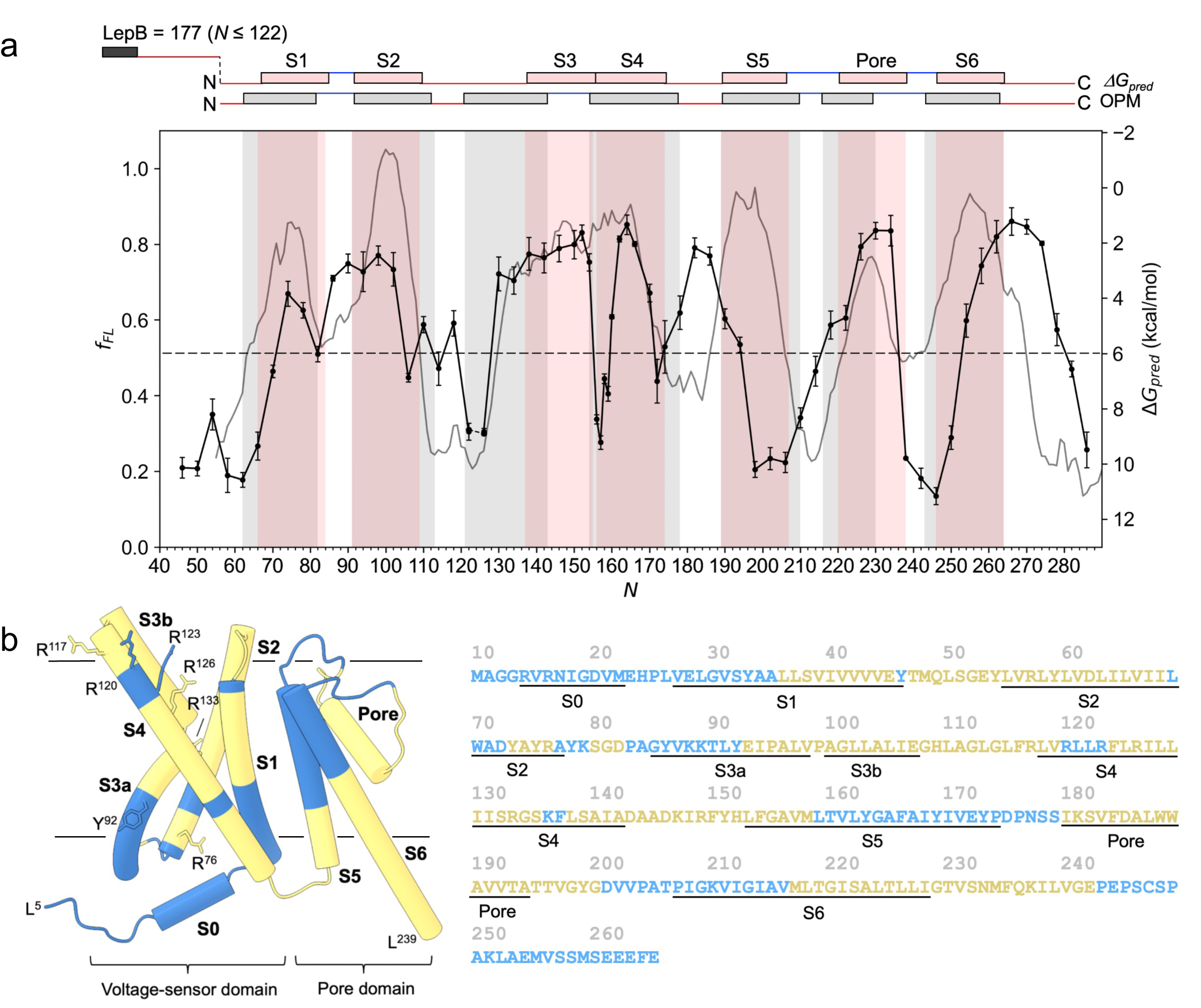
The KvAP force profile. (a) KvAP FP (black; error bars show SEM values), plotted alongside the predicted free energy of membrane insertion (*ΔG_pred_*; gray). *ΔG_pred_* was calculated as in (3) for 19-residue wide sequence segments whose centers are 45 residues upstream from the PTC at the corresponding value of *N*. Vertical shadings show predicted transmembrane domains (also shifted by 45 residues relative to the PTC) from the OPM (31) and *ΔG_pred_* algorithms (using a 19-residue window). (b) FP mapped onto the structure (18) and amino acid sequence of KvAP (PDB 6UWM). The S0 helix from the NMR structure PDB 2KYH (35) was manually added to the structure. A cutoff of *f_FL_* = 0.51 (midway between the highest and lowest *f_FL_* value in the FP; horizontal dashed line in panel *a*) was selected to define *N_start_* and *N_end_* values; regions with *f_FL_* values higher than the cutoff are in yellow, while those below the cutoff are in blue. Approximate membrane boundary locations are indicated as black lines according to the OPM database (cytoplasm down). The sequence used in this study is the same as in PDB 6UWM, where the initiator Met residue is counted as residue number 10; for ease of reference, we use the PDB numbering when referring to individual residues in the text and therefore subtract 45 - 9 = 36 residues from a given *N* value when referring to the residue 45 residues away from the PTC. The N-terminal sequence differs from the corresponding UniProt entry Q9YDF8, which matches the sequence used here from residue G^12^ onwards (numbered G^25^ in UniProt). OPM-predicted transmembrane domains are underlined. S0 is also underlined as defined in the NMR structure of the KvAP VSD (35).

We previously found that a model 19-residue TMH generates a peak in the FP that reaches half-maximal amplitude at an *N* value (*N*_start_) where its N-terminal end is ∼45 (±3) residues away from the PTC, and drops to half-maximal amplitude at an *N* value (*N*_end_) where its C-terminal end is ∼45 residues from the PTC (6); similar *N*_start_ and *N*_end_ values have been found for TMHs in other inner membrane proteins, both by FPA (10, 13) and a real-time FRET assay (30). Furthermore, if we overlay the KvAP FP and the predicted *ΔG* of insertion (*ΔG_pred_*, water-to-bilayer) free energy profile (3) calculated for the 19-residue sequence window centered 45 residues from the PTC, Fig. 2a, the peak positions correlate, with some exceptions. This indicates that, during cotranslational membrane insertion, the distance between the PTC in the ribosome and the membrane insertion site in the translocon can be bridged by a ∼45 residue-long, mostly extended polypeptide. The FP thus reflects membrane insertion events taking place ∼45 residues from the PTC, with a small uncertainty stemming from the fact that the conformation of the ∼15 residues between the constant HA/AP part and the insertion site may not be the same for all constructs (e.g., short helical segments could potentially form, depending on the sequence). However, this variation in the physical length of the NC should be small, for two reasons: (i) as soon as a sizeable pulling force is generated on the NC, the variable segment should adopt a more or less extended conformation regardless of sequence, (ii) segments corresponding to loops between TMHs generally have low helix propensities, and TMH segments need to be somewhere between 10-20 residues long to form helices in the exit tunnel even in the absence of a pulling force (24). Because the variable segment between the HA/AP and the insertion site is only ∼15 residues long, it is unlikely to have a sufficiently strong helix potential to markedly affect the ∼45 residues length needed to bridge the PTC and the insertion site.

Hence, by shifting the sequence by 45 residues relative to the FP, we can visualize the FP on the sequence and on the structure, Fig. 2b. Since the *f_FL_* values in the FP vary between 0.15 and 0.87, *i.e*., the half-maximal amplitude is *f_FL_* = 0.51 (dashed line in Fig. 2a), we have colored all residues with *f_FL_* ≥ 0.51 yellow and those with *f_FL_* < 0.51 blue in Fig. 2b. The *f_FL_* = 0.51 cutoff is also used to calculate the *N_start_* and *N*_end_ values below.

In the KvAP FP, Fig. 2a, the first peak has *N_start_* = 71±2 residues and *N*_end_ = 82±2 residues. A^35^ (residue numbering as in PDB file 6UWM) is ∼45 residues away from the PTC at *N_start_* and Y^46^ is ∼45 residues away from the PTC at *N*_end_, roughly matching the C-terminal half of the membrane-embedded part of S1 predicted by the hydrophobicity-based *ΔG*_pred_ and structure-based OPM (Orientation of Proteins in Membranes) algorithms (3, 31), Fig. 2b. The second peak has *N_start_* = 83±2 residues and *N_end_* = 105±2 residues, with T^47^ in the S1-S2 loop and L^69^ at the end of the hydrophobic core of S2 placed ∼45 residues away from the PTC, respectively. The peak has a shoulder that extends to *N* = 120±2 residues, apparently corresponding to the membrane insertion of the mildly apolar C-terminal end of S2 (W^70^-A^77^) and the S2-S3 loop (Y^78^-A^84^).

The S3–S5 region generates a more complex series of peaks, between *N* ≈ 128-194 residues. At *N_start_* = 128±2 residues, Y^92^ in the middle of S3a (the first half of S3) is ∼45 residues away from the PTC; notably, the preceding part of S3a (blue in Fig. 2b) is polar and not expected to generate pulling force. There is a sharp drop in *f_FL_* at *N* = 155±1 residues, coinciding with the first two Arg residues in S4 (R^117^, R^120^) arriving at ∼45 residues from the PTC. The next peak has *N_start_* = 161±1 residues, with L^125^ located ∼45 residues from the PTC. L^125^ is at the beginning of a hydrophobic stretch in the middle of S4 (F^124^LRILLIIS^132^) that presumably is responsible for the increase in *f_FL_*. The peak ends at *N_end_* = 171±2 residues, corresponding to a short polar stretch near the end of S4 (R^133^GSK^136^) being ∼45 residues from the PTC; at this point, all the 5 Arg residues in S4 should have reached the translocon/membrane insertion site. Finally, there is an unexpected peak between *N_start_* = 174±2 residues (with L^138^ ∼45 residues from the PTC) and *N_end_* = 194±2 residues. The segment from approximately G^134^ to D^146^ forms the amphipathic C-terminal part of S4 that is thought to “kink-and-slide” along the membrane surface during channel closing (32, 33), suggesting that the peak reflects the integration of the apolar face of this part of S4 into the cytoplasmic leaflet of the membrane.

Surprisingly, there is no peak in the FP corresponding to the relatively hydrophobic C-terminal two-thirds of the S5 TMH, but a major peak that extends from *N_start_* = 216±2 to *N_end_* = 236±2 residues that coincides with the amphipathic and mildly apolar Pore segment (encompassing the Pore helix and the selectivity filter residues) reaching ∼45 residues from the PTC. The final peak, with *N_start_* = 253±2 and *N_end_* = 280±2 residues, starts when M^217^, located near the middle of the apolar part of S6, is ∼45 residues from the PTC, and ends when E^244^ at the C-terminal end of S6 is ∼45 residues from the PTC; notably, the last part of S6 (G^229^TVSNMFQKILV^240^) is amphipathic and sticks out of the membrane, and partakes in the formation of the tetramer interface in the cryo-EM structure (PDB 6UWM). Similar to the C-terminal part of S4, the last part of the S6 peak likely reflects the integration of the amphipathic helix into the membrane interface region.

### Upstream TMHs affect the membrane insertion of neighboring TMHs in KvAP

In previous studies, it was shown that certain KvAP TMHs that are not sufficiently hydrophobic to insert on their own (*e.g*., S3) can nevertheless insert as helix pairs, or “hairpins” (19). To probe such insertion events in more detail, we recorded partial FPs for constructs where one or more N-terminal TMHs had been deleted and replaced by an N-terminal fragment of the LepB protein (see Materials and Methods), an inner membrane protein with a periplasmic N terminus, two N-terminal TMHs, and a large C-terminal periplasmic domain (34). For KvAP constructs lacking an odd number of TMHs we used a 232-residue LepB fragment that includes both TMH1 and TMH2 to ensure correct topology of the remaining KvAP TMHs in the membrane; likewise, for KvAP constructs lacking an even number of TMHs we used a 177-residue LepB fragment that includes only TMH1, Fig. 3 (FPs for the different deletions are plotted separately in Supplemental Fig. S3).

**Figure 3.**
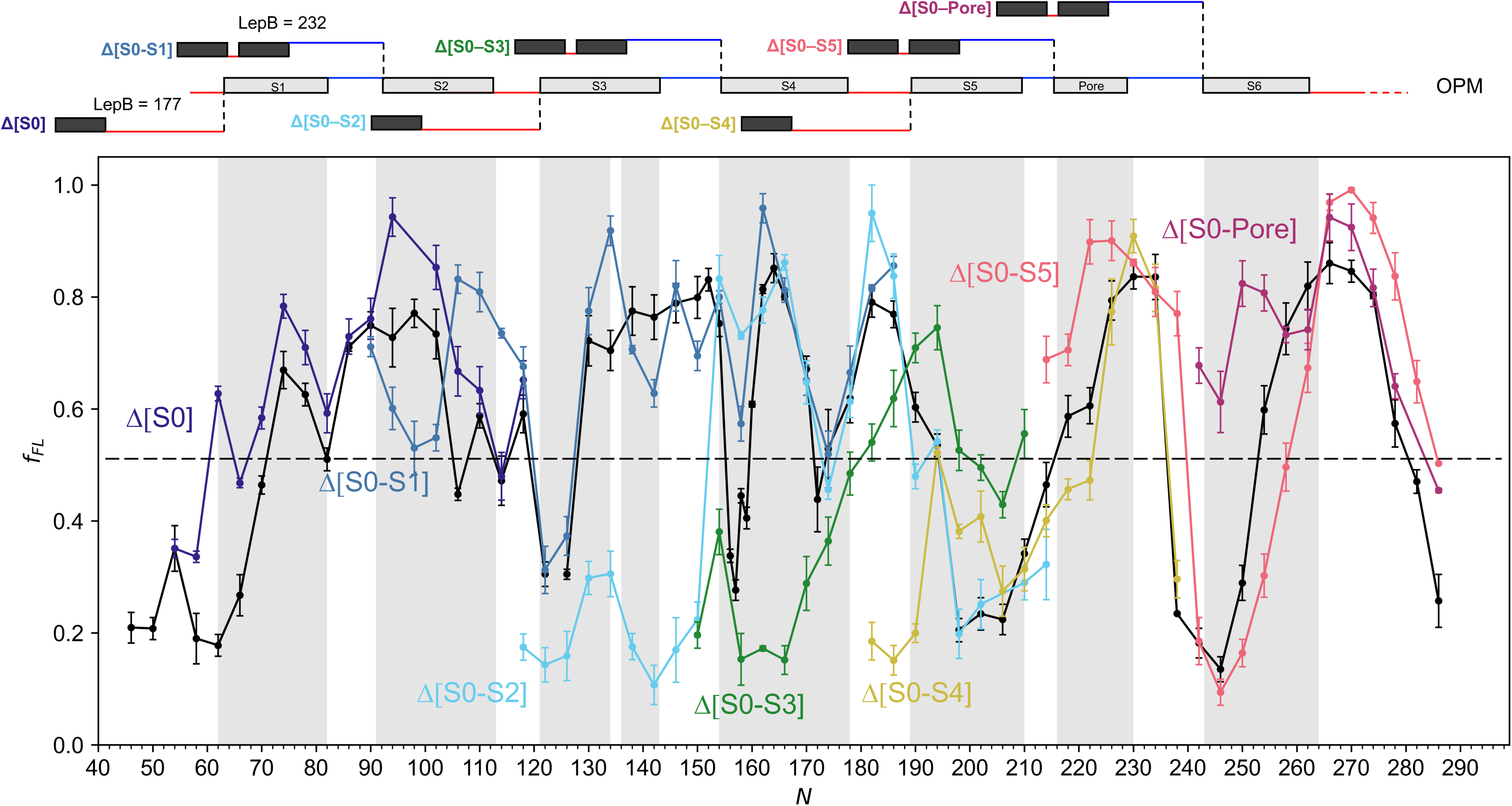
FPs for serial TMH deletion mutants. FPs are plotted alongside the KvAP FP from Fig. 2 (black). OPM-predicted transmembrane regions are shaded. The schematic above outlines the various construct designs used to maintain proper topology: a LepB-TMH1-THM2 fragment (232 residues) was fused to the N terminus of constructs lacking an odd number of TMHs and a LepB-TMH1 fragment (177 residues) was fused to constructs lacking an even number of TMHs. Separate plots of the individual deletion FPs are included in Supplemental Fig. S3.

We first deleted residues 10-18, that include most of the short amphipathic N-terminal S0 helix (R^14^-M^22^) seen in the NMR structure of the KvAP VSD (35). The Δ[S0] FP (dark blue) is similar to the wildtype KvAP FP, except that the S1 peak has *N_start_* = 62±2 residues (rather than 71±2 residues in the wildtype FP) and the S2 peak has a higher amplitude. Thus, in the Δ[S0] FP, the S1 peak starts when L^26^ at the very N terminus of S1 reaches ∼45 residues from the PTC, *i.e*., in the wildtype protein, the presence of the S0 amphipathic helix delays the interaction between S1 and the insertion site by ∼10 residues.

The Δ[S0-S1] series of constructs yielded the FP shown in blue in Fig. 3. The dip at *N* = 94-102 residues corresponds to a situation where R^57^ and D^62^, two polar residues that interrupt the hydrophobic N-terminal half of S2, are located ∼45 residues from the PTC. This dip is not seen in the original FP (Fig. 2a), which suggests that S1 helps to accommodate the S2 polar residues in the membrane. In contrast, the C-terminal half of S2 generates a stronger pulling force in the Δ[S0-S1] FP than in the original FP, and S3 generates a more variable pulling force with a peak at *N* = 134 (with the kink between S3a and S3b ∼45 residues from the PTC) and dips at *N* = 142 (with the polar residues E^107^GH^109^ at the C-terminal end of S3b ∼45 residues from the PTC) and *N* = 150 (with G^112^LG^114^ that eventually form the loop between S3b and S4 located ∼45 residues from the PTC).

The Δ[S0-S2] FP (light blue) has a low-amplitude peak at *N* = 130-134, where P^95^ and P^99^, which form the kink between S3a and S3b, are ∼45 residues away from the PTC. The Δ[S0-S2] FP has low *f_FL_* values in the hydrophobic S3b region, suggesting that S3 does not insert into the membrane in the absence of S1 and S2. A major peak with *N_start_* = 152±2 residues (where F^116^ at the N-terminus of S4 is ∼45 residues from the PTC) corresponds to the membrane insertion of S4; interestingly, however, there is no sharp drop at *N* ≈ 155 residues, unlike in the original FP.

For the Δ[S0-S3] FP (green), there is a low-amplitude peak at *N* = 154±2 residues but the high-amplitude peak in the original FP at *N_start_* = 161±1 residues (corresponding to the membrane insertion of the central apolar part of S4) is missing. Instead, *f_FL_* increases gradually from *N* ≈ 170 residues (where G^134^, which is located just before the C-terminal amphipathic part of S4, is ∼45 residues from the PTC) to a maximum at *N* = 194±2 residues; at this point M^158^ in the early part of S5 is ∼45 residues from the PTC. Thus, the passage of the main part of S4 into the translocon does not generate pulling force in the absence of S3, consistent with the relatively low insertion efficiency into rough microsomes seen for the isolated KvAP S4 helix (17, 19). Assuming that the isolated S4 helix is mostly translocated across the membrane, the “ramp” between *N* ≈ 170-194 residues corresponds to the entry into the translocon of the amphipathic C-terminal end of S4, the S4-S5 linker, and the apolar N-terminal third of S5.

The Δ[S0-S4] FP (yellow) has a mid-amplitude peak between *N_start_* = 192±2 residues and *N_end_* = 204±2 residues that corresponds to the N- and C-terminal ends of S5 reaching ∼45 residues from the PTC. Interestingly, unlike in the original FP, this peak agrees better with both the *ΔG*_pred_- and OPM-predicted transmembrane region for S5. The following high-amplitude peak at *N_start_* = 222±2 residues matches the original FP and corresponds to the membrane insertion of the Pore segment.

In the Δ[S0-S5] FP (pink), only the Pore segment and S6 remain of KvAP. Nevertheless, the peaks in the FP corresponding to the insertion of the Pore segment and S6 are similar to those in the full-length FP, except that *N_end_* for the Pore peak and *N_start_* and *N_end_* for the S6 peak are all increased by 4-5 residues.

Finally, in the Δ[S0-Pore] FP (purple), the peak has *N_start_* = 248±2 residues (with I^212^ at the N-terminal end of S6 ∼45 residues from the PTC), and has approximately the same *N_end_* value as seen in the full-length FP.

Thus, upstream TMHs affect the pulling forces generated by downstream TMHs for the whole S1-S5 region, but do not much affect those generated by the Pore segment and S6. These results are broadly in agreement with, and add considerable detail to, previous studies that have shown that KvAP S1 and S2 can insert independently into the membrane of ER-derived RMs, but that S3 cannot insert in the absence of S4, and S4 inserts inefficiently by itself (17, 19).

### Detailed analysis of the VSD FP

Given the complexity of the FP in the S4 region, we carried out two additional sets of experiments: first, recording a FP in the presence of 2 mM indole in the growth medium, a treatment known to reduce the proton-motive force (pmf) across the inner membrane (9, 36), and, second, recording FPs for mutated constructs in which either the positively charged Arg residues in S4 are mutated to positively charged Lys, negatively charged Asp, or polar but uncharged Gln, or the charged residues E^45^, D^62^, D^72^, R^76^, E^93^, and E^107^ in S2 and S3 that could potentially interact with S4 during membrane insertion are mutated to Ala, Fig. 4 and Supplemental Fig. S4.

**Figure 4.**
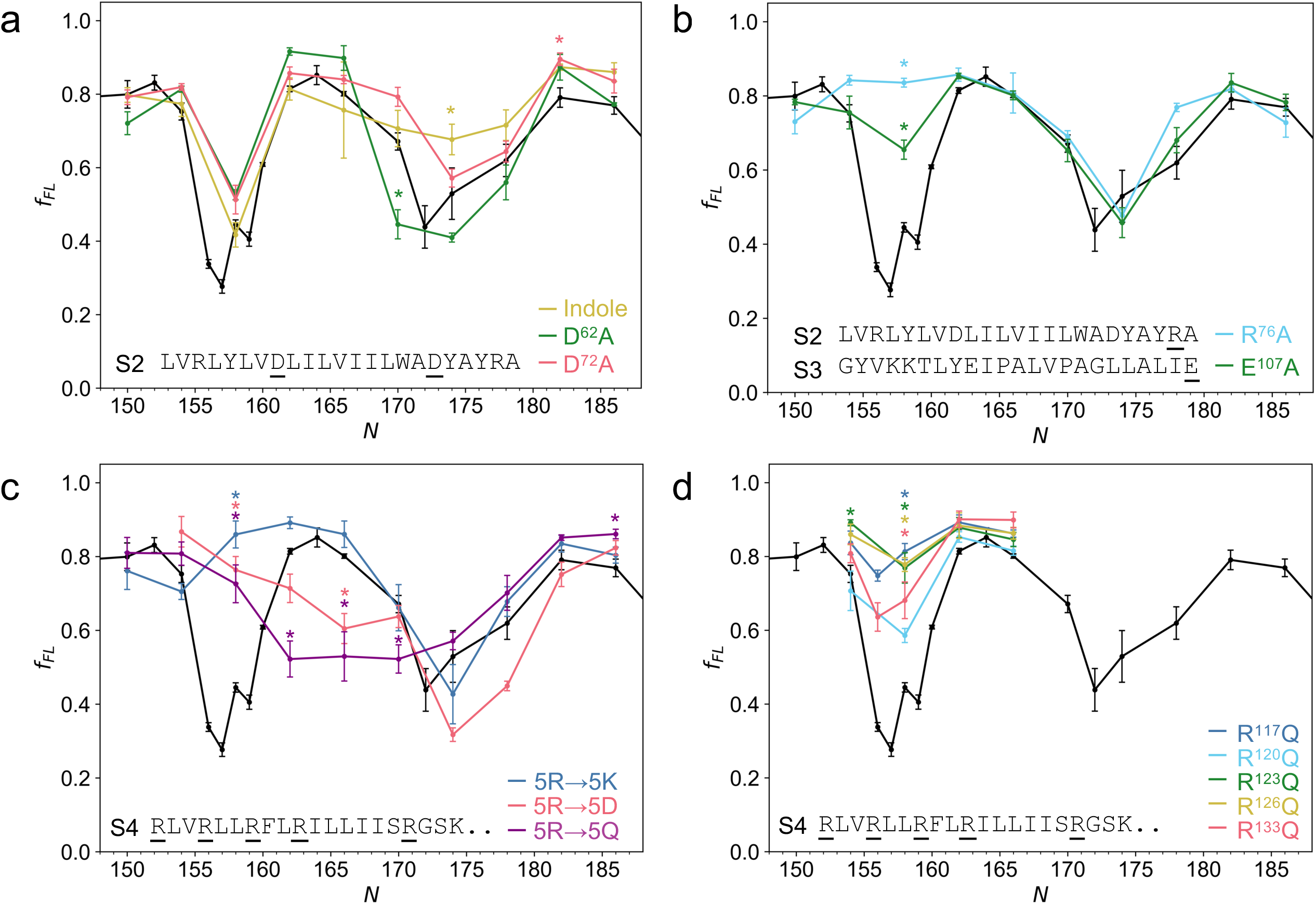
Mutational analysis of the S4 region of the FP. Each panel shows the S4 region of the KvAP FP in black alongside the FPs discussed in the main text. (a) FP expressed in the presence of 2 mM indole (yellow). FPs for the D^62^A and E^72^A mutants are shown in green and red, respectively. (b) FPs for the R^76^A (light blue) and E^107^A (green) mutants. (c) FPs for the [R^117^K, R^120^K, R^123^K, R^126^K, R^133^K] (blue), [R^117^D, R^120^D, R^123^D, R^126^D, R^133^D] (red), and [R^117^Q, R^120^Q, R^123^Q, R^126^Q, R^133^Q] mutants (purple). (d) FPs for the individual R^117^Q (blue), R^120^Q (cyan), R^123^Q (green), R^126^Q (yellow), and R^133^Q (red) mutants. Sequence contexts of the mutations are shown along the bottom of the panels. Significant differences (Dunnett’s test, p < 0.05) between mutant and wildtype *f_FL_* values are indicated by *.

Strikingly, the dip at *N* ≈ 174 residues disappears with indole in the medium while mutation D^62^A in the middle of S2 significantly deepens the dip at *N* = 170 residues, Fig. 4a; in both cases, the dip at *N* ≈ 157 residues is unaffected. The D^72^A mutation appears to reduce the dip at *N* ≈ 174 residues, but the only point that is significantly different from the wildtype FP is *N* = 182 residues. In contrast, mutations R^76^A at the C-terminal end of S2 and E^107^A in S3b completely eliminates (R^76^A) or strongly reduces (E^107^A) the dip at *N* ≈ 157 residues, but do not affect the dip at *N* ≈ 174 residues. Fig. 4b. Notably, R^76^ is not present in the Δ[S0-S2] series of constructs, for which the FP also does not have a dip at *N* ≈ 157 residues (Fig. 3). Mutations E^45^A and E^93^A have no significant effects on the FP, Supplemental Fig. S4.

With the five Arg residues in S4 simultaneously mutated to either Lys, Asp, or Gln, the dip at *N* ≈ 157 residues disappears in the corresponding FPs, Fig. 4c. In addition, the 5R→5D and 5R→5Q (but not 5R→5K) mutants significantly reduce the amplitude of the central peak between *N_start_* = 160±1 residues and *N_end_* = 172±2 residues, suggesting poor membrane interaction of these mutated S4 sequences, possibly because they cannot interact favorably with negatively charged residues in S2 (19). Single R→Q mutations in S4 reduce the dip at *N* ≈ 157 residues to somewhat varying degrees but have no effect on the central peak, Fig. 4d.

Thus, the sharp dip at *N* ≈ 157 residues is seen only when R^76^ at the cytoplasmic end of S2, E^107^ in S3b, and all five Arg residues in S4 are present, while the dip at *N* ≈ 174 residues requires an intact pmf across the inner membrane.

## Discussion

As it is clear from Fig. 2, the simple *ΔG_pred_* hydrophobicity plot provides at best a very rough guide to understand the cotranslational membrane insertion of KvAP. FPA goes further, both in terms of identifying regions in the protein for which the insertion process does not reflect the hydrophobicity profile, and because of its high resolution which allows the effects of individual residues on the insertion process to be discerned.

Based on the FPA data and the “sliding” model for membrane protein insertion (37), we propose the following scheme for the insertion process, Fig. 5. S1 starts to insert when approximately half of it has reached the translocon/membrane insertion site, ∼45 residues away from the PTC (Fig. 2a, b); in contrast, if the S0 amphipathic helix is deleted, S1 starts to insert as soon as its N-terminal end reaches the insertion site (Fig. 3). The continued partitioning of S1 into the cytoplasmic leaflet of the inner membrane generates high pulling force on the NC until the charged S1-S2 loop enters the insertion site (seen as the small dip in the FP at *N* ≈ 82 residues). When the first few apolar residues in S2 arrive at the insertion site (*N* ≈ 85 residues), *f_FL_* starts to increase again and remains high even though S2 has two charged residues (R^57^, D^62^) in its N-terminal half. Presumably, this is because the highly hydrophobic C-terminal part of S1 (V^39^IVVVV^44^) moves deeper into the membrane in step with the N-terminal part of S2, Fig. 5a (left), thus offsetting the energetic cost of inserting the charged residues. This interpretation is consistent with the Δ[S0-S1] FP, where the dip expected from R^57^ and D^62^ is visible at *N* ≈ 94-102 residues (Fig. 3), as in this case there is no upstream apolar segment that co-inserts with S2, Fig. 5a (right). At *N* ≈ 122 residues, the force drops. By this length, the entire S1-S2 region is sufficiently far from the PTC to have access to the membrane interior. This implies that a few residues in the S2-S3 loop must reach the insertion site on the cytoplasmic side of the membrane to allow full membrane insertion of the S1-S2 hairpin.

**Figure 5.**
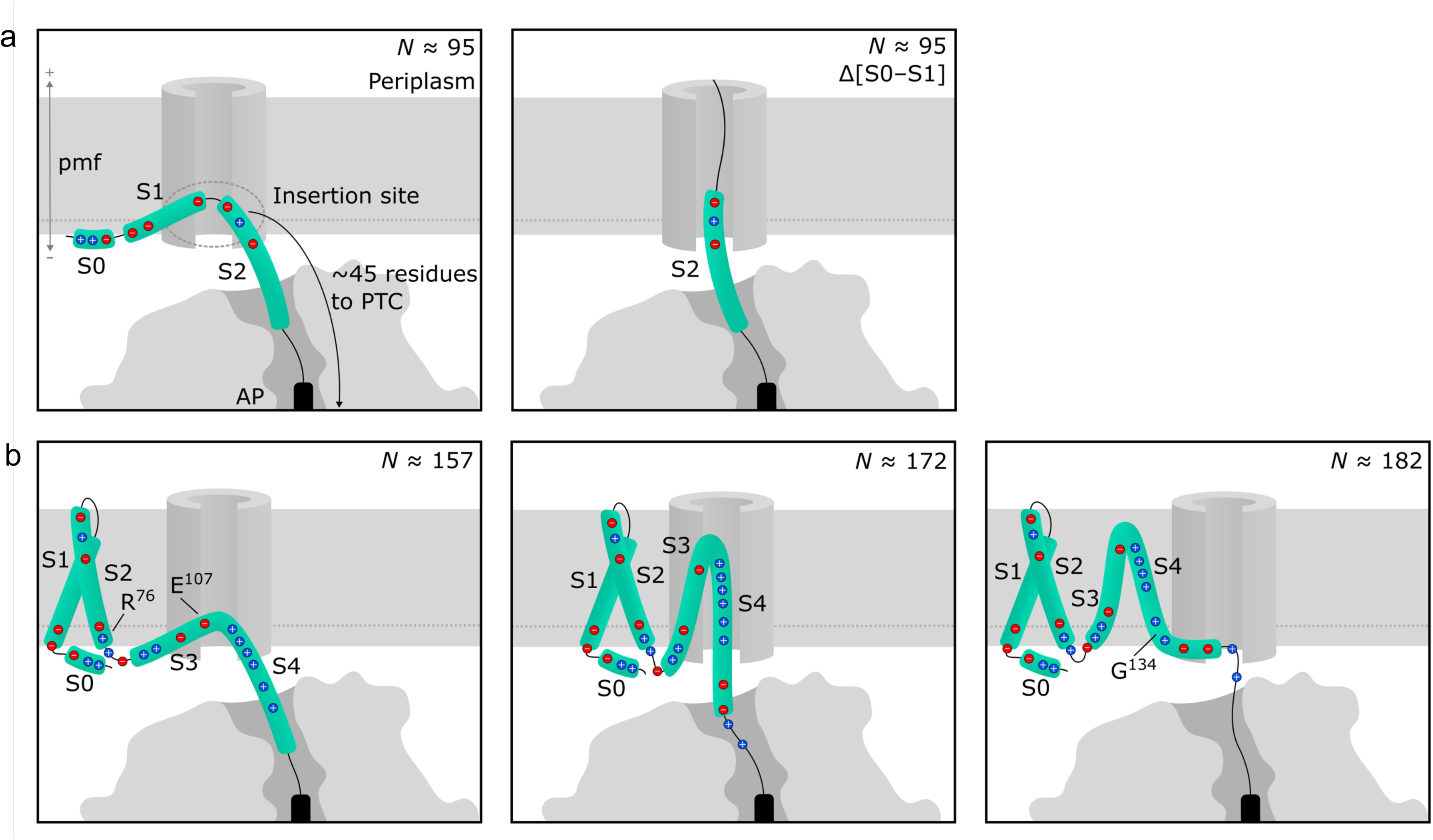
Model for the membrane insertion of KvAP. (a) The S1-S2 hairpin of KvAP generates high *f_FL_* when the N-terminus of S2 reaches the insertion site (*N* ≈ 95; left). In contrast, the Δ[S0-S1] mutant does not generate high *f_FL_* until more of S2 is exposed, possibly due to R^57^ and D^62^ (right). (b) The S3-S4 region in KvAP generates low *f_FL_* at *N* ≈ 157 (left) and *N* ≈ 172 (middle), when the positively charged stretches in S4 traverse the insertion site. In contrast, high *f_FL_* is generated at *N* ≈ 182 (right) when the ‘kinked’ end of S4 and the start of the S4–S5 linker reaches the insertion site. The SecM(*Ec*) AP is indicated in black. As the relative importance of the YidC and SecYEG translocons in the membrane insertion of KvAP is not known, only a “generic” translocon is depicted in the Figure.

Membrane insertion of the S3-S5 region is more complex, both because there is interdependency between TMHs (19), and because of the unique composition of S4 with five regularly spaced, positively charged Arg residues. In the cryo-EM structure (PDB 6UWM), S3 has a break at P^95^-P^99^, between the more polar S3a and more apolar S3b helices. S3 starts to insert into the cytoplasmic membrane leaflet at *N* ≈ 128 residues when the C-terminal end of S3a reaches the insertion site, and S3b keeps generating a high pulling force until *N* ≈ 155 residues when the first Arg residues (R^117^, R^120^) in S4 appear at the insertion site. As shown by the Δ[S0-S2] FP (Fig. 3), S3 generates a high pulling force only if S2 is present. In the Δ[S0-S2] constructs, S3 is preceded by a long, polar cytoplasmic segment from LepB and S3’s modest hydrophobicity (Fig. 2a) might not be sufficient for re-targeting the ribosome to the SecYEG translocon for efficient membrane insertion (38, 39). In agreement with these results, previous studies on KvAP (19) have also shown that, in the absence of S1 and S2, S3 does not insert by itself but only when part of the S3-S4 hairpin.

The sharp drop in *f_FL_* at *N* ≈ 155-160 residues appears to be almost entirely dependent on the S4 Arg residues. Mutating any or all of the five Arg residues to Gln results in increased *f_FL_* at *N* = 158 residues (Fig. 4c, d). An effect is even seen for R^133^Q, which is only ∼35 residues away from the PTC at this point and therefore presumably has not reached the insertion site. Mutating all of the Arg residues to Lys or Asp also results in increased *f_FL_* (Fig. 4c), while dissipation of the pmf with indole has no effect (Fig. 4a). Notably, Arg – but not Lys – residues have been observed to be rate-limiting for (apparently pmf-independent) translocon-mediated membrane translocation of proteins into *E.coli*-derived proteoliposomes or inverted inner-membrane vesicles (40), possibly due to the much higher pKa of the Arg side chain that prevents its de-protonation even in non-polar environments (41, 42).

We also tested several Ala substitutions of charged residues in S2 and S3 (Fig. 4b, Supplemental Fig. S4). Remarkably, mutations R^76^A (at the cytoplasmic end of S2) and E^107^A (near the C-terminal end of S3b) cause a significant increase in *f_FL_* at *N* = 158 residues; the same effect is seen in the Δ[S0-S2] FP, which lacks R^76^, and, albeit less pronounced, in the Δ[S0-S1] FP (Fig. 3). The E^107^A mutation increases the hydrophobicity of S3b, which may help the S3-S4 hairpin insert into the membrane, Fig. 5b (left). The effect of the R^76^A mutation is more difficult to understand. One possibility is that the increase in *f_FL_* at *N* = 158 residues caused by this mutation might be due to disruption of the native contacts between the C-terminal end of S2 (D^72^-K^79^) and the amphipathic N-terminal S0 helix (35). This scenario suggests a previously unobserved long-range interaction between S0, S2, and the tip of the S3-S4 hairpin during the early stages of membrane insertion of the latter, Fig. 5b (left).

Once the apolar segment in the middle of S4 (F^124^LRILLIIS^132^) arrives at the insertion site at *N* = 162 residues, there is a rapid increase in *f_FL_*. This peak is absent or diminished in the Δ[S0-S3], 5R→5Q, and 5R→5D mutants, but not in 5R→5K, nor in any of the single R→Q mutants (Fig. 3, Fig. 4c, d). Thus, this peak seems to minimally require S3 and one positive charge flanking the hydrophobic stretch (R^126^ILLIISR^133^). The 5R→5Q mutation increases *f_FL_* at *N* = 158 residues and decreases it at *N* = 162-166 residues, further demonstrating the intricate effects caused by the S4 Arg residues during membrane insertion, and the difficulty of predicting *f_FL_* from simple hydrophobicity plots.

A second dip in the S4 region of the FP at *N* ≈ 172 residues coincides with the arrival at the insertion site of two positively charged residues (R^133^, K^136^) at the end of the membrane-embedded part of S4, Fig. 5b (middle). This dip is not seen when constructs are expressed in the presence of indole (Fig. 4a), suggesting that it is caused by an electrostatic interaction between positively charged residues in S4 and the membrane potential. As discussed in Allen et al. (40), the role played by the pmf during membrane translocation is still not fully understood, and our finding that indole has no effect on the Arg-dependent dip at *N* ≈ 157 residues but does affect the dip at *N* ≈ 172 residues underscores this point. In contrast to indole, the D^62^A mutation deepens the dip at *N* = 170 residues, suggesting that D^62^ in the middle of S2 facilitates the insertion of the second half of S4; in agreement with this, a similar mutation (D^62^V) has been shown to reduce the overall membrane insertion efficiency of the S3-S4 hairpin (19).

The following peak (*N* ≈ 178-194 residues) is displaced by ∼10 residues compared to what would be expected for a peak generated by S5. Its position is unchanged in the Δ[S0-S2] FP but is shifted towards S5 in the Δ[S0-S3] FP and overlaps fully with the predicted membrane-embedded region of S5 in the Δ[S0-S4] FP (albeit with a low amplitude). The similarity between the original FP and the Δ[S0-S2] FP in this region rules out the possibility that the peak originates from a folding transition of the whole VSD after the final insertion of S4. Instead, the data suggest that the amphipathic C-terminal end of S4 and the early part of S5 are responsible: our working hypothesis is that S4 kinks near G^134^ such that the amphipathic part does not enter the translocon but rather partitions into the cytoplasmic leaflet of the membrane, Fig 5b (right), as proposed for the closed conformation of the KvAP channel (32).

The amphipathic Pore segment generates a strong pulling force, despite its rather low overall hydrophobicity. A similarly strong pulling force was previously recorded for an amphipathic re-entrant helix in BtuC (13). This suggests that NC tension has a similar effect on the ribosome whether it is induced by polypeptide segments destined to become TMHs or those destined to become re-entrant helices, meaning that a similar insertion mechanism is likely responsible for both cases. Since the re-entrant helix of BtuC and the Pore helix of KvAP are amphipathic, we propose that amphipathic helices “slide” across the membrane via the lateral gate region in the SecYEG translocon (or the cleft in the YidC translocon), with their polar side facing the translocon channel and their apolar side facing the surrounding lipid (37). In this way their apolar face can generate a strong pulling force while the polar face prevents partitioning into the lipid bilayer and formation of a TMH. Once the Pore helix has translocated across the membrane, its amphipathic character is also required for the proper folding of the KvAP monomer (43).

The Pore segment delays the insertion of S6 by 8-10 residues compared to the insertion of S6 by itself (the Δ[S0-Pore] FP, Fig. 3), and, likewise, the amphipathic S0 helix delays the insertion of S1 by a similar number of residues (the Δ[S0] FP, Fig. 3). As noted above, the amphipathic C-terminal end of S4 also seems to have an attenuating effect on the insertion of S5. Interestingly, a similar delay in the insertion of an immediate downstream TMH was seen for the amphipathic re-entrant helix in BtuC (13) and was traced to a stretch of hydrophobic residues in the re-entrant helix (11). Further studies will be required to understand the precise mechanism behind this effect; a speculative hypothesis is that amphipathic helices may bind transiently in the Sec61 channel and delay or block entry of a downstream TMH in a similar way that Sec61 inhibitors such as cotransin-related compounds (44) or mycolactones (45) block the entry of signal peptides.

On a more general level, for the serial TMH deletion mutants (Fig. 3) we find that the FP is most affected in the region immediately following the deletion. For example, in the Δ[S0-S1] deletion mutant the S2 region is most disturbed, the S3 region less so, and the S4 region less still. Hence, cotranslational insertion forces are primarily locally determined and correlate with the local segmental hydrophobicity, while major disagreements between the hydrophobicity profile and the FP suggest longer-range intra- or inter-molecular folding events (12). Notably, despite the fact that the purified monomeric S0-S4 paddle domain can fold by itself (16, 35) – and hence presumably also folds cotranslationally during membrane insertion – the only strong evidence for long-range interactions in our data are the effects of the D^62^A, R^76^A and Δ[S0-S1] mutations on the insertion of the early part of S4 (the dip at N ≈ 158 residues, Fig. 3).

In summary, the cotranslational membrane insertion of KvAP proceeds through three semi-independent steps: (*i*) insertion of the S1-S2 hairpin, (*ii*) insertion of the S3-S5 region, and (*iii*) insertion of the Pore-S6 region. Each stage presents a more complex FP than excepted from a simple hydrophobicity plot, highlighting the importance of concerted insertion of helical hairpins, the – sometimes dramatic – influence of positively charged residues, and the unexpectedly strong forces, and effects on downstream TMHs, elicited by amphipathic helical segments, *e.g*., S0, the C-terminal ends of S4 and S6, and the re-entrant Pore helix. Overall, the peculiar insertion characteristics of the S1-S4 domain are presumably shared between KvAP and VSDs in other voltage-sensitive ion channels, while the strong pulling forces generated by amphipathic helical segments may be a common phenomenon among multi-spanning helix-bundle membrane proteins. Thus, at the level of detail afforded by FPA, translocon-mediated membrane-protein insertion reveals itself as a much richer process than implied by the simple picture apparent from hydrophobicity plots or coarse-grained molecular dynamics simulations (13, 46).

## Materials and Methods

### Enzymes and chemicals

Cloning enzymes, reagents, and buffers were acquired from New England BioLabs (Ipswich, Massachusetts, U.S.) and ThermoFisher Scientific (Waltham, Massachusetts, U.S.). [^35^S] L-Methionine was ordered from PerkinElmer (Shelton, Connecticut, U.S.). All other chemicals were acquired from Sigma-Aldrich (subsidiary of Merck Life Science, Darmstadt, Germany) unless otherwise stated.

### Cloning and mutagenesis

We refer to the amino acid numbering presented in (18). Therefore, the first residue of K_V_AP in our constructs is numbered as residue 10. Constructs were designed to include various lengths of KvAP truncated from the C-terminus, an SGMG linker (both to flexibly isolate KvAP and to add an extra Met residue for more labeling), an HA-tag for immunoprecipitation, the SecM(*Ec*) AP, and a C-terminal fragment of LepB (residues 306-324) to allow separation of the arrested and full-length fragments by SDS-PAGE (Fig. 1c). In constructs in which KvAP is heavily truncated (*N* ≤ 122) we added an N-terminal fragment of LepB (residues 1-226, but excluding TM2 (residues 58-114) to ensure proper membrane orientation of the KvAP part). Not only does the added mass improve visualization by SDS-PAGE but the LepB fragments also promotes efficient cotranslational insertion of fusion proteins into the inner membrane (13). The added LepB residues are ignored in the calculation of the *N* values; see the Supplementary Materials for full sequences of all constructs. An initial set of 60 constructs was ordered from Invitrogen GeneArt Gene Synthesis Services (ThermoFisher Scientific, Waltham, Massachusetts, U.S.) in a pBAD/*Myc*-His A background (47). Further mutagenesis was either similarly ordered, or performed in-house by a combination of Q5 PCR and Gibson assembly (48) with a homemade reaction mixture.

### In vivo pulse-labeling analysis

For expression, constructs were transformed into BL21 (DE3) cells (49). Cultures (1-4 mL) were seeded to an OD_600_ of 0.1 in minimal media (1x M9 salts [M6030, Sigma-Aldrich, subsidiary of Merck, Darmstadt, Germany], 1 g/L complete supplement mixture without methionine (CSM-Met, 4510712, MP Biomedicals, California, U.S.), 0.1 mg/mL thiamine, 0.4% (w/v) fructose, 100 µg/mL ampicillin, 2 mM MgSO_4_, and 0.1 mM CaCl_2_). After reaching OD_600_ of 0.5, expression was induced with 0.2% arabinose for 5 minutes, then the cultures were radiolabeled by supplementation with 10-20 µCi [^35^S]-Met (NEG009T001MC, Perkin-Elmer, Shelton, Connecticut, U.S.) for an additional 2 minutes before stopping expression by precipitation with ice-cold trichloroacetic acid (10% final). For experiments using indole (Fig. 4a), 2 µL of a 1.5 M stock solution was added (final concentration 3 mM), one minute before adding [^35^S]-Met. The precipitate was washed with 1 mL of ice-cold acetone before resuspension.

Proteins were resuspended in 10 mM Tris, 2% SDS, pH 7.5. The suspension was centrifuged to remove insoluble material, then mixed with 10 µL of protein G-agarose beads (PROTGA-RO, Roche, Basel, Switzerland) and incubated for 15 minutes on ice to capture proteins that non-specifically bind to the beads. Then, the supernatant was transferred to a tube containing 10 µL of fresh beads and 1 µL of α-HA antibody (901515, BioLegend, San Diego, California, U.S.) and incubated under gentle agitation overnight at 4° C. Beads were washed first with 10 mM Tris, 150 mM NaCl, 2 mM EDTA, 0.2% (w/v) Triton X-100, then again with only 10 mM Tris. After removing as much buffer as possible, proteins were recovered from the beads by denaturation with sample buffer. Peptidyl-tRNA was then digested with RNAse A (400 µg/µL). Samples were run on NuPAGE 12% Bis-Tris pre-cast gels, fixed in 30% ethanol and 10% acetic acid, and then soaked in Gel-Dry Drying Solution (Invitrogen, ThermoFisher, Waltham Massachusetts, U.S.) before drying on a Hoefer GD 2000 gel dryer (Hoefer Inc., U.S.) for 1 hour at 65° C. Phosphorimaging plates (BAS-IP, Fujifilm, Tokyo, Japan) were exposed to gels for 2-3 days before imaging on a FLA-9000 imager (Fujifilm, Tokyo, Japan).

### Quantification

Full-length and arrested band intensity profiles were extracted with ImageJ. The profiles were then fit to one or more gaussian curves using an in-house python script to determine the integrated band intensities. *f*_FL_ was calculated as 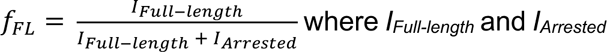 are the integrated intensities of the full-length and arrested bands, respectively. For certain constructs where band assignment was ambiguous, control constructs were cloned and expressed. Full-length controls were made by mutating the terminal proline of SecM(*Ec*) to alanine, which prevents SecM(*Ec*)-induced stalling. Arrested controls were made by mutating the terminal proline of SecM(*Ec*) to a stop codon.

### Statistical analysis

The significance of observed differences between the wildtype KvAP FP and FPs for the mutants reported in Fig. 4 was estimated by the multiple comparisons Dunnett’s test (*p* < 0.05) (50), using the Python (version 3.12) library SciPy (51). For the data in Supplemental Fig. S1, a Bonferroni-corrected t-test (52) was used to estimate the significance of observed differences between the FPs obtained with and without the N-terminal LepB fusion, and to correct for the average difference in *f_FL_* values (0.06) between the ±LepB FPs, 0.06 was added to the measured *f_FL_* values for the -LepB FP before performing the t-test.

## Acknowledgments

This work was supported by grants from the Knut and Alice Wallenberg Foundation (2017.0323), the Novo Nordisk Fund (NNF18OC0032828), the Swedish Research Council (2020–03238), and the Vallee Foundation (VF2021) to GvH. The plasmid encoding the KvAP sequence was a kind gift from Dr. Roderick MacKinnon, Rockefeller University.

## Supporting Information

**Supplemental Figure S1.**
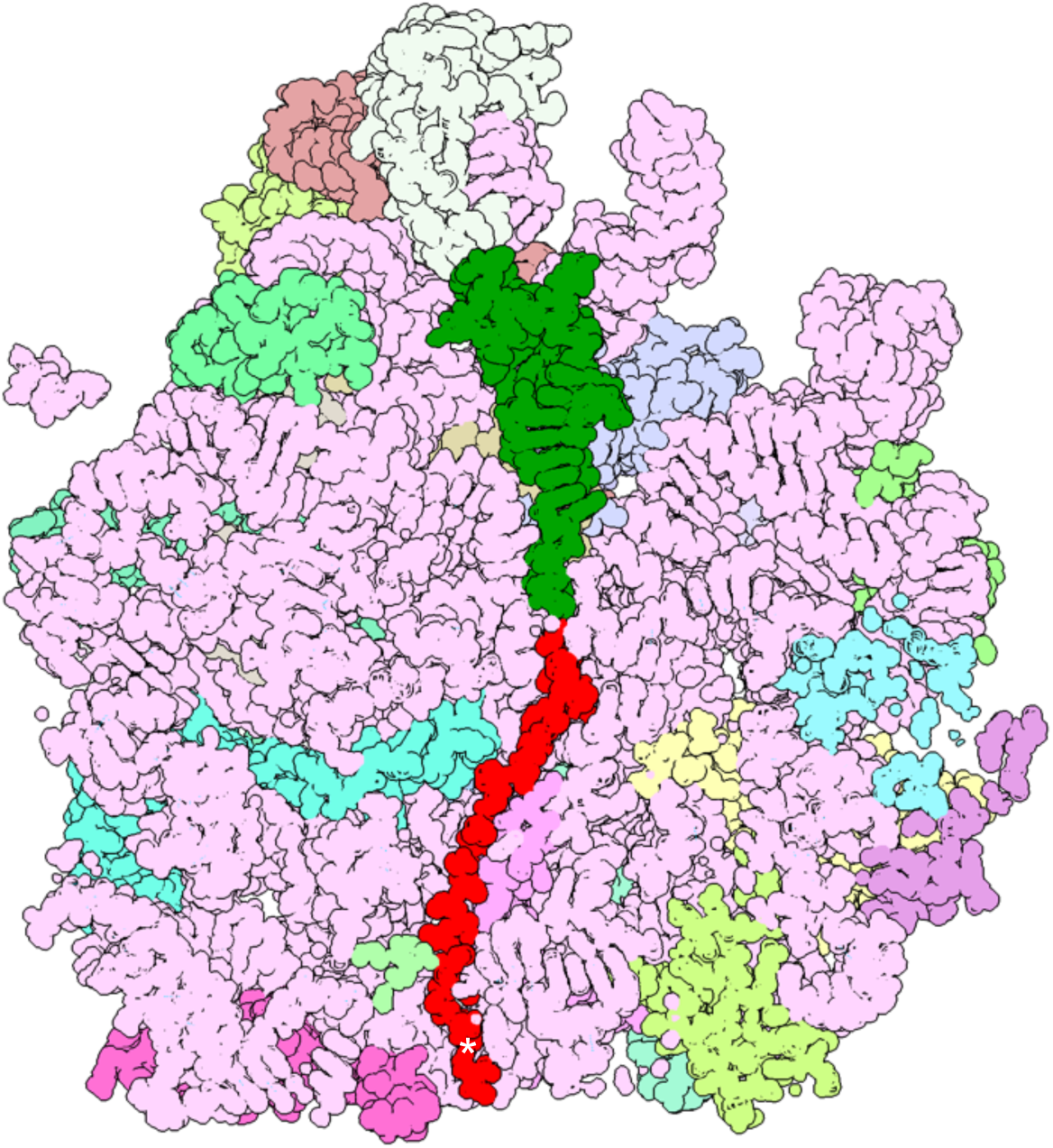
Structure of a SecM-stalled ribosome-nascent chain complex (Ahn M*, et al.* (2022) *Nat Commun* 13:4243. doi: 10.1038/s41467-022-31906-z; PDB 7ZP8). The visible part of the NC (in red) is 33 residues long and has an extend conformation; the NC also includes a folded N-terminal FLN5 domain (not visible in the structure) that generates a substantial pulling force on the NC. The N-terminal residue of the 30-residue HA/AP constant part in the KvAP constructs (*c.f*., Fig.1 in the main text) would correspond to residue E^137^ (marked by *). The SecM AP used is a mutant version of the wildtype *E. coli* SecM AP that causes stronger translational arrest (necessary to withstand the pulling force generated by the folding of the FLN5 domain). The P-site tRNA is in dark green, RNA is in pale pink, and other colors indicate ribosomal proteins. Image prepared in ChimeraX (Goddard TD*, et al.* (2018) *Protein Sci* 27:14-25).

**Supplemental Figure S2.**
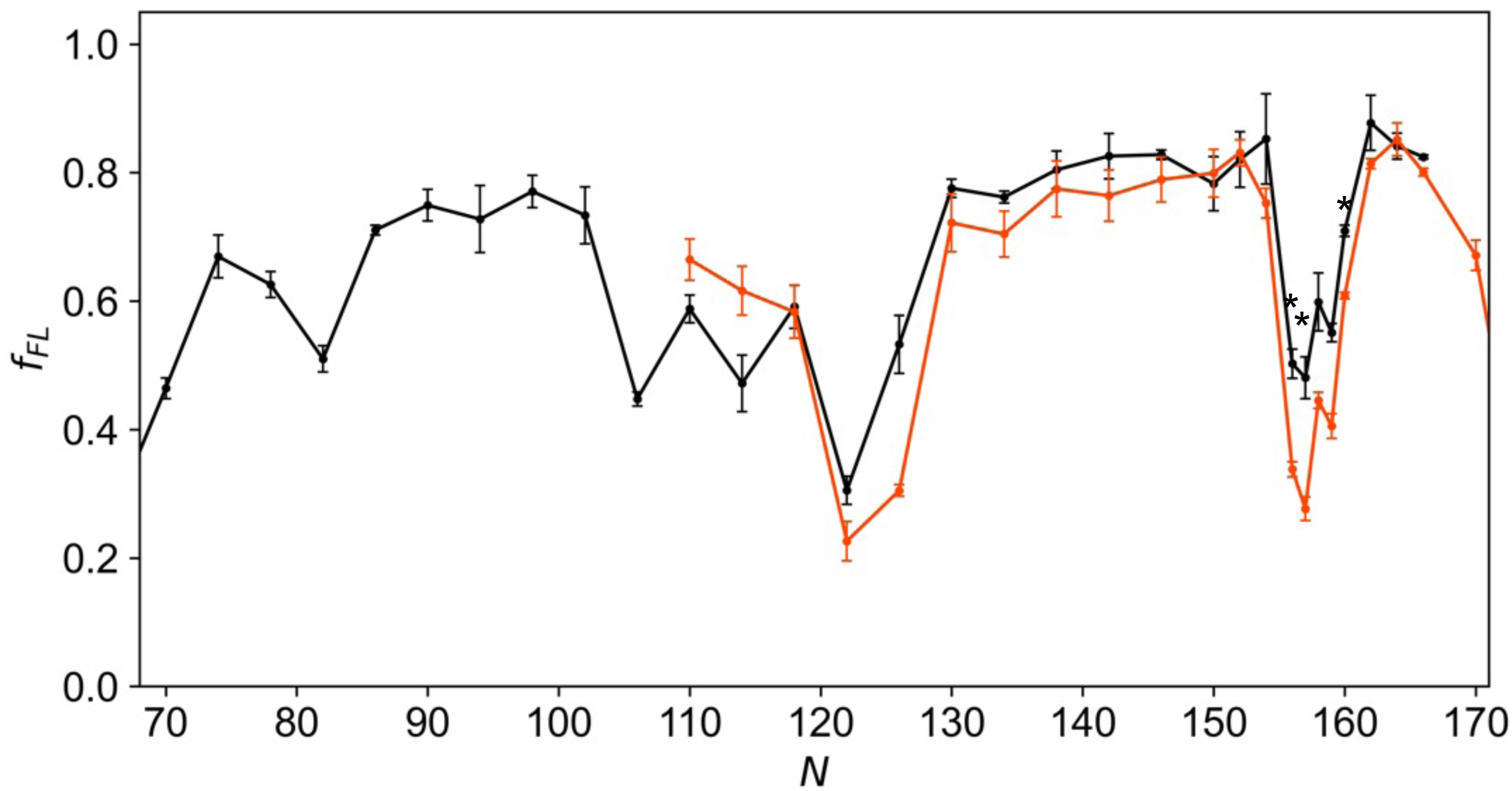
Overlapping KvAP FPs obtained with (black) and without (red) an N-terminal LepB (1–177) fusion (c.f., Fig. 2 in the main text). Significant differences (t-test with Bonferroni correction, *p* < 0.05) between mutant and wildtype *f_FL_* values are indicated by *. These differences become non-significant if a correction is made for the average difference of 0.06 in the *f_FL_* values for the ±LepB FPs.

**Supplemental Figure S3.**
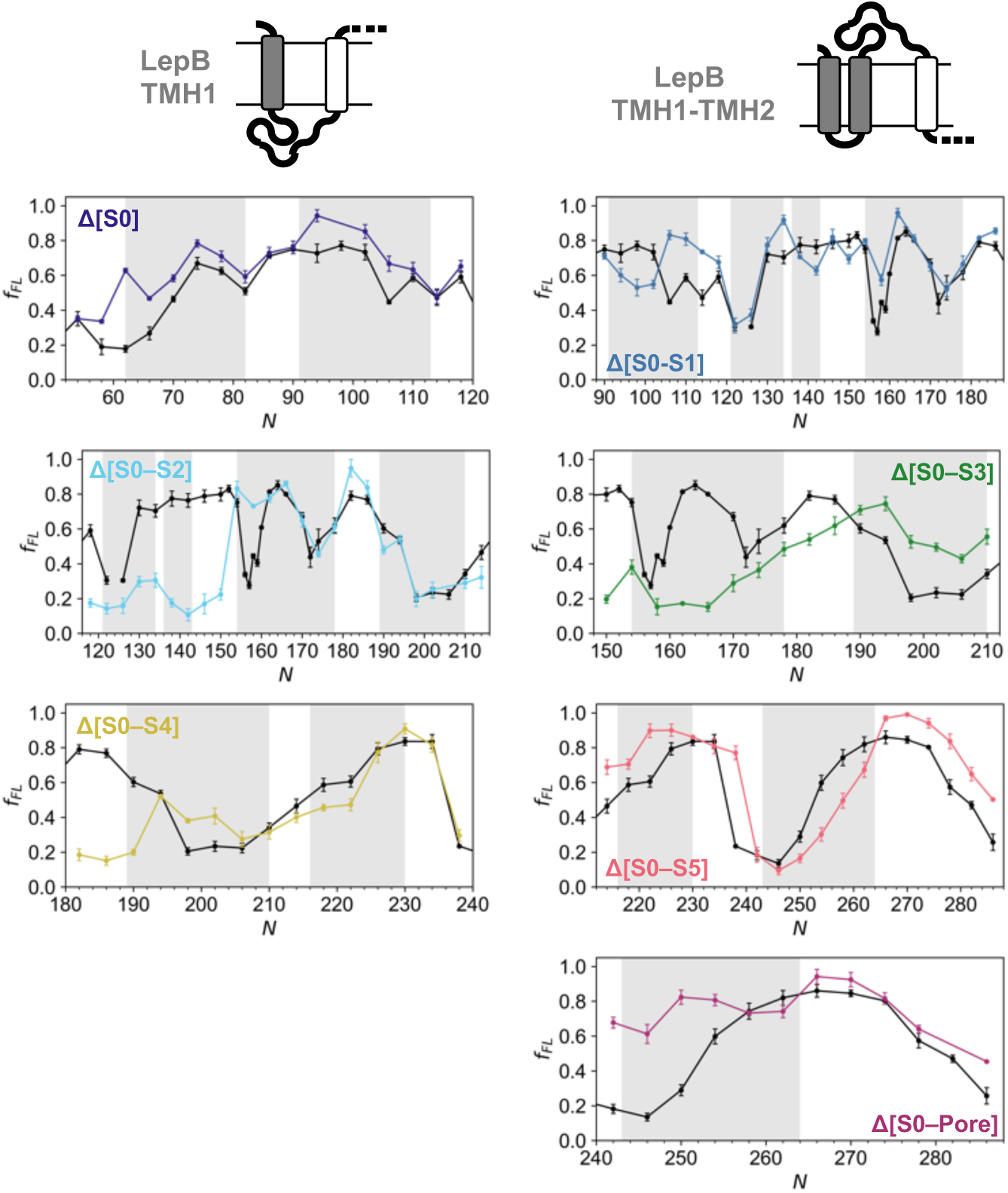
Same as in Fig. 3 (main text), but with each deletion FP plotted separately. The relevant portions of the full KvAP FP (Fig. 2) are in black.

**Supplemental Figure S4.**
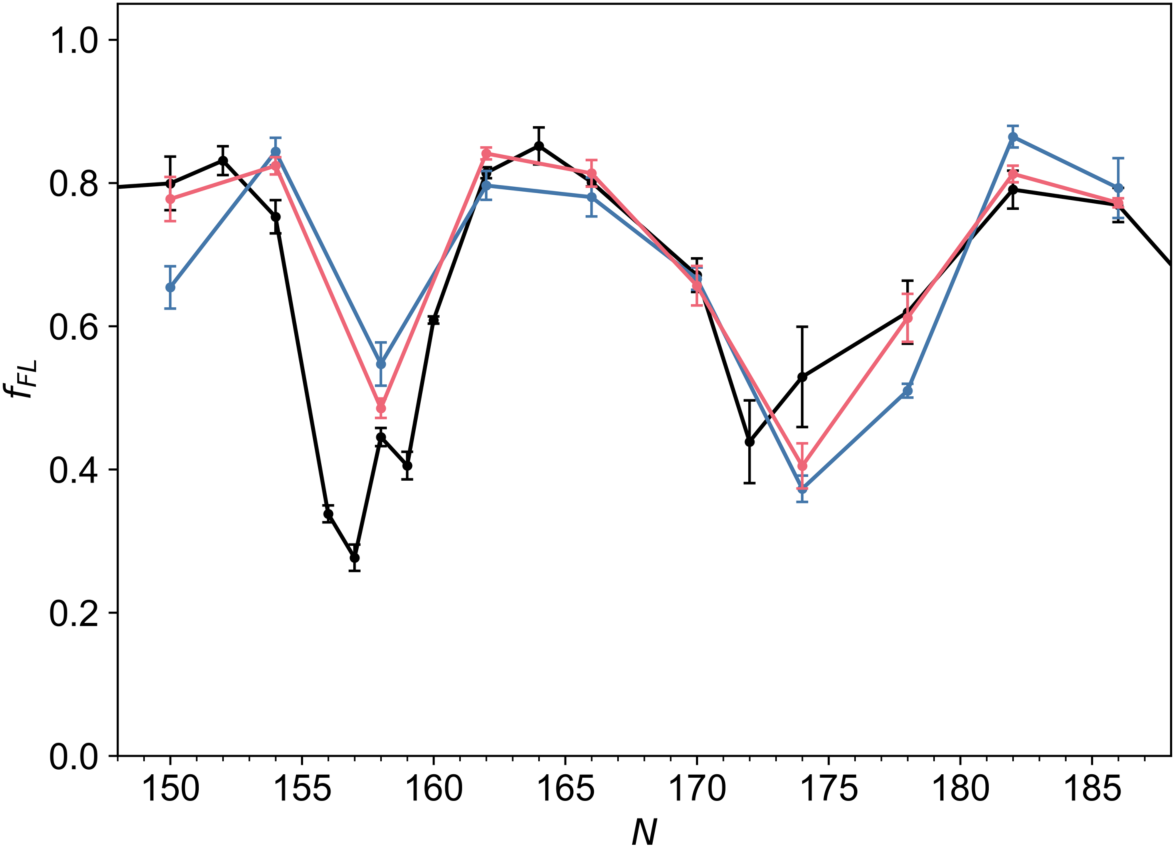
Same as in Fig. 4 (main text), but for mutations E^45^A and E^93^A.

**Table S1.**
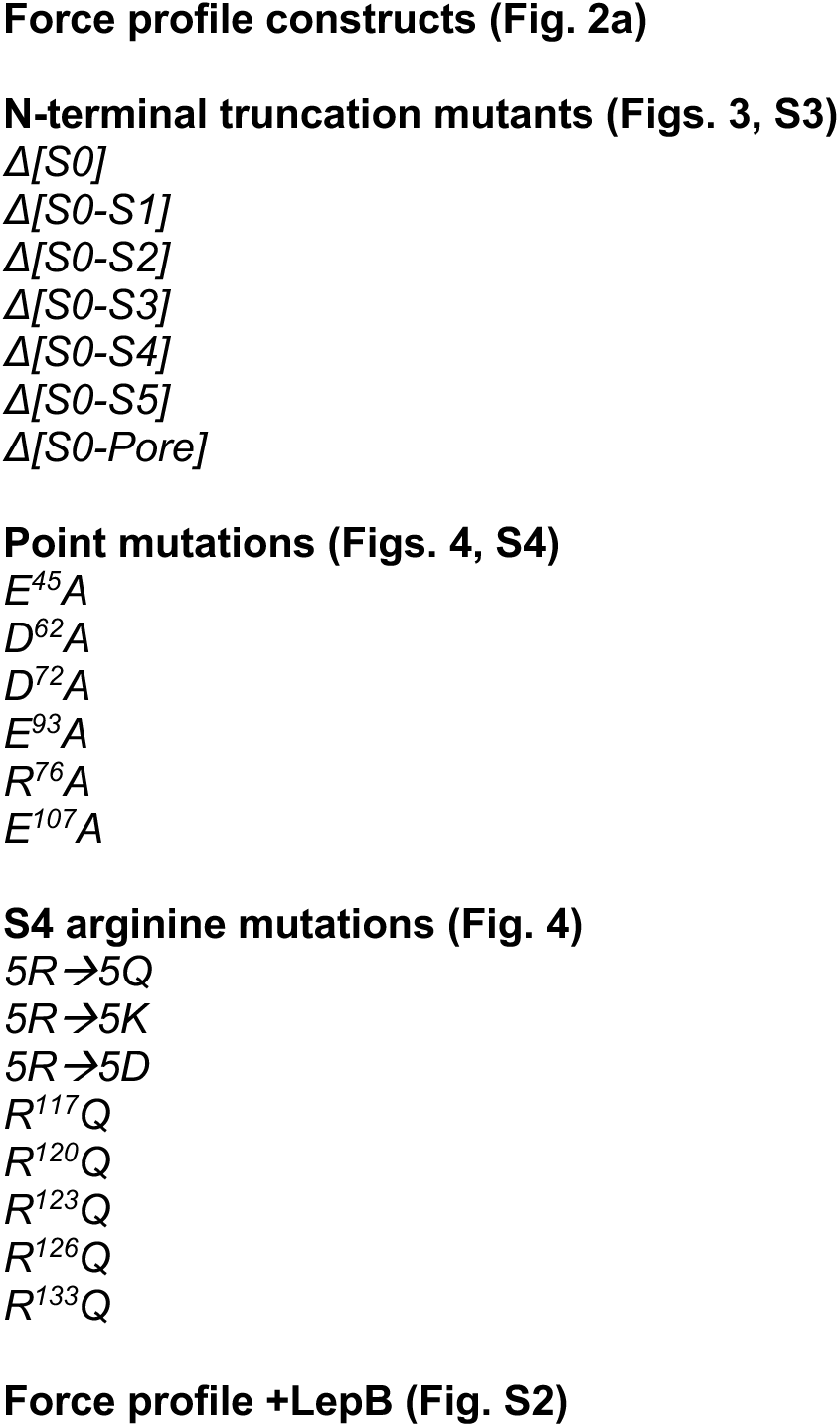
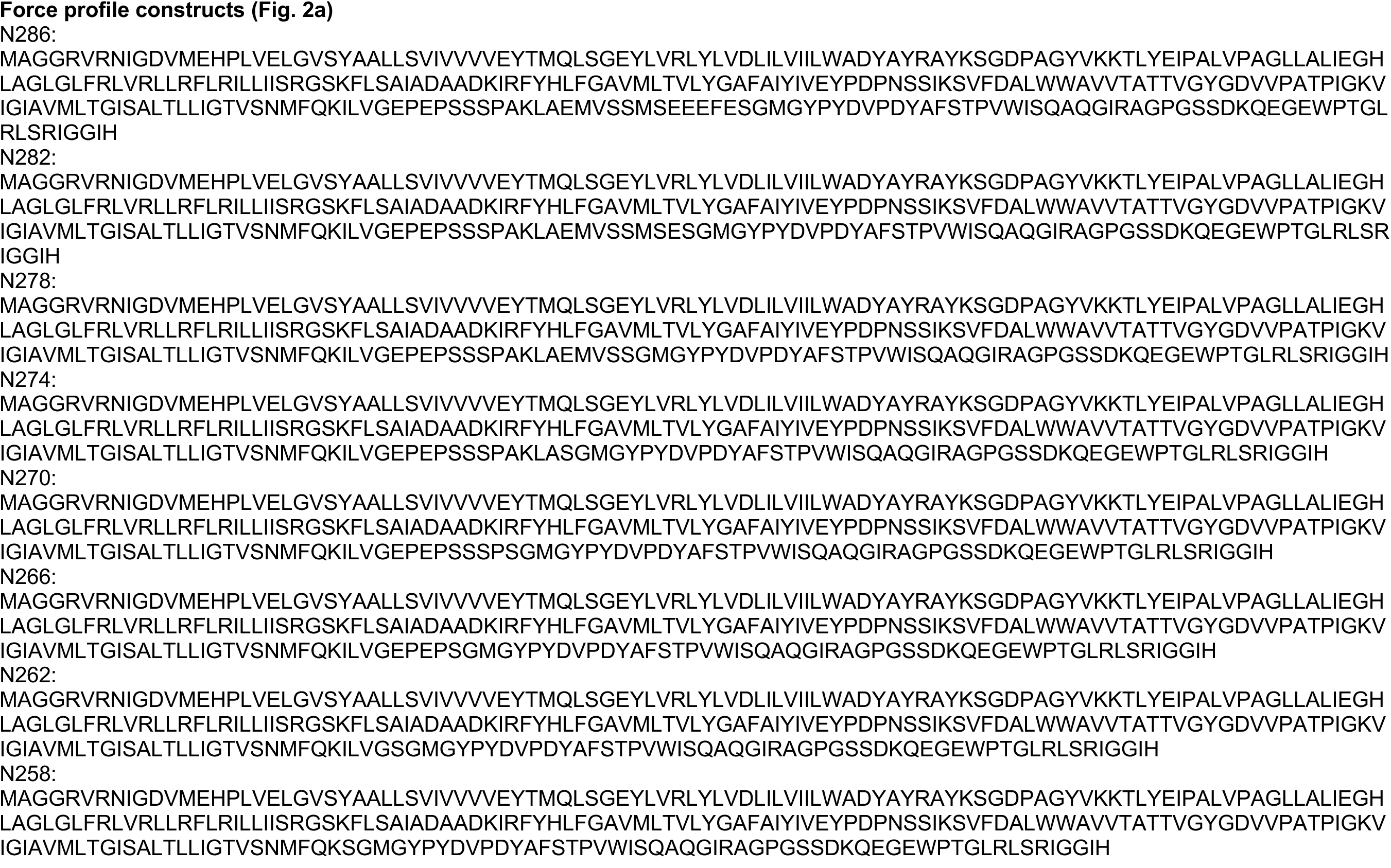
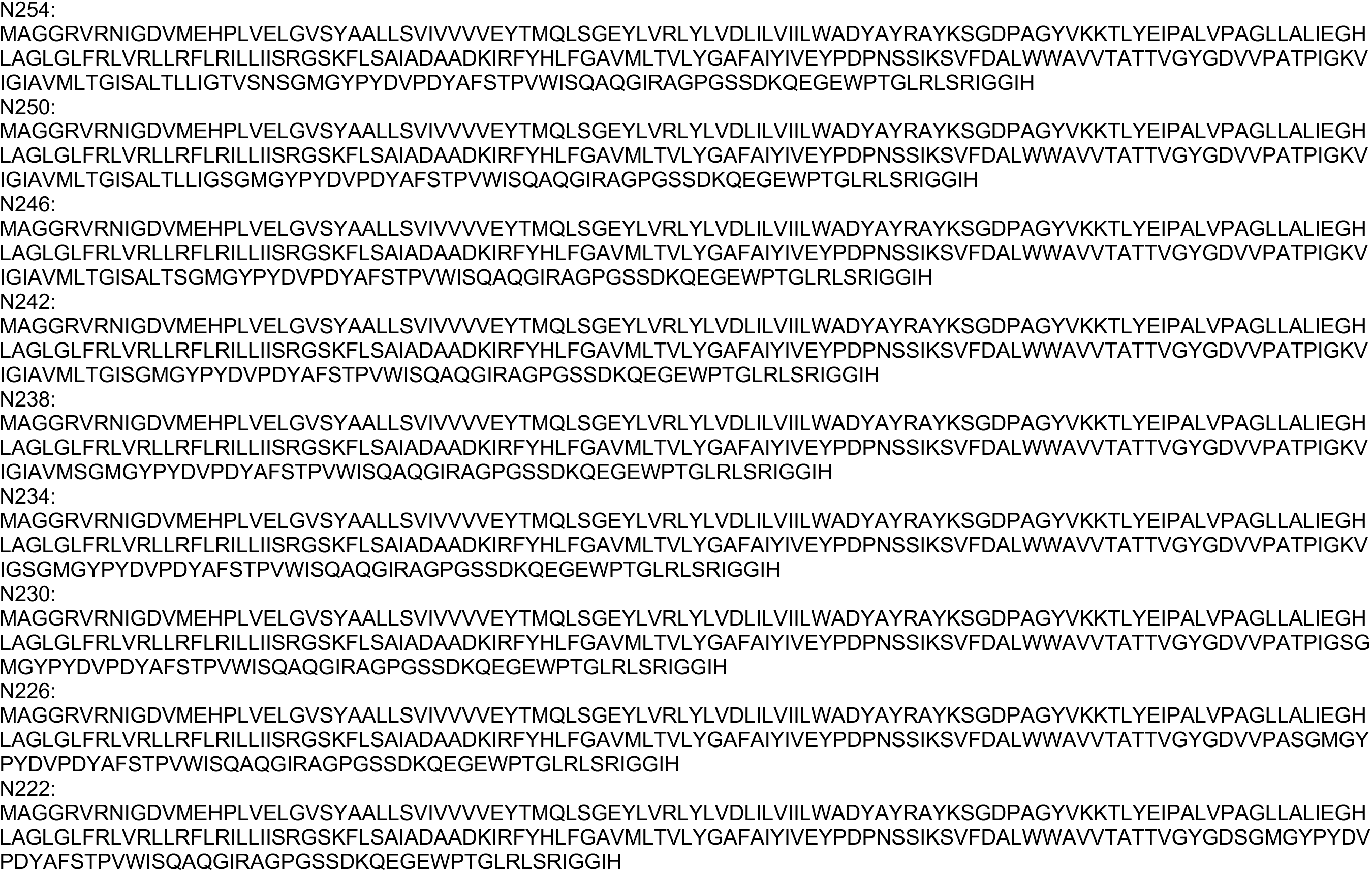
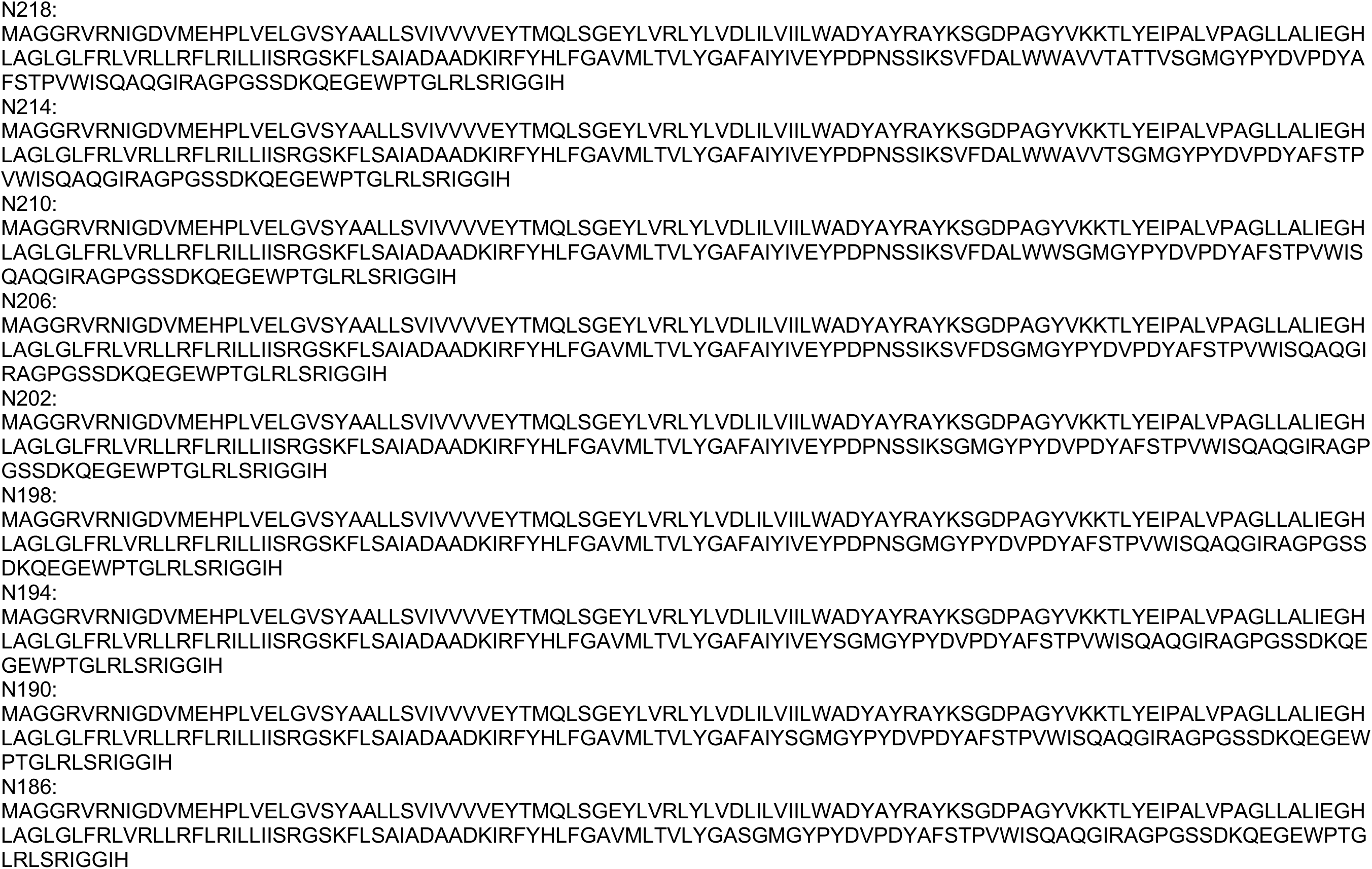
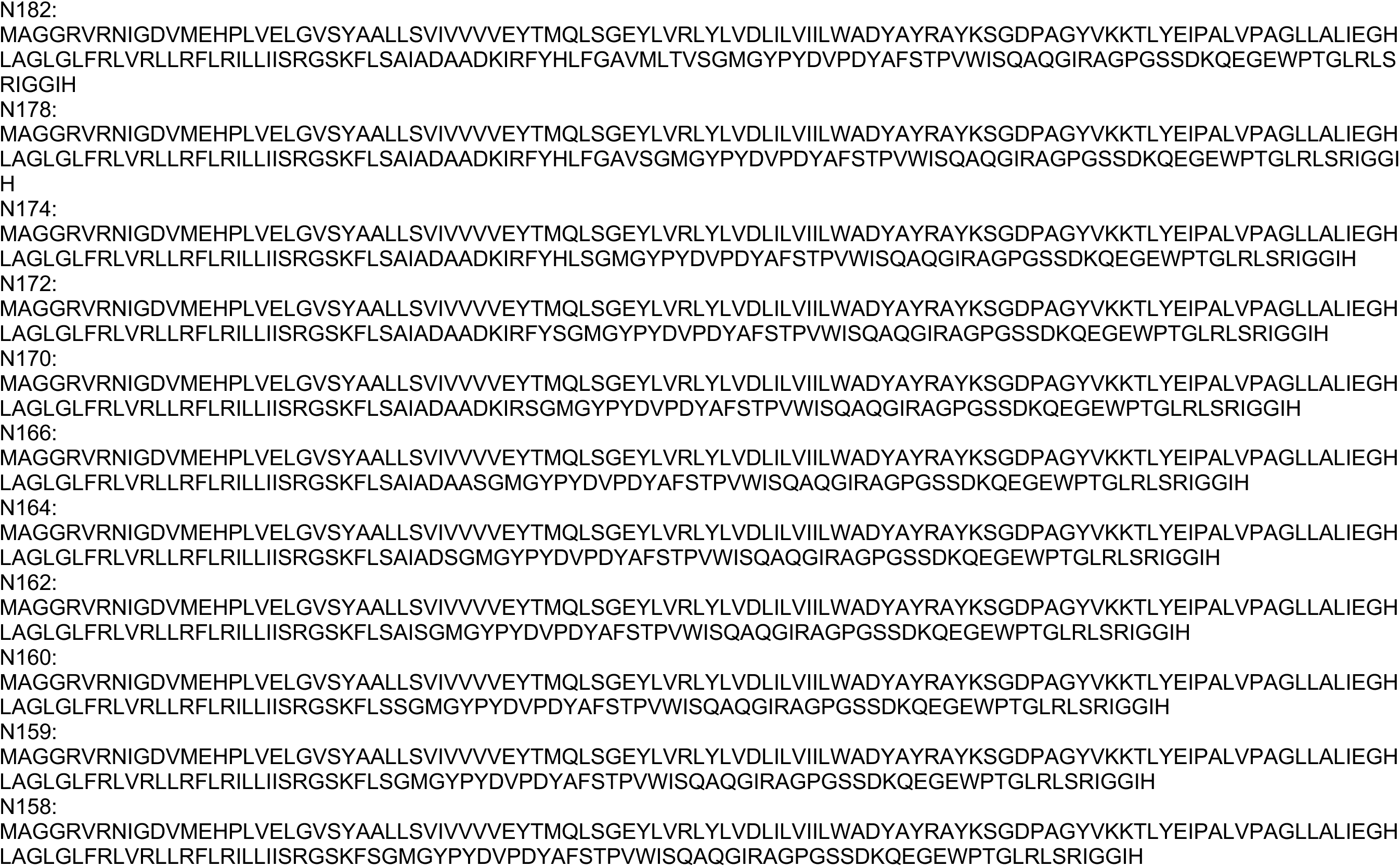
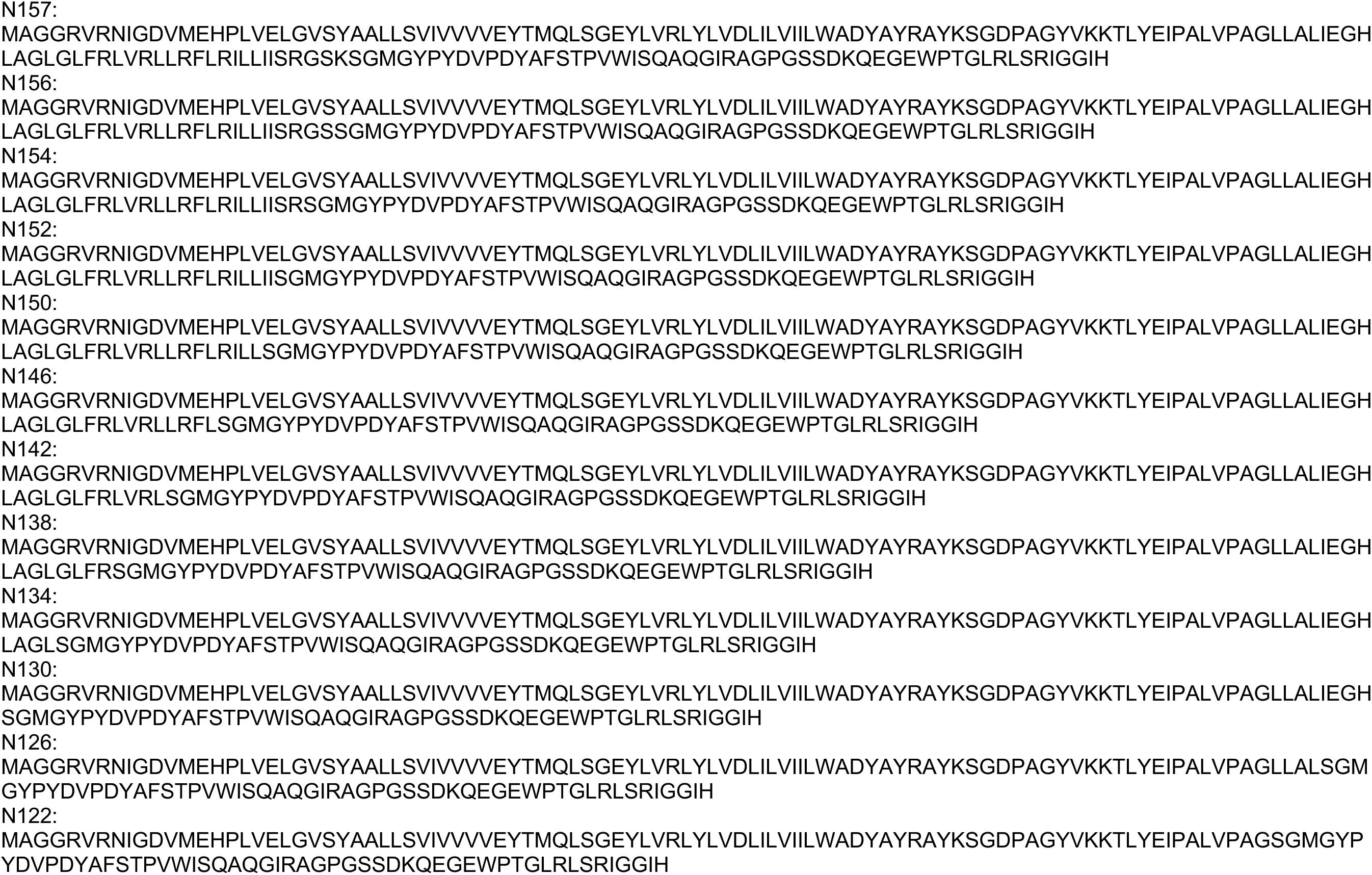
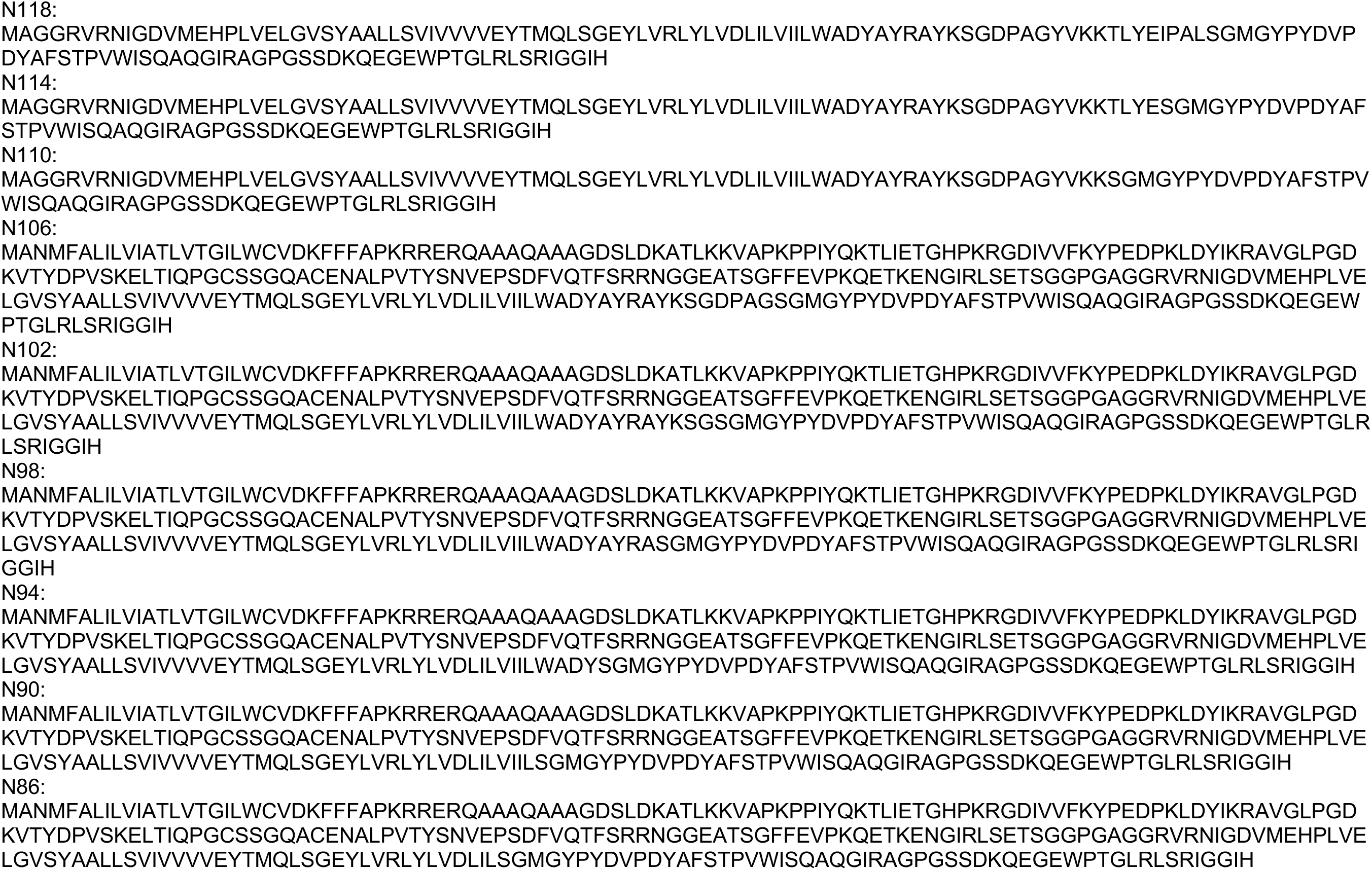
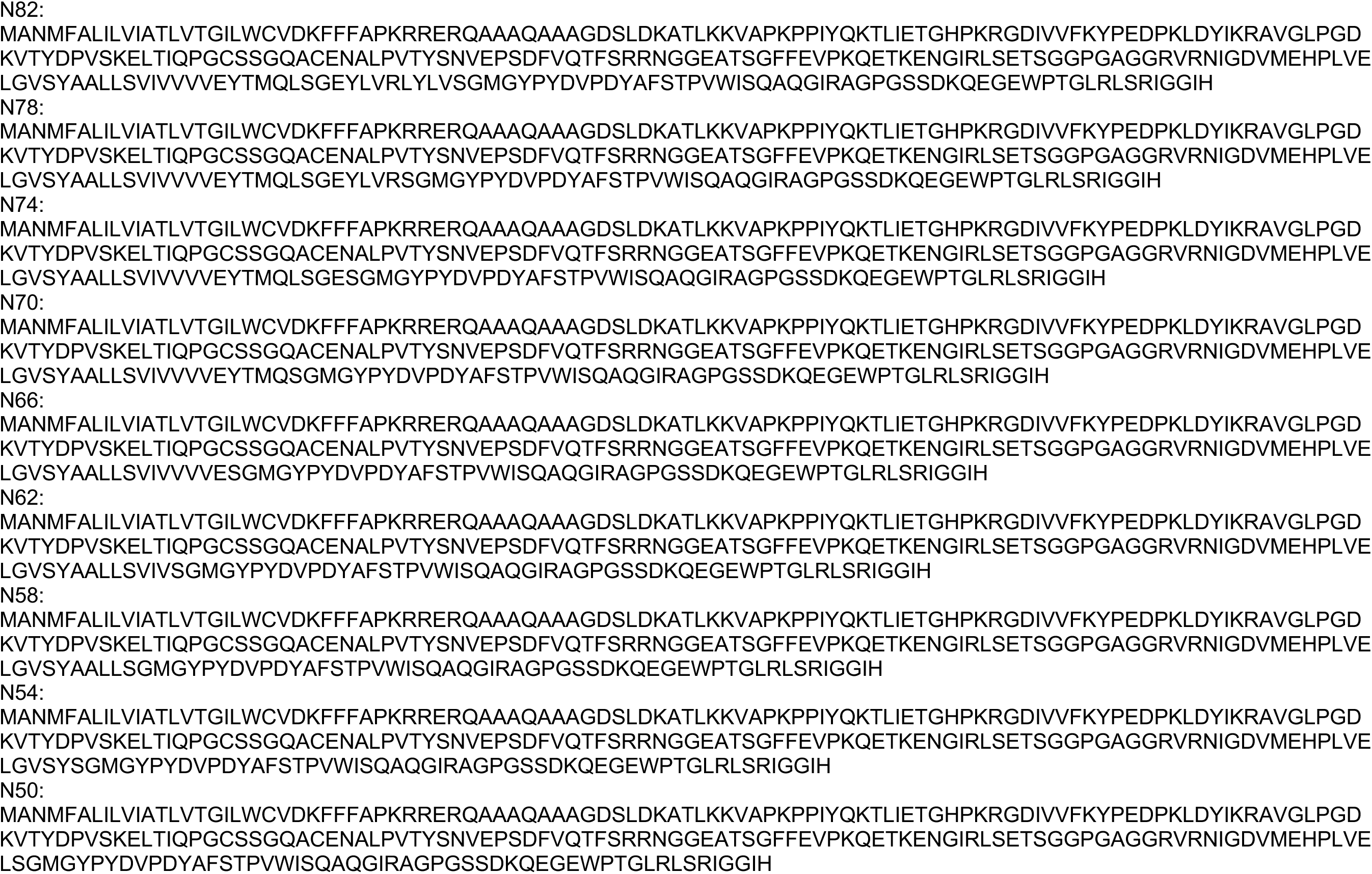
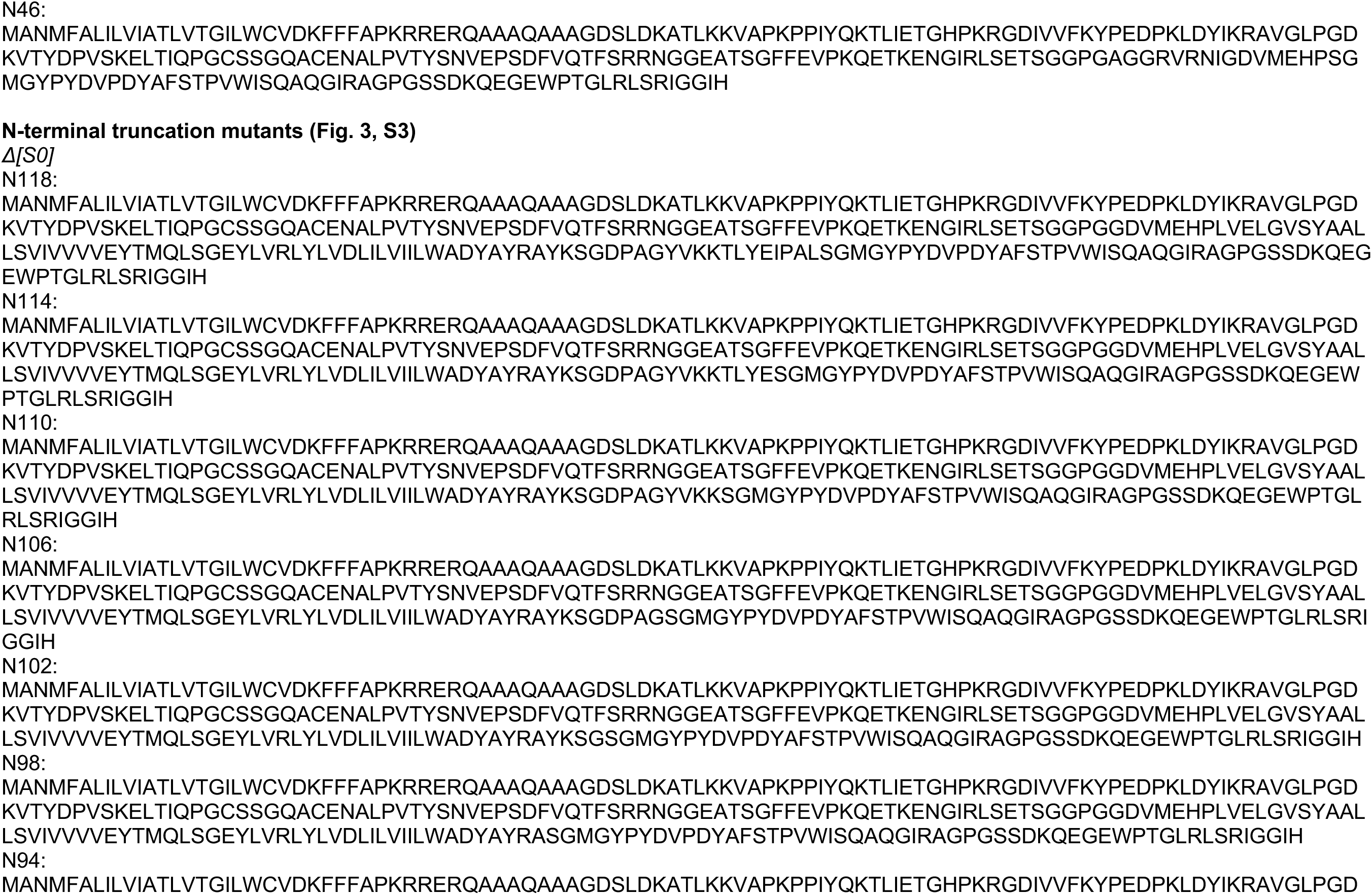
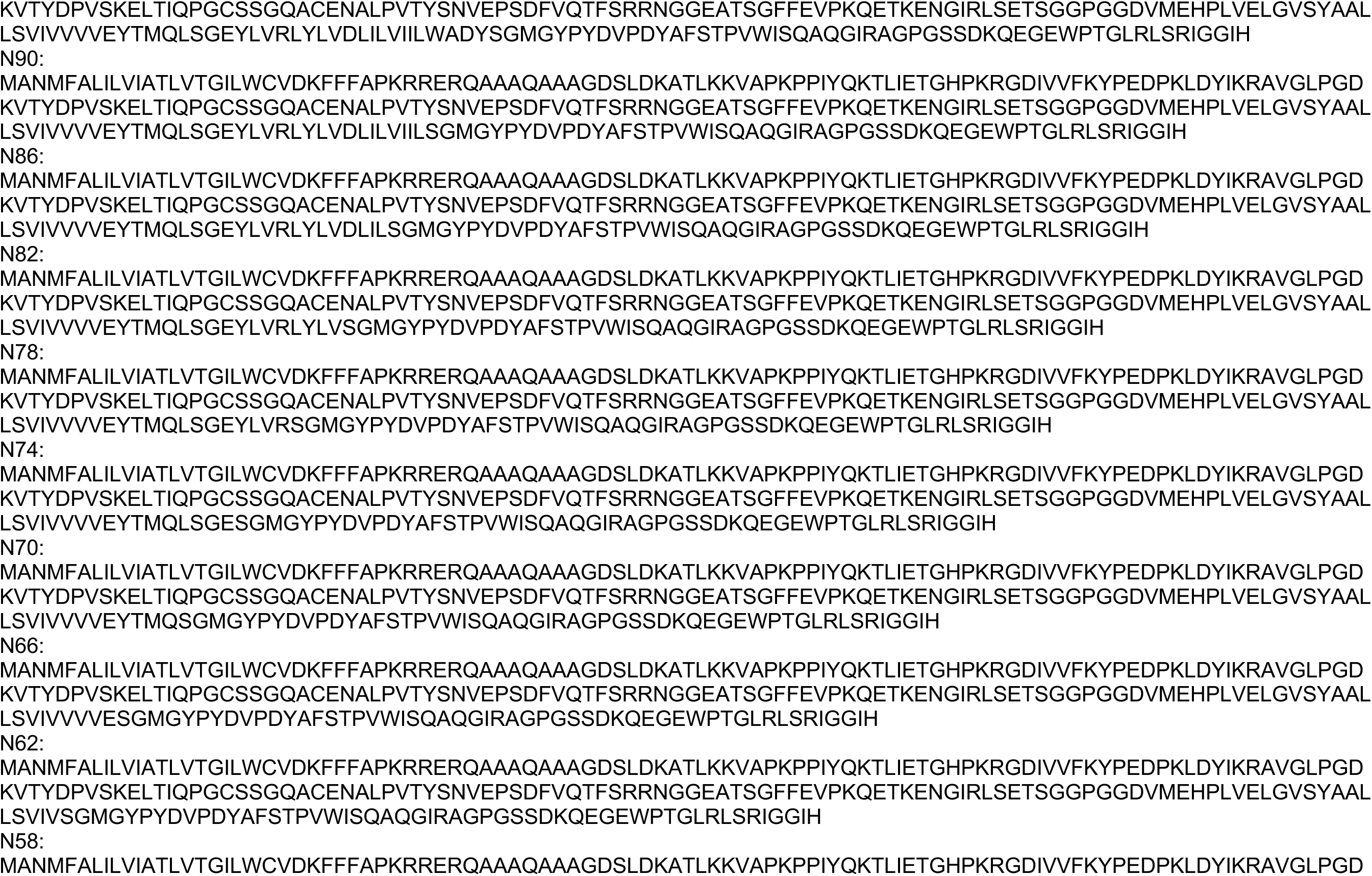
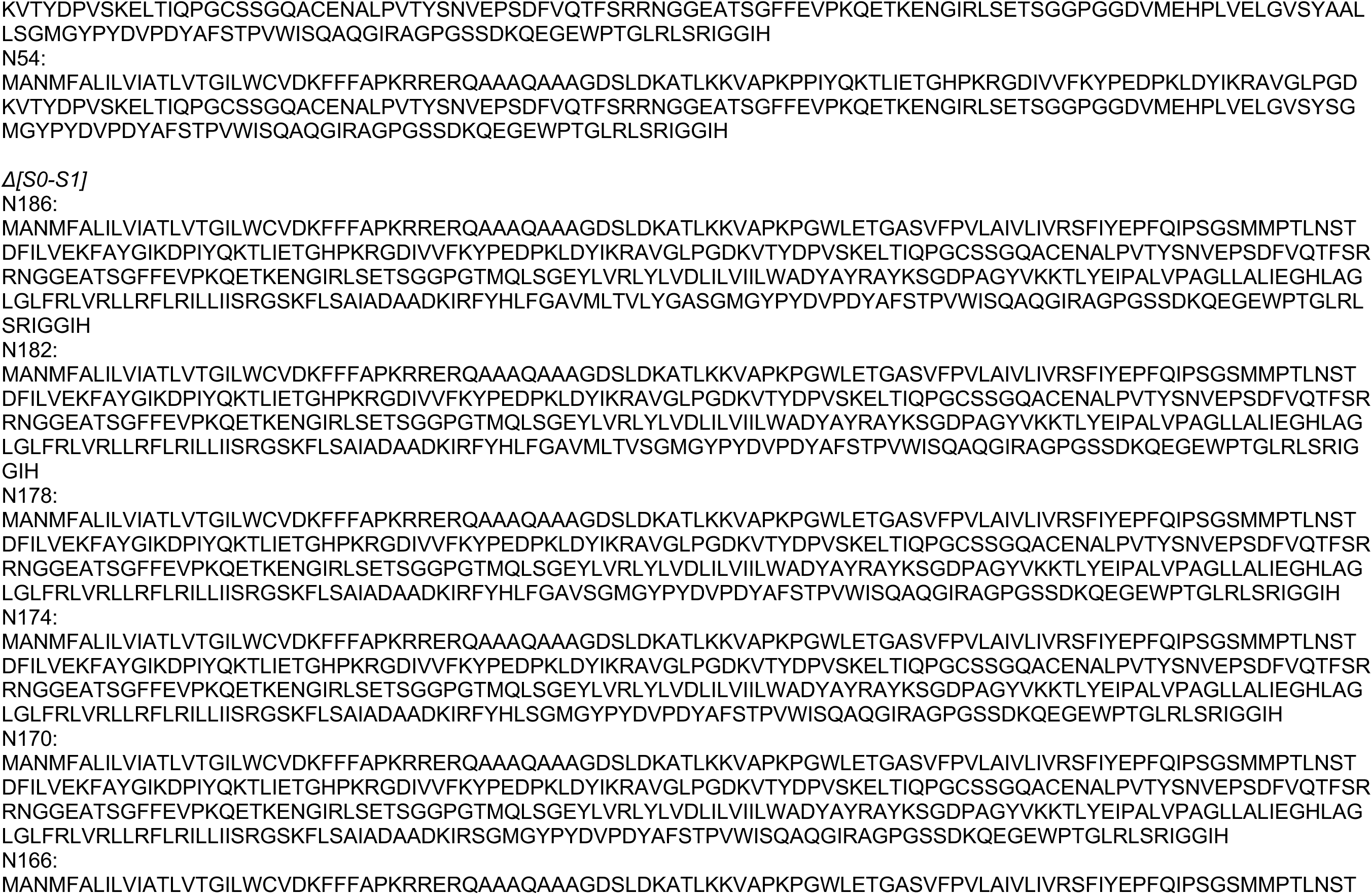
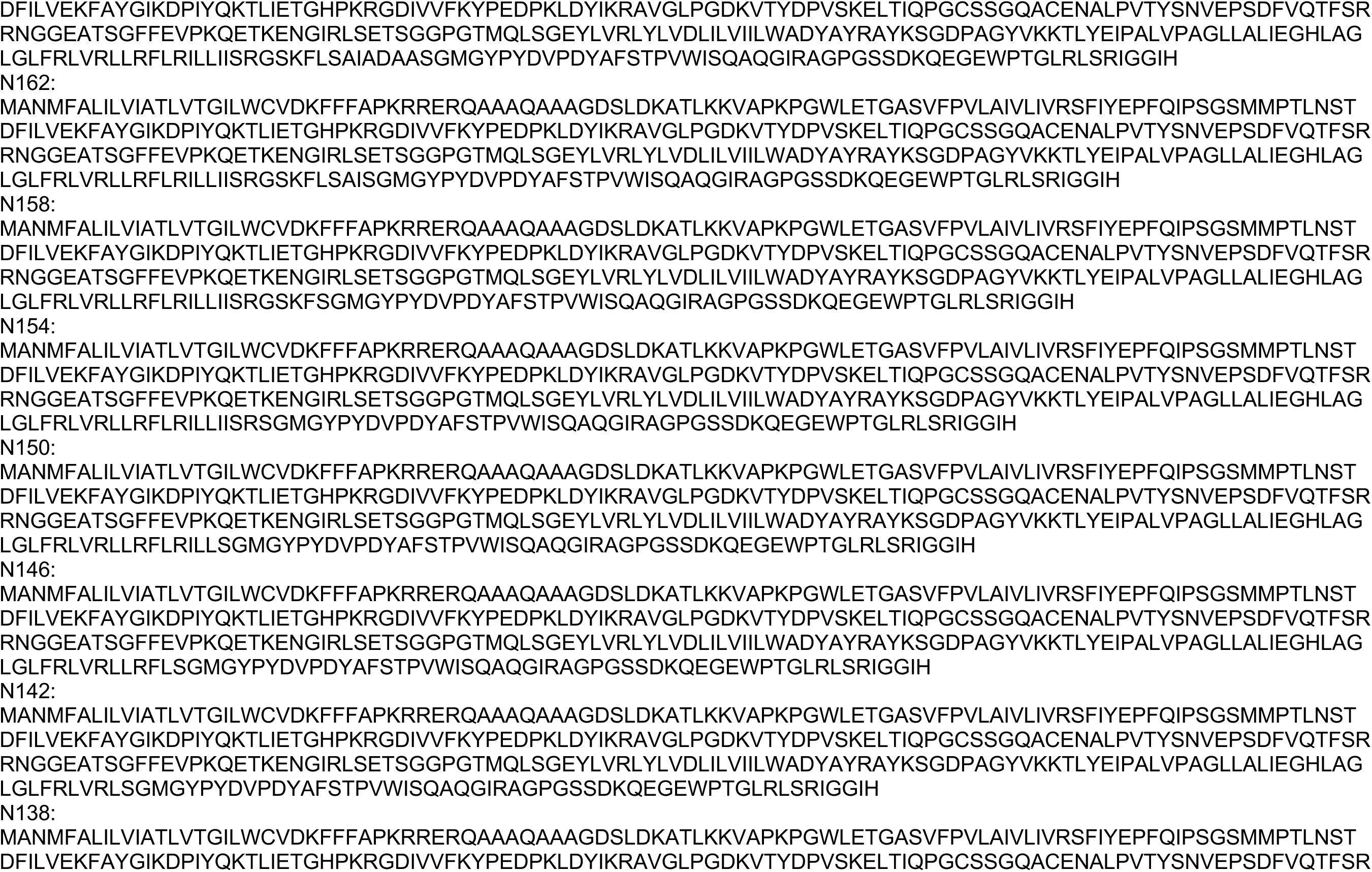
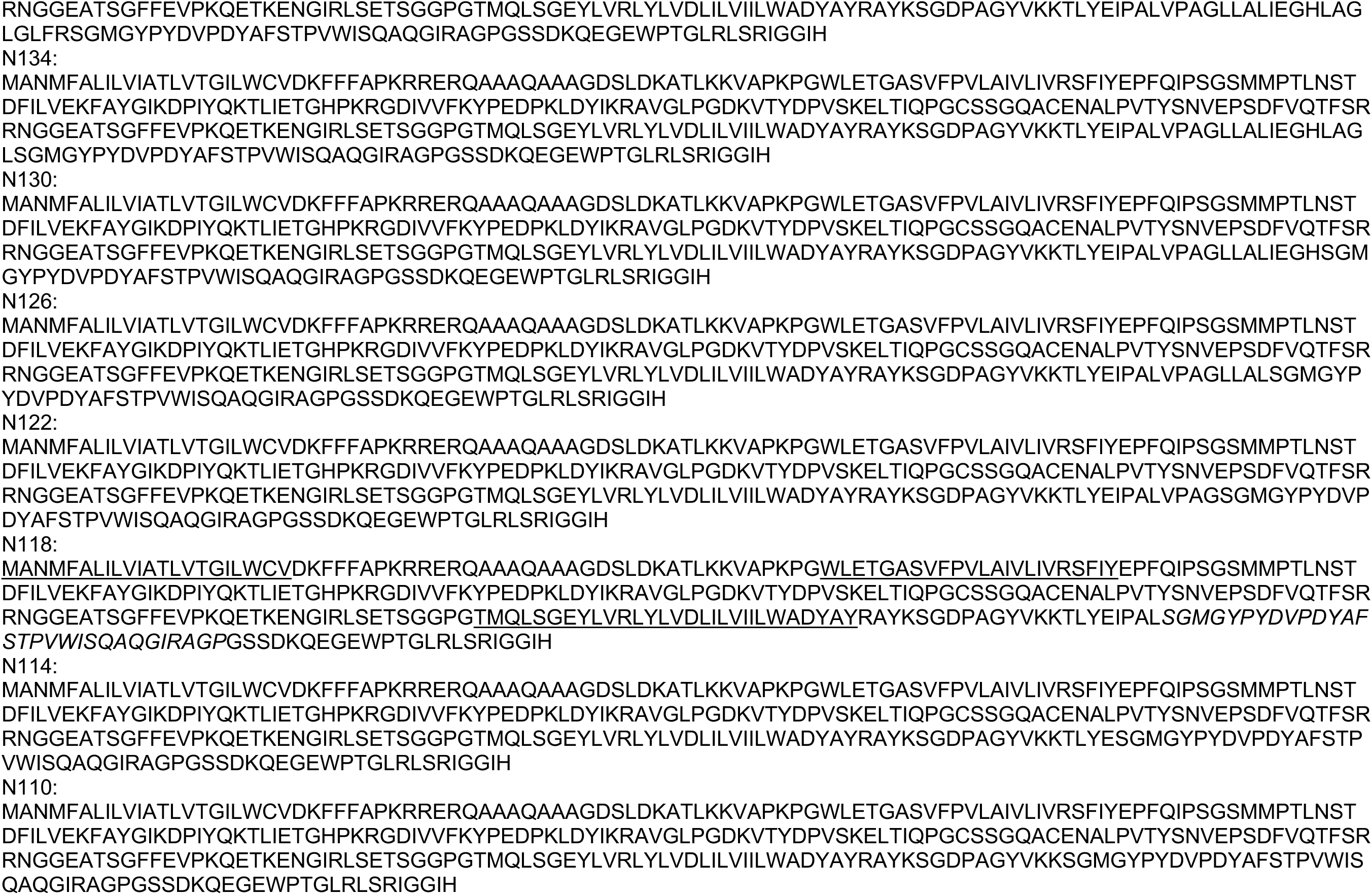
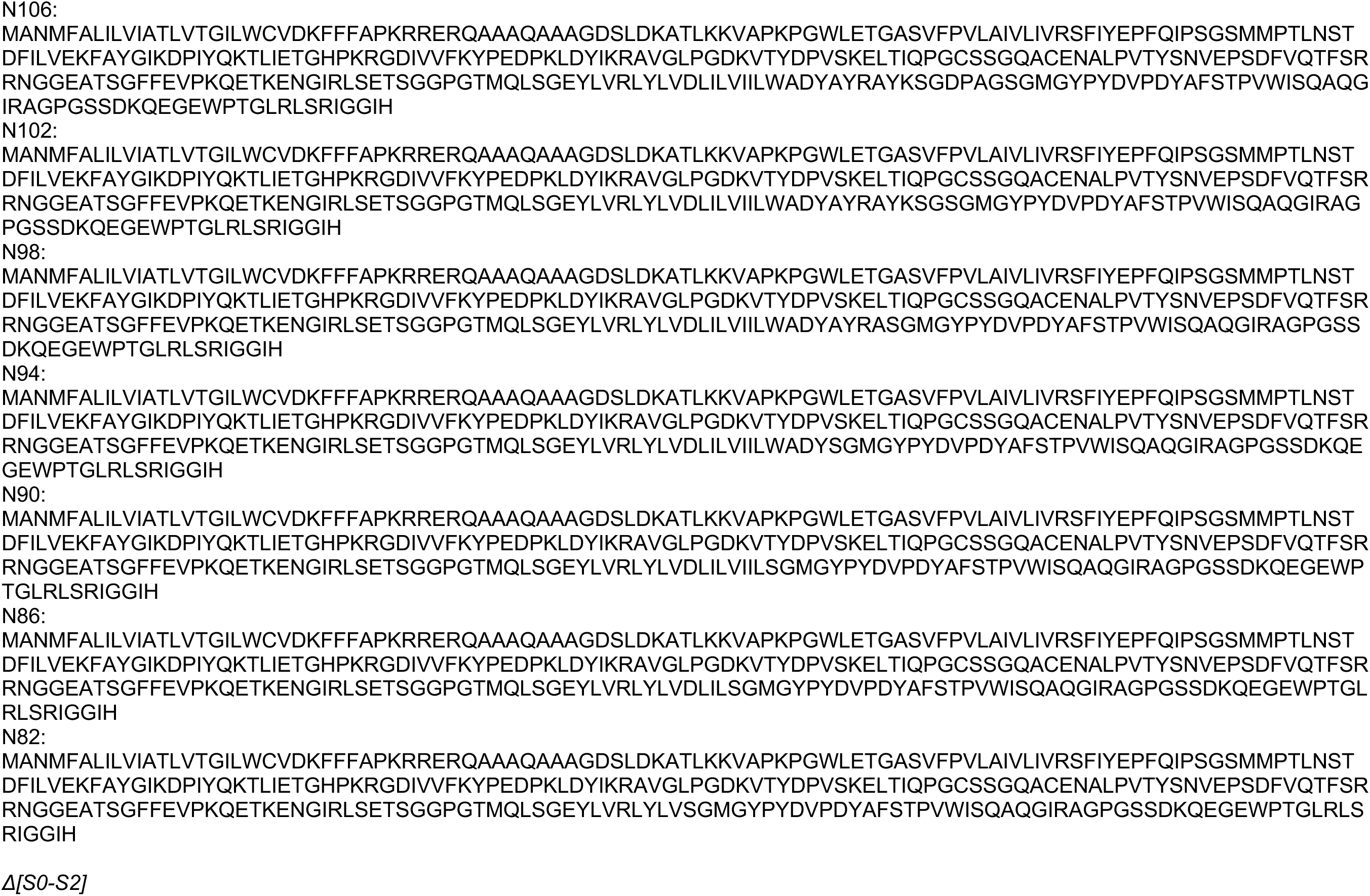
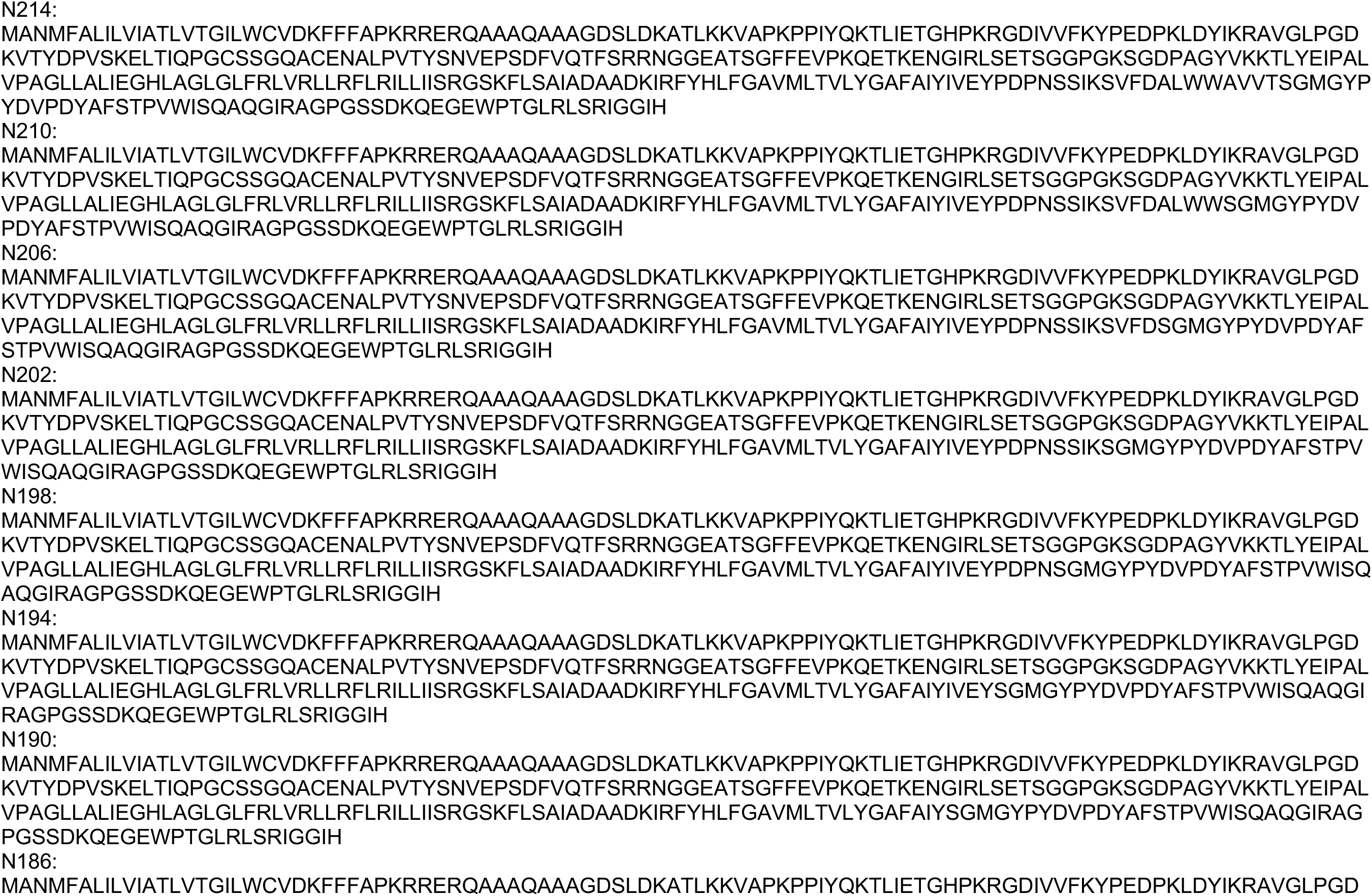
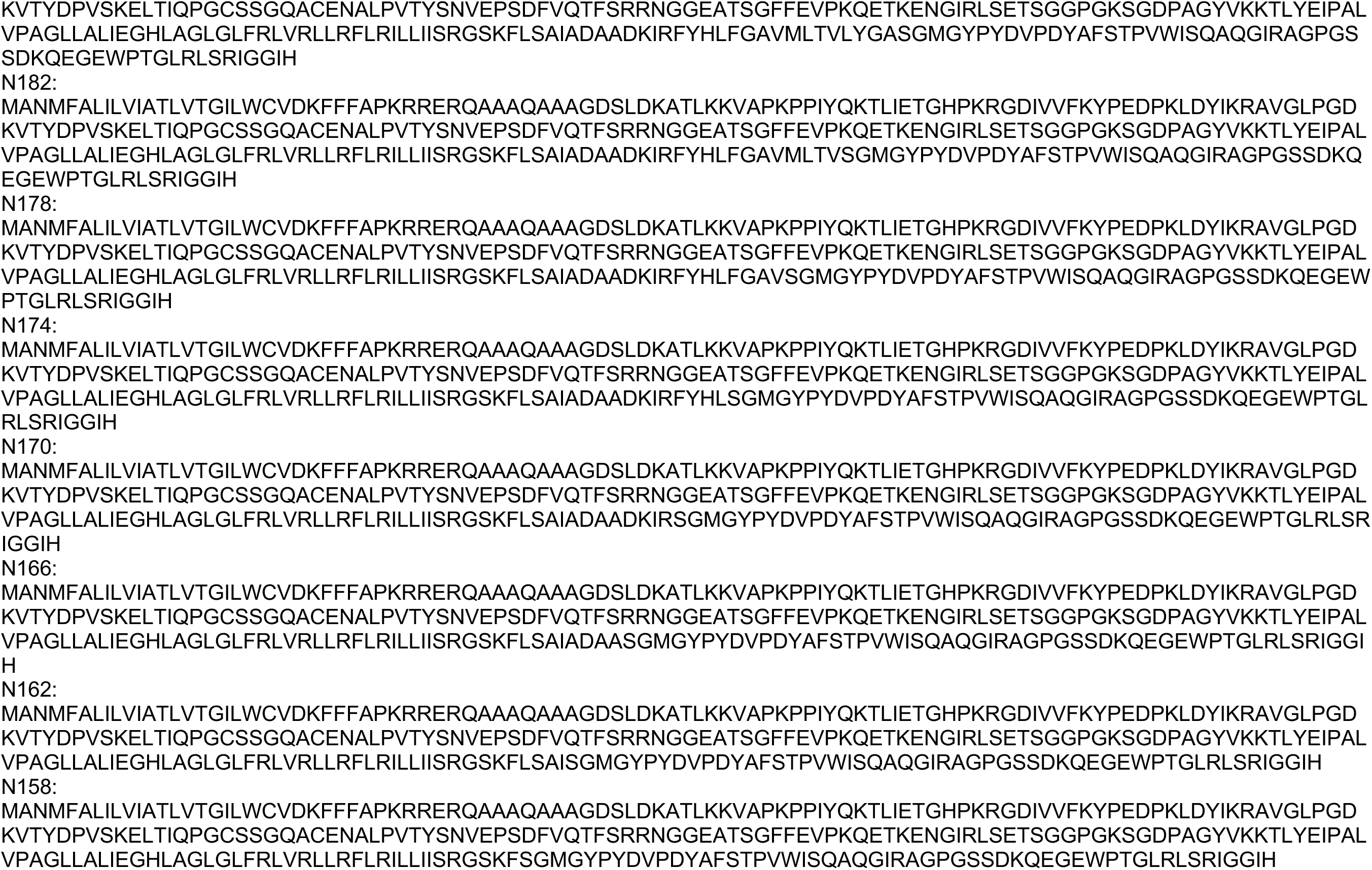
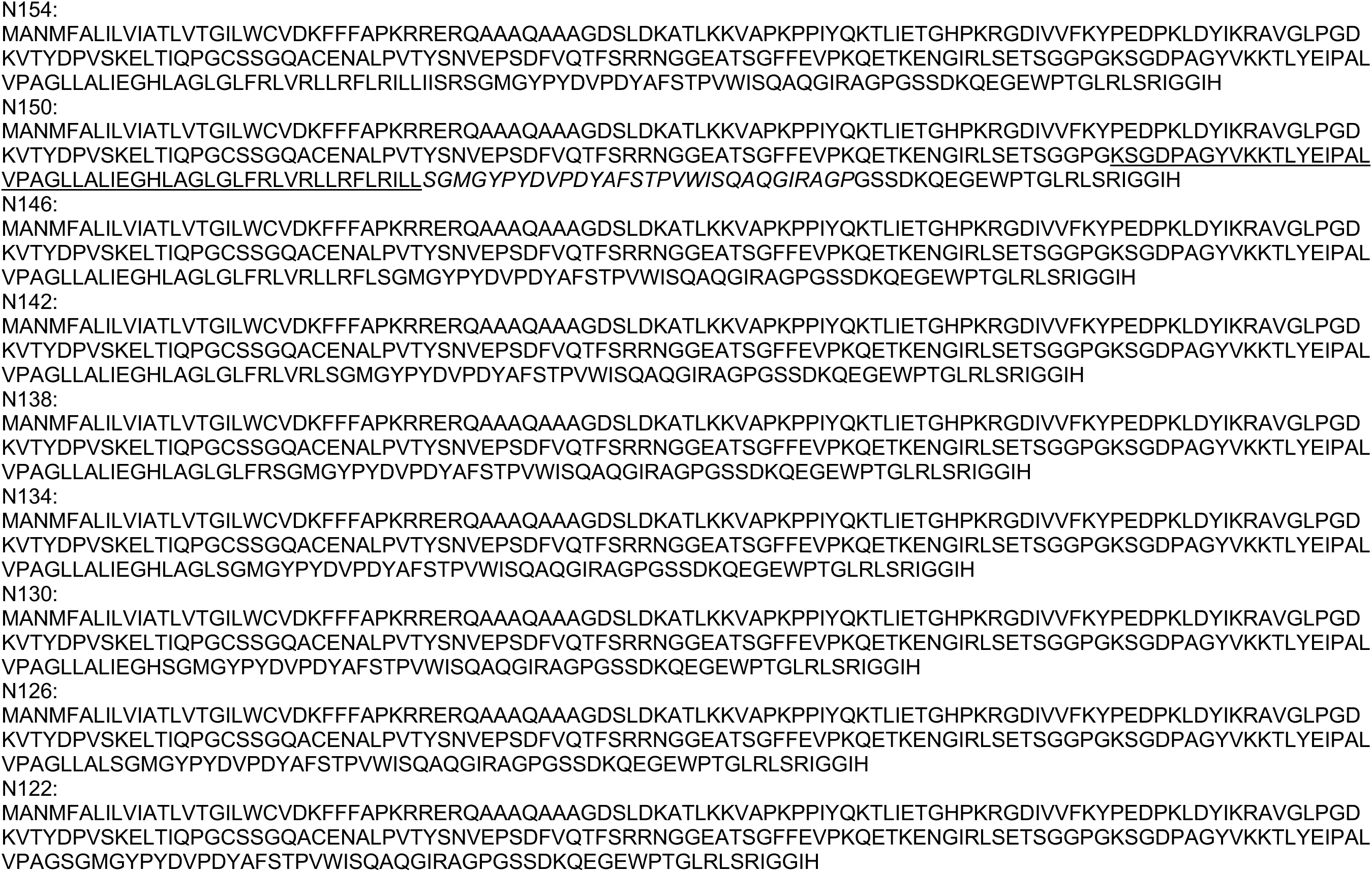
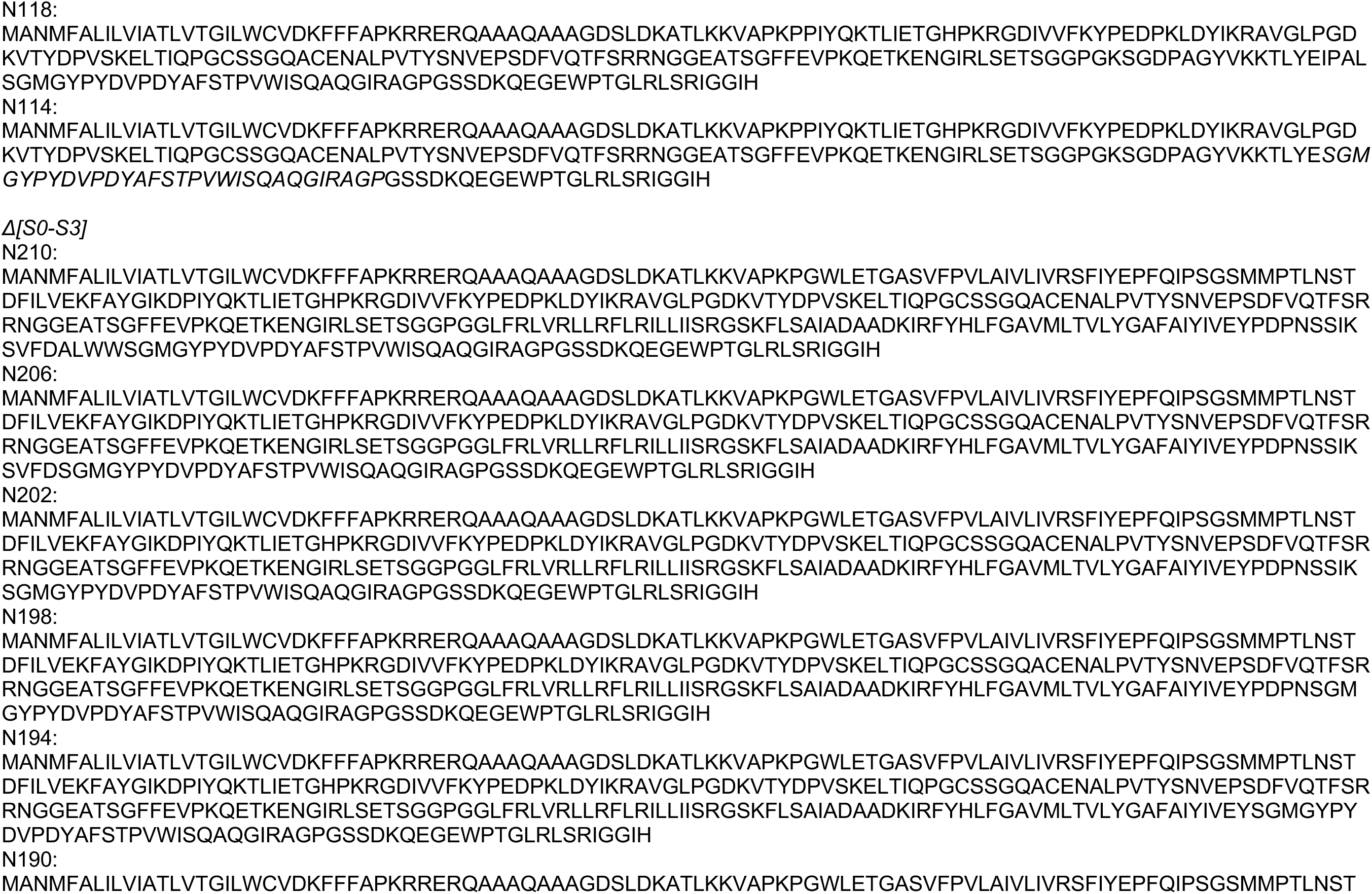
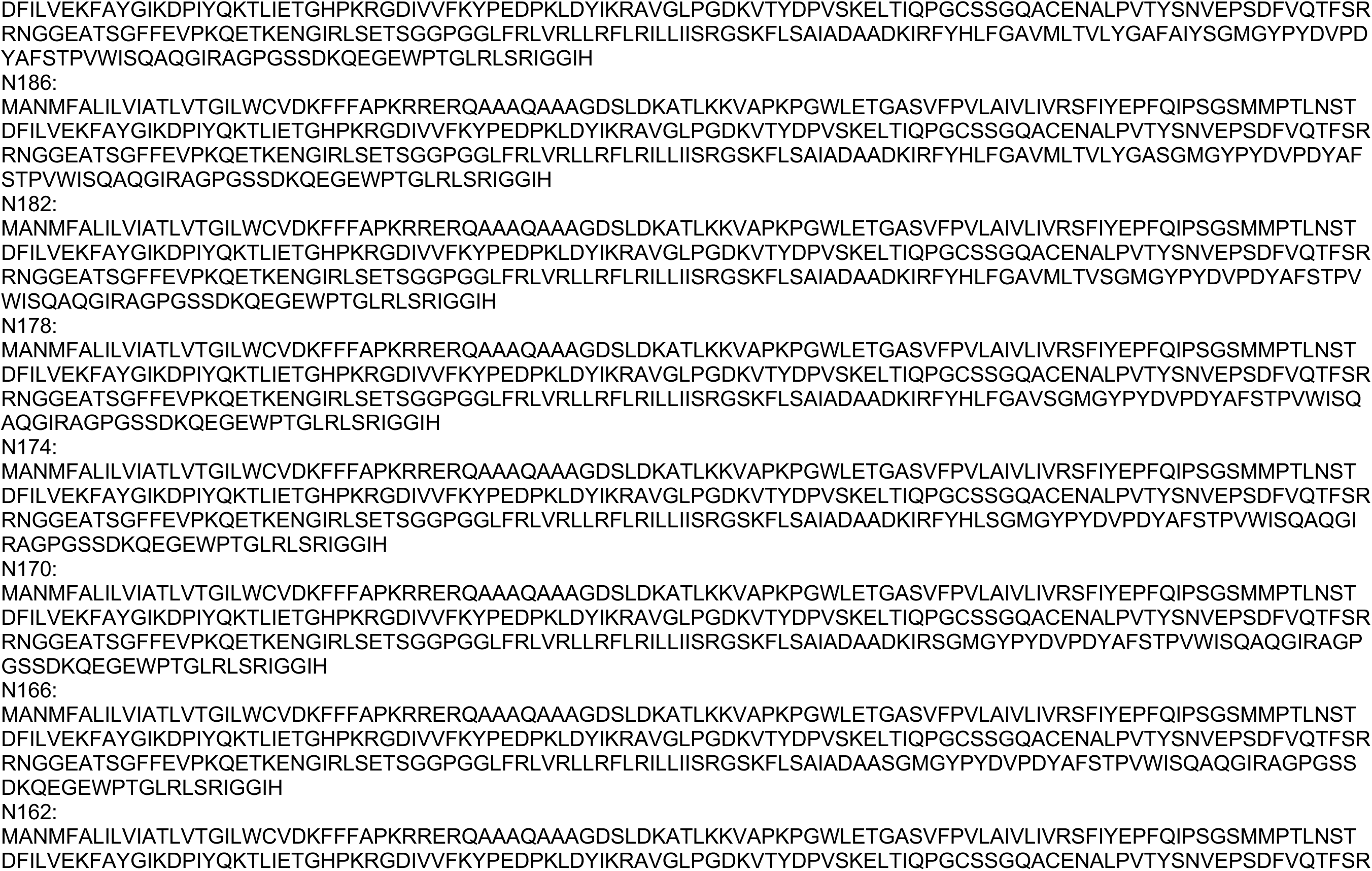
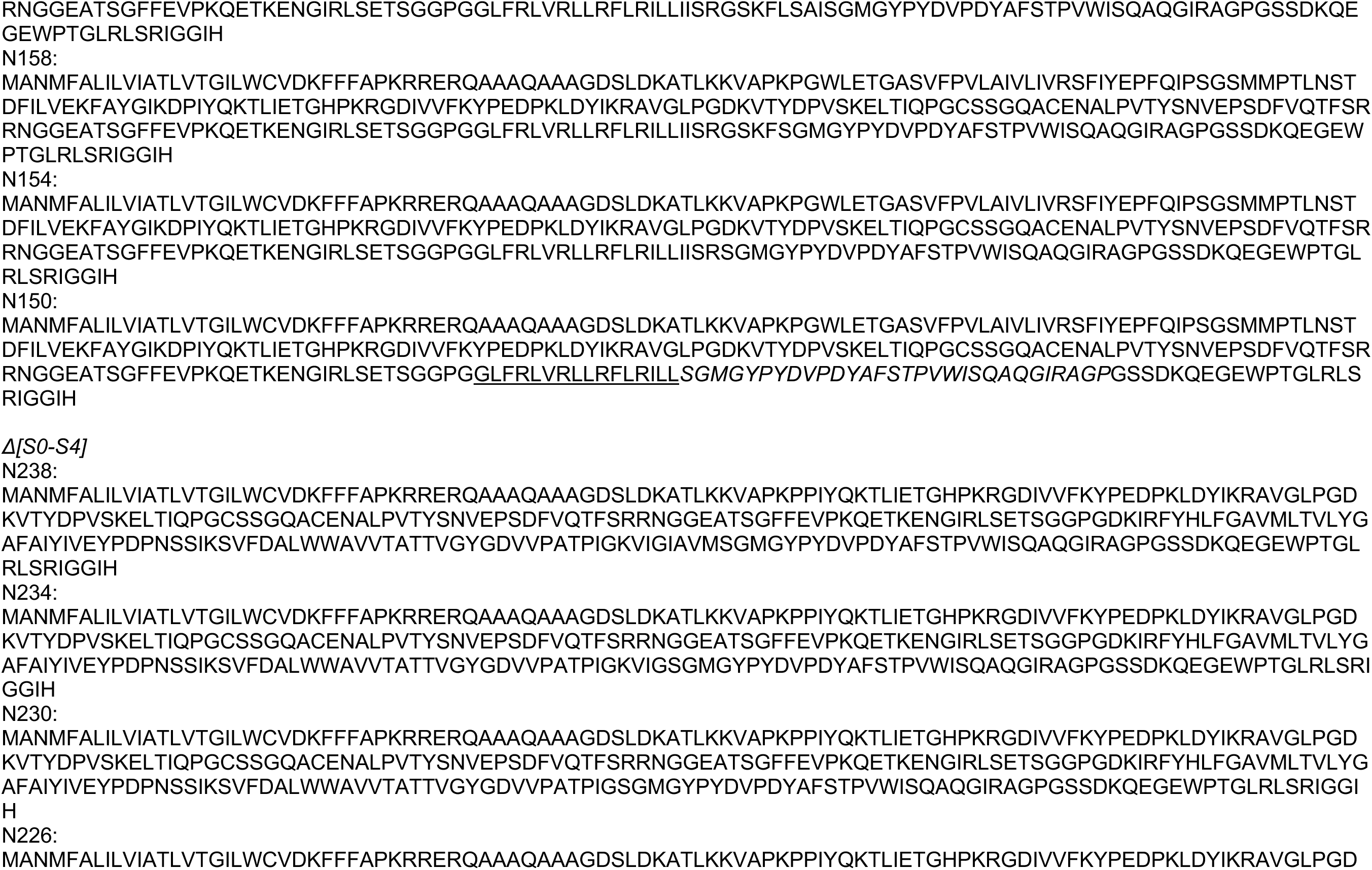
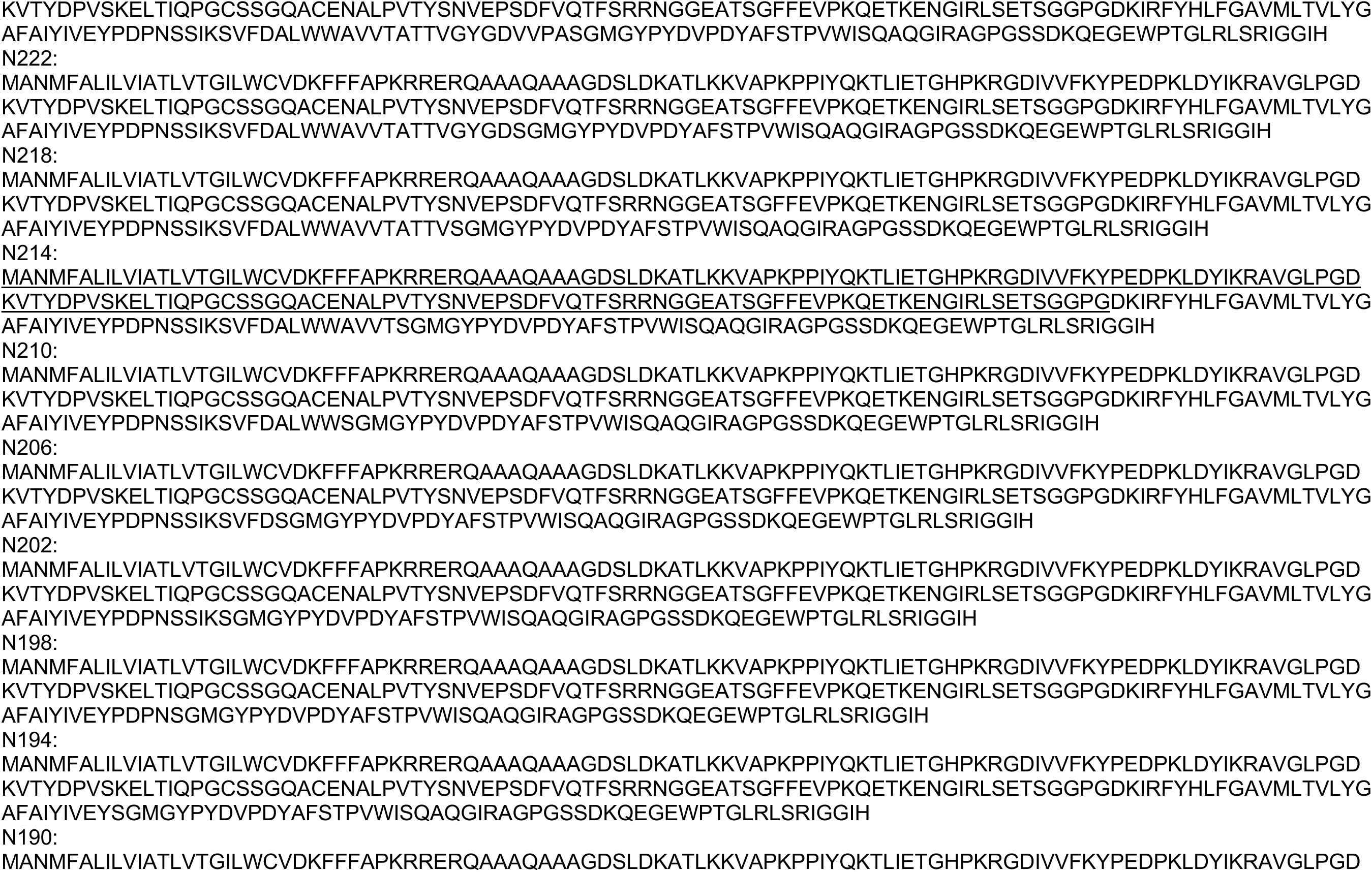
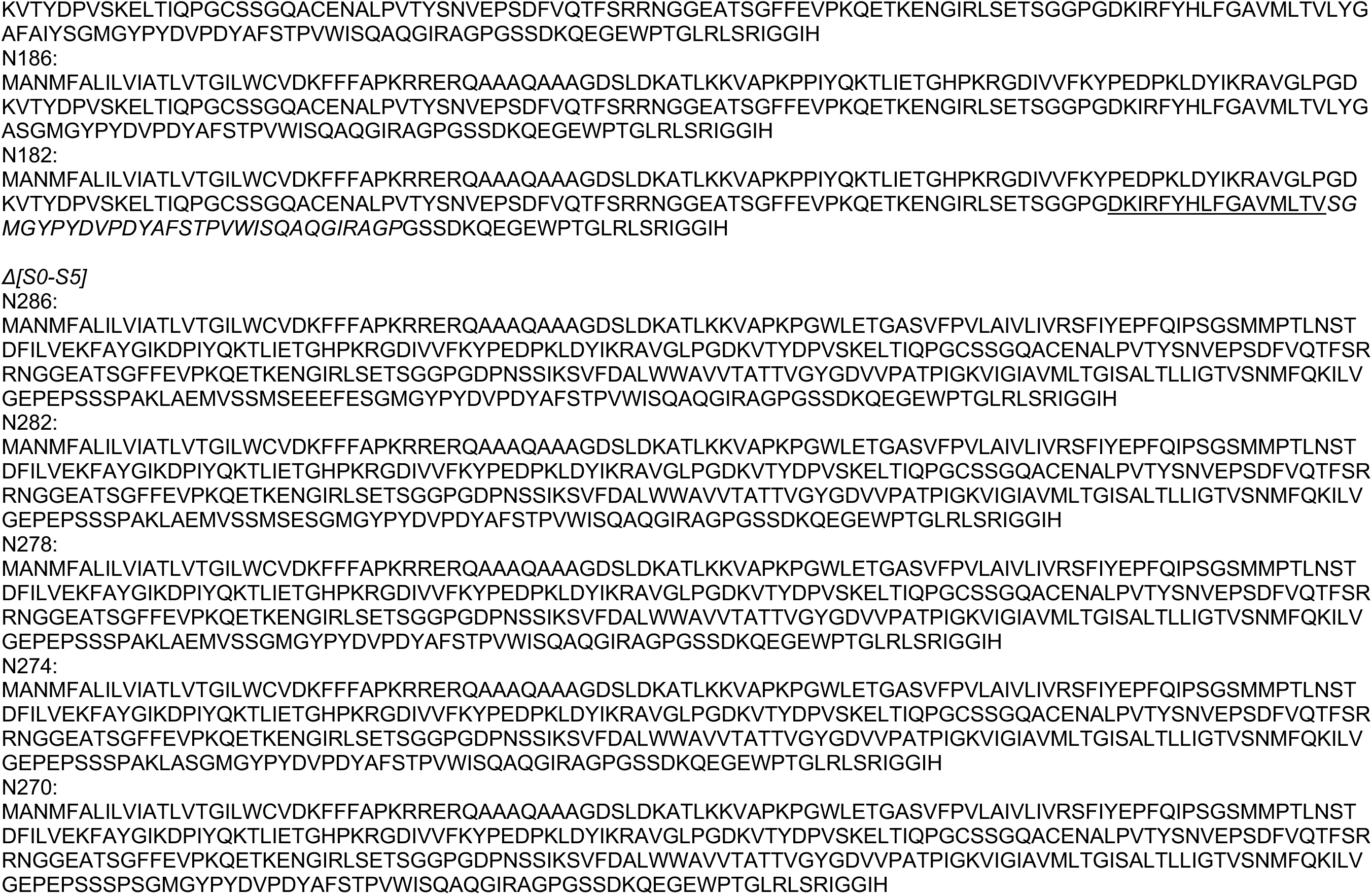
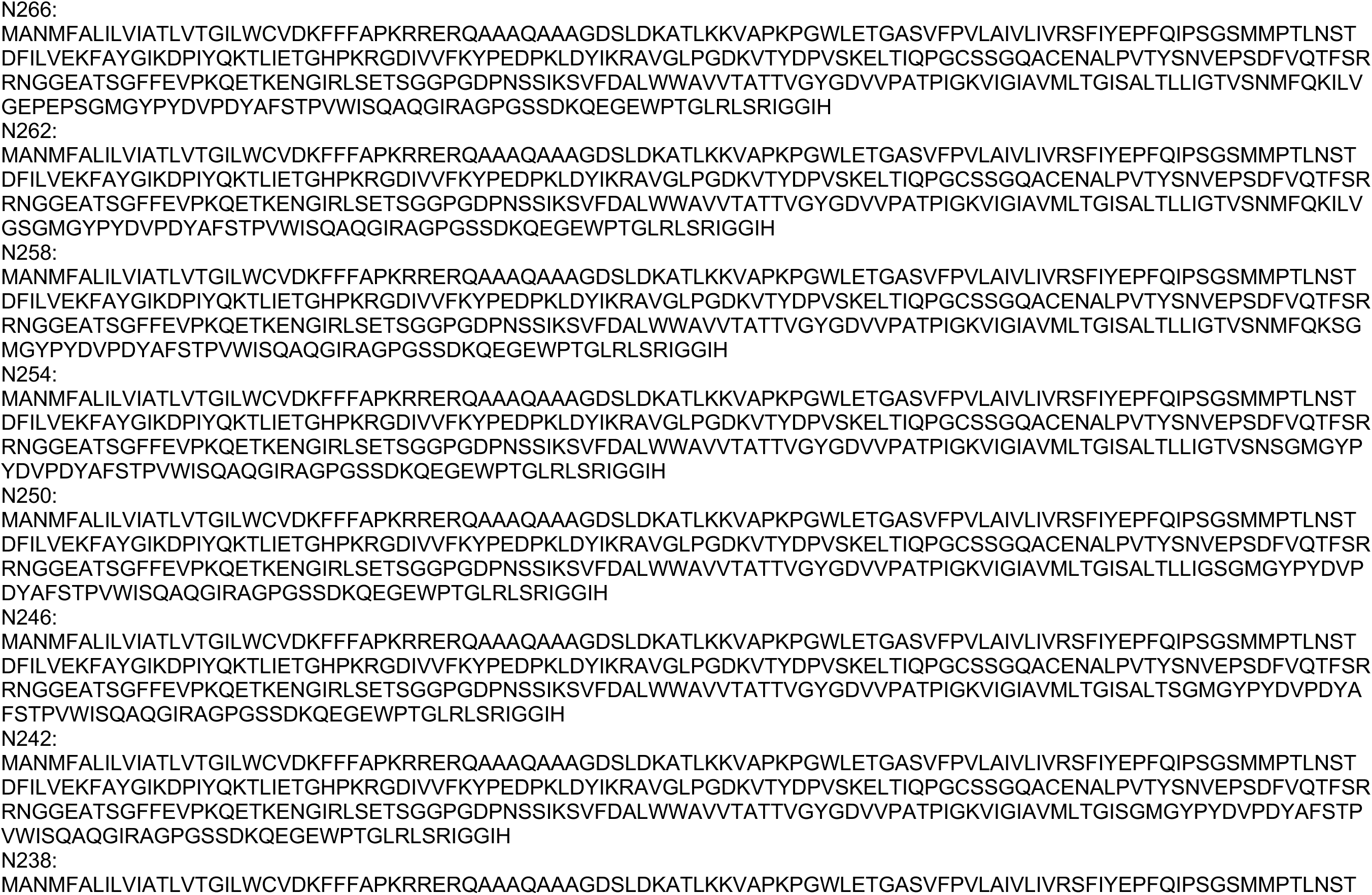
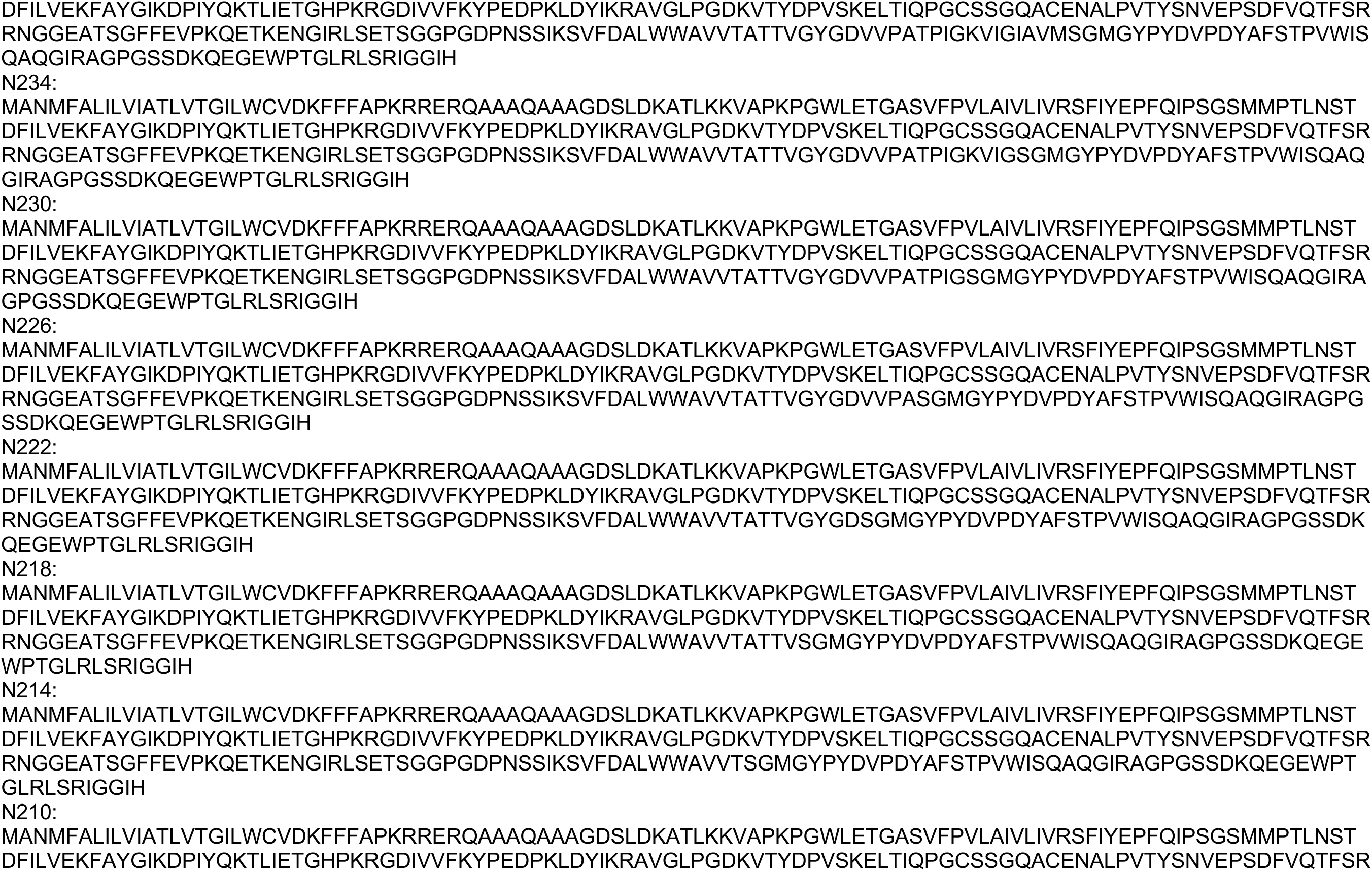
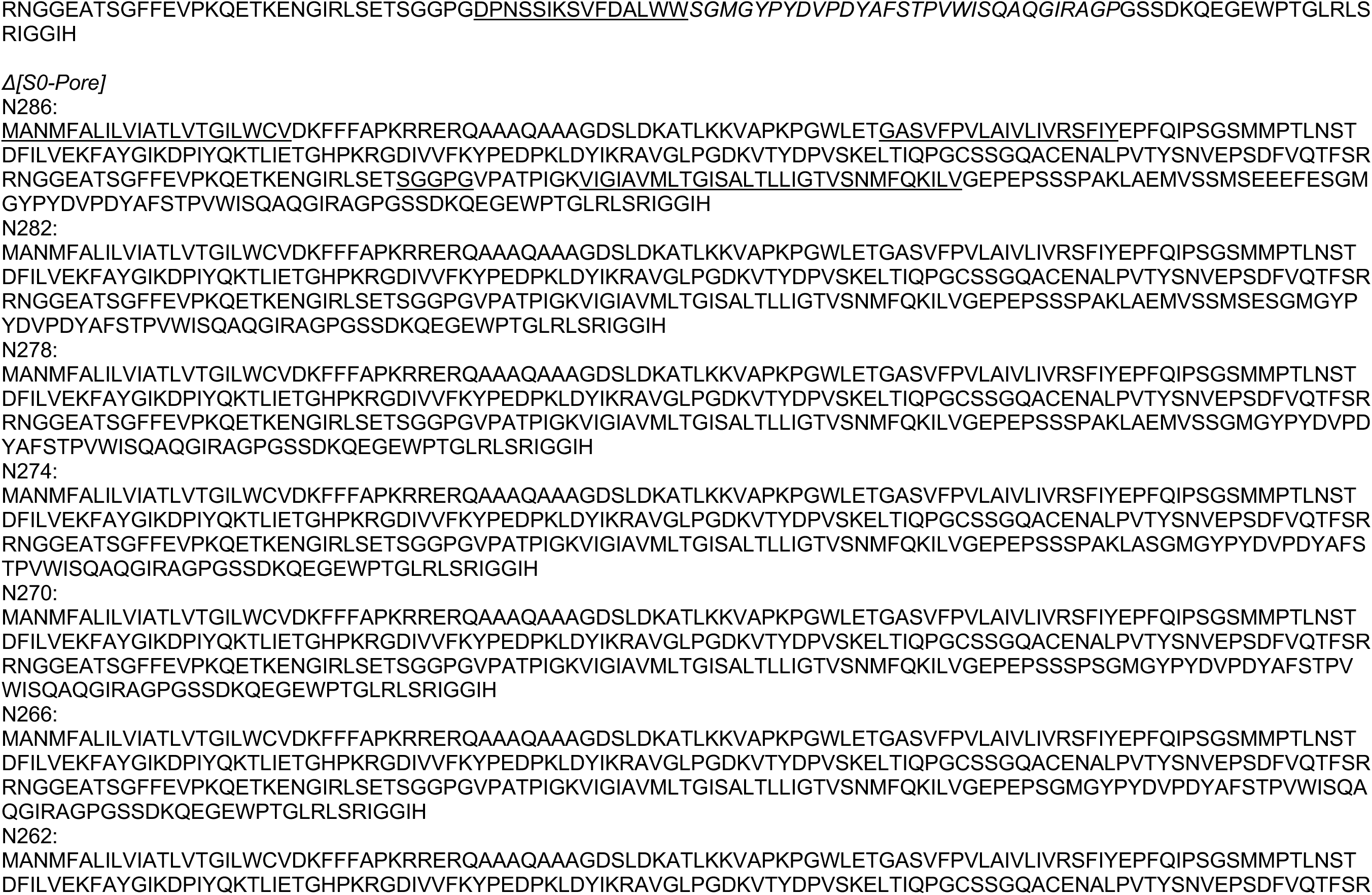
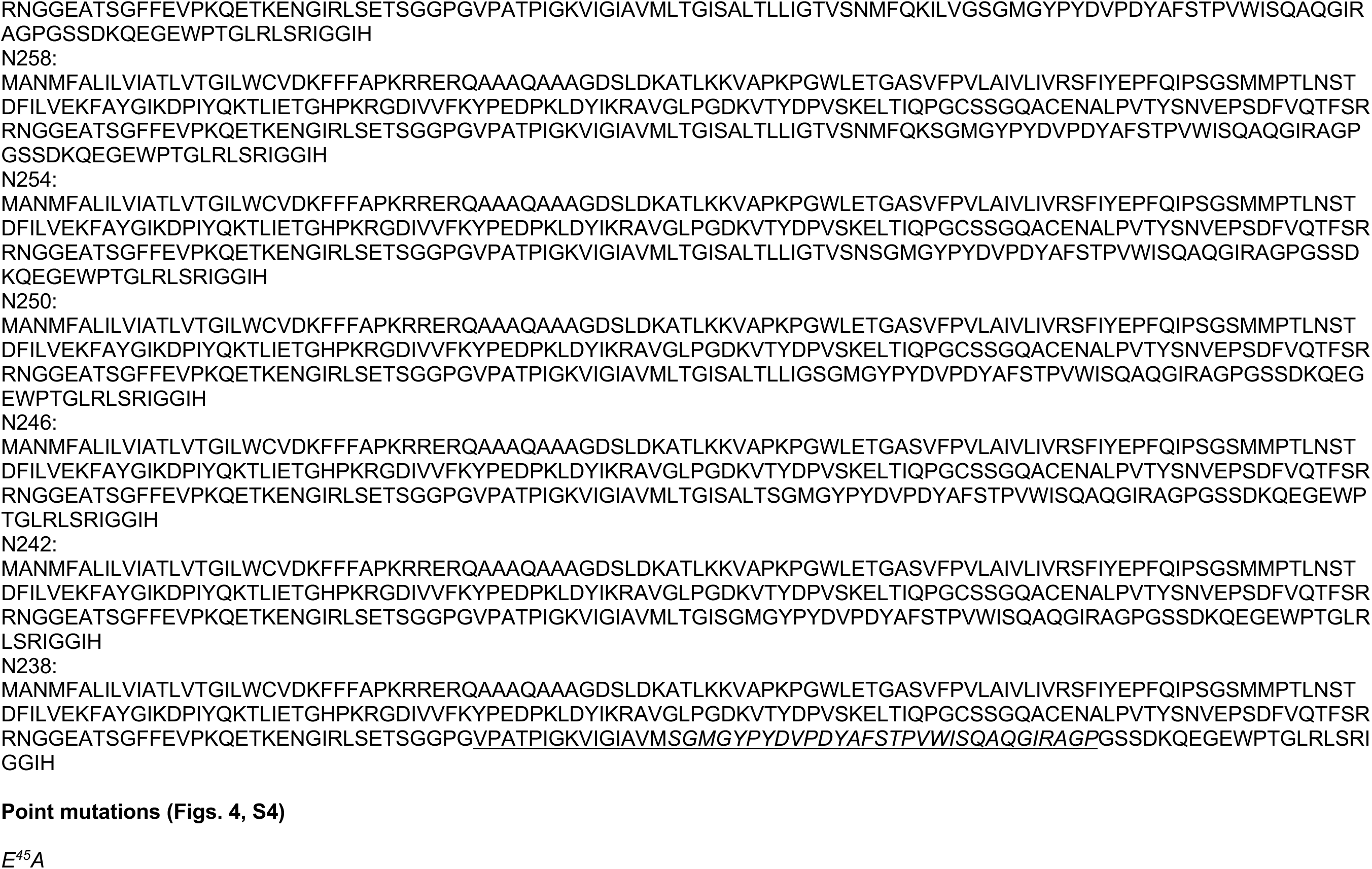
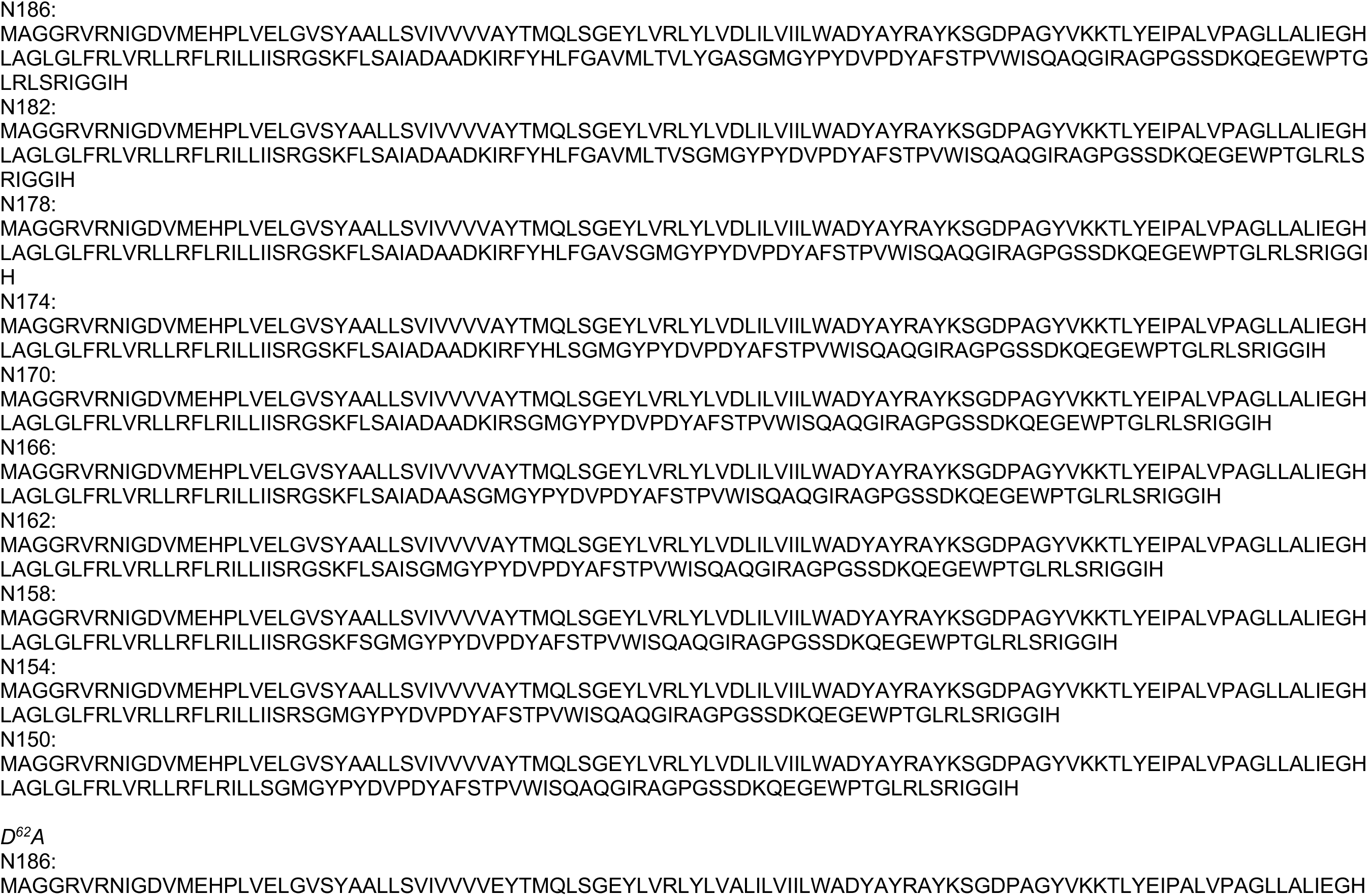
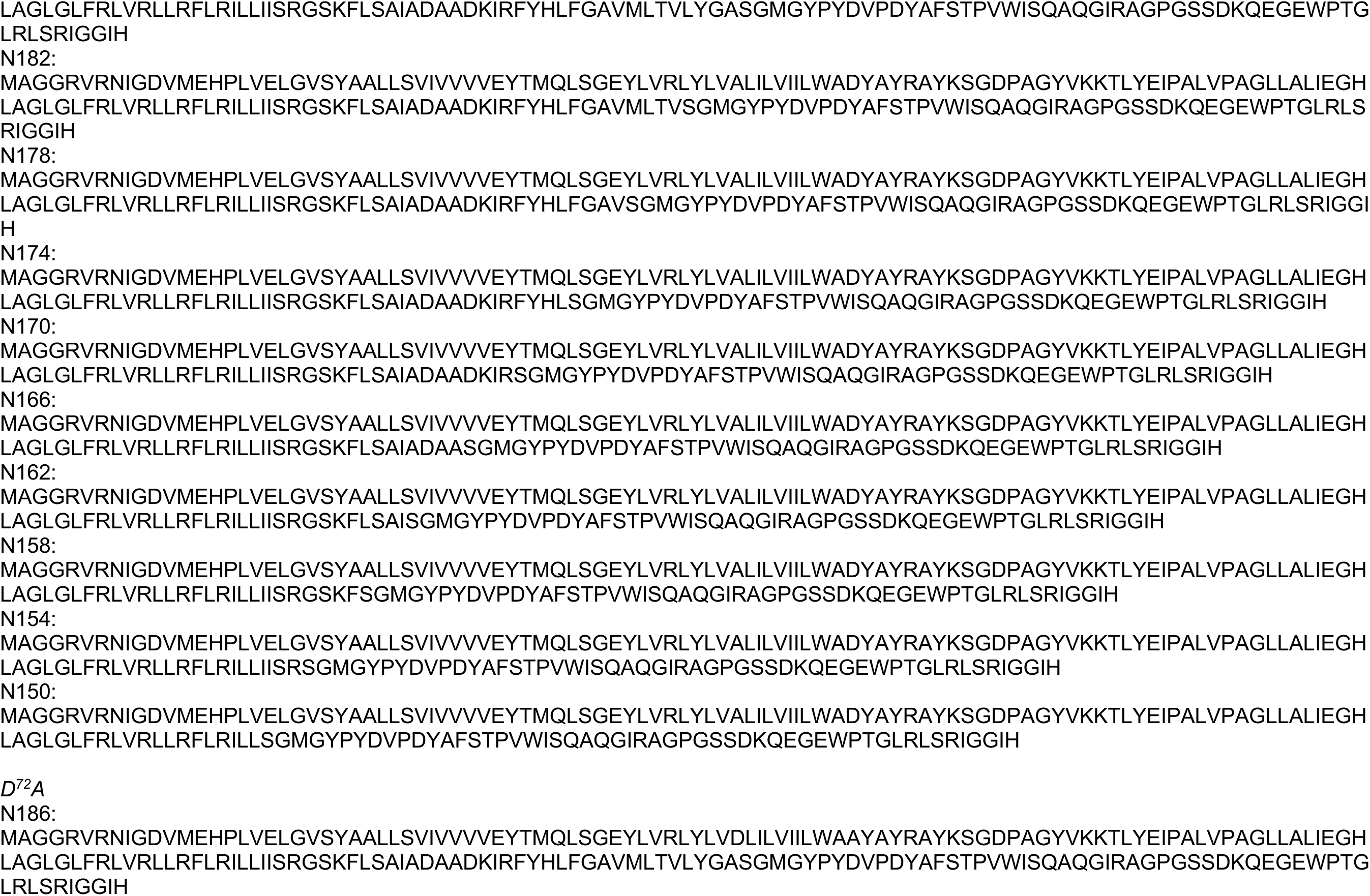
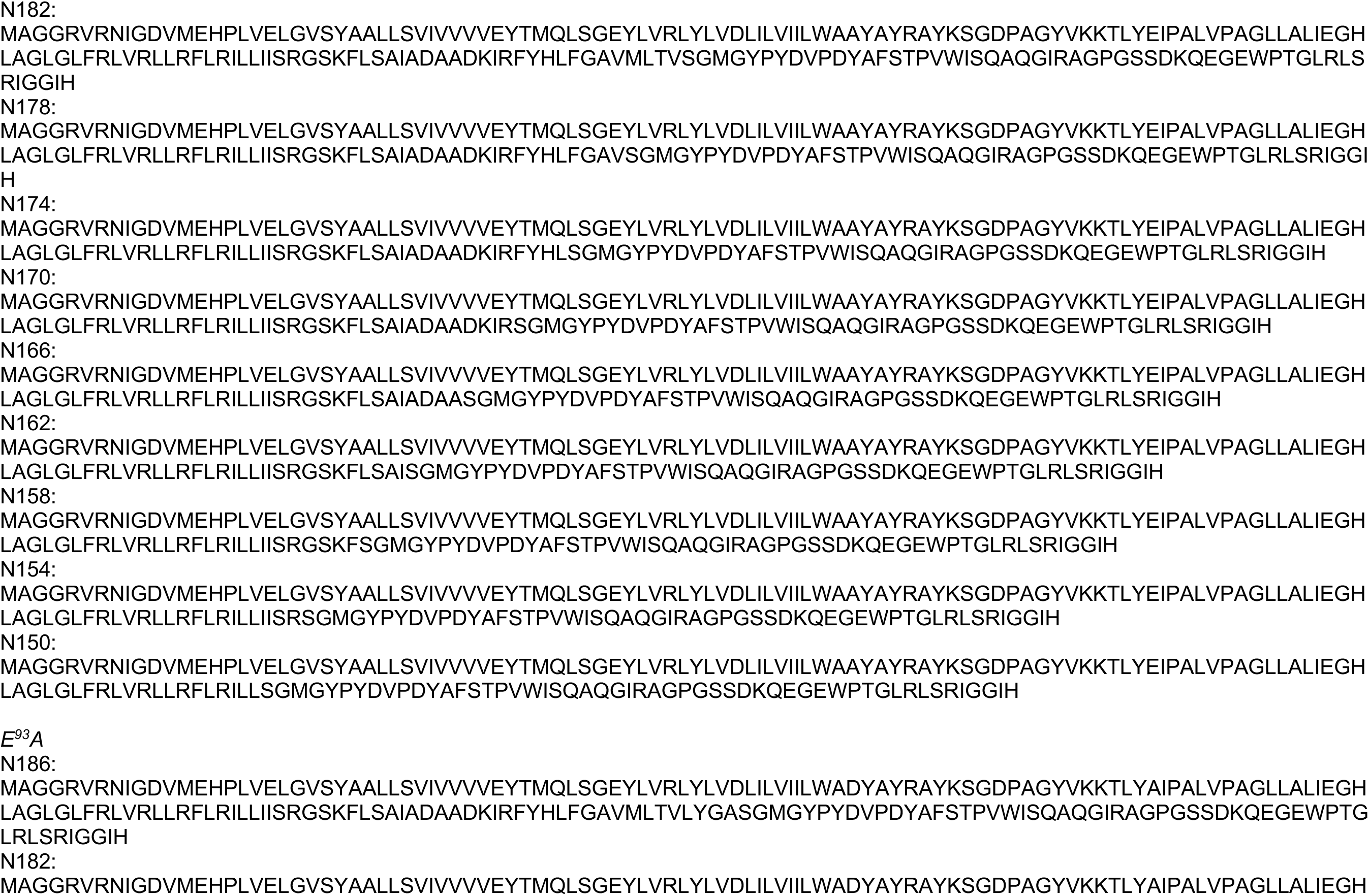
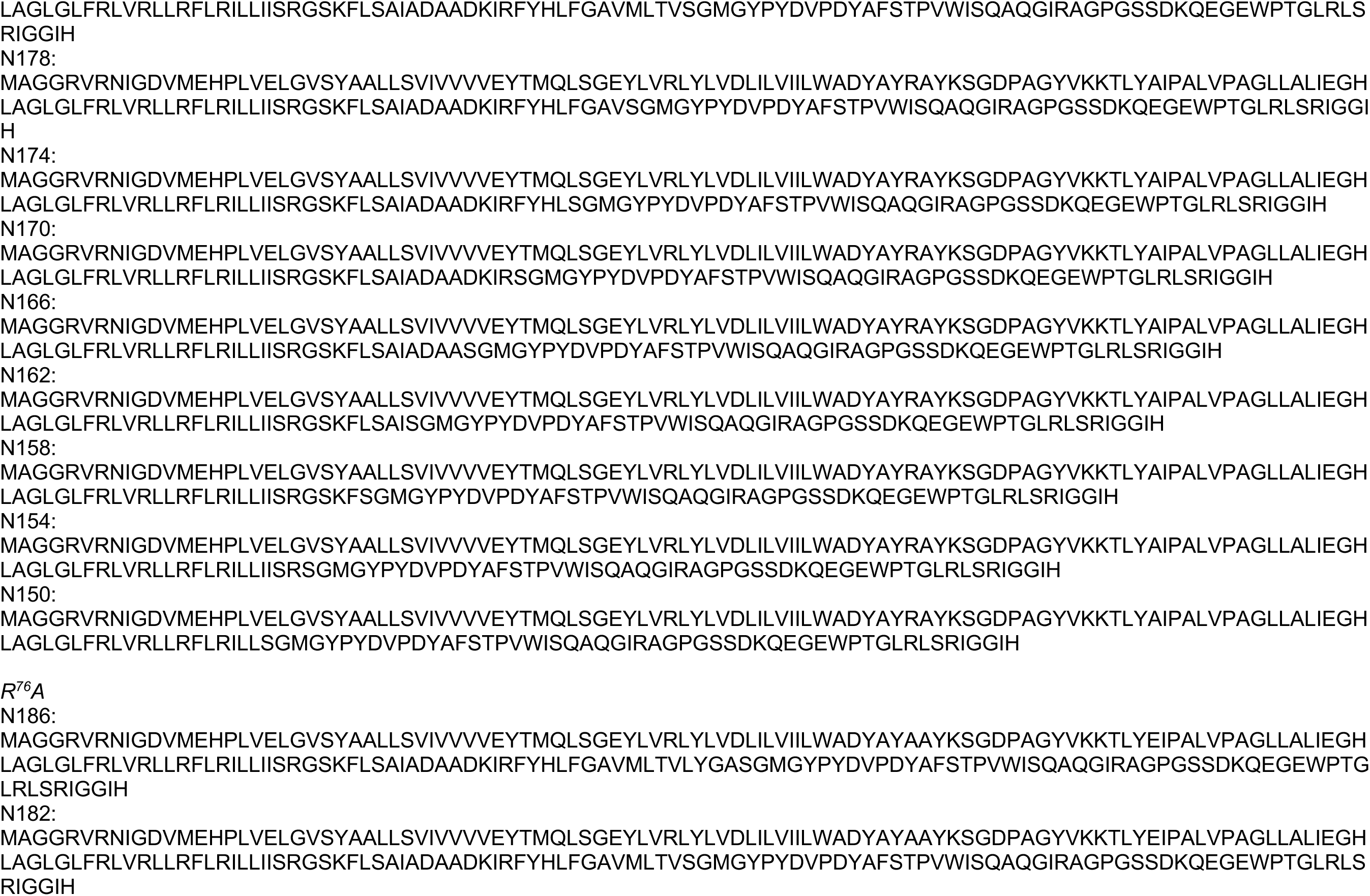
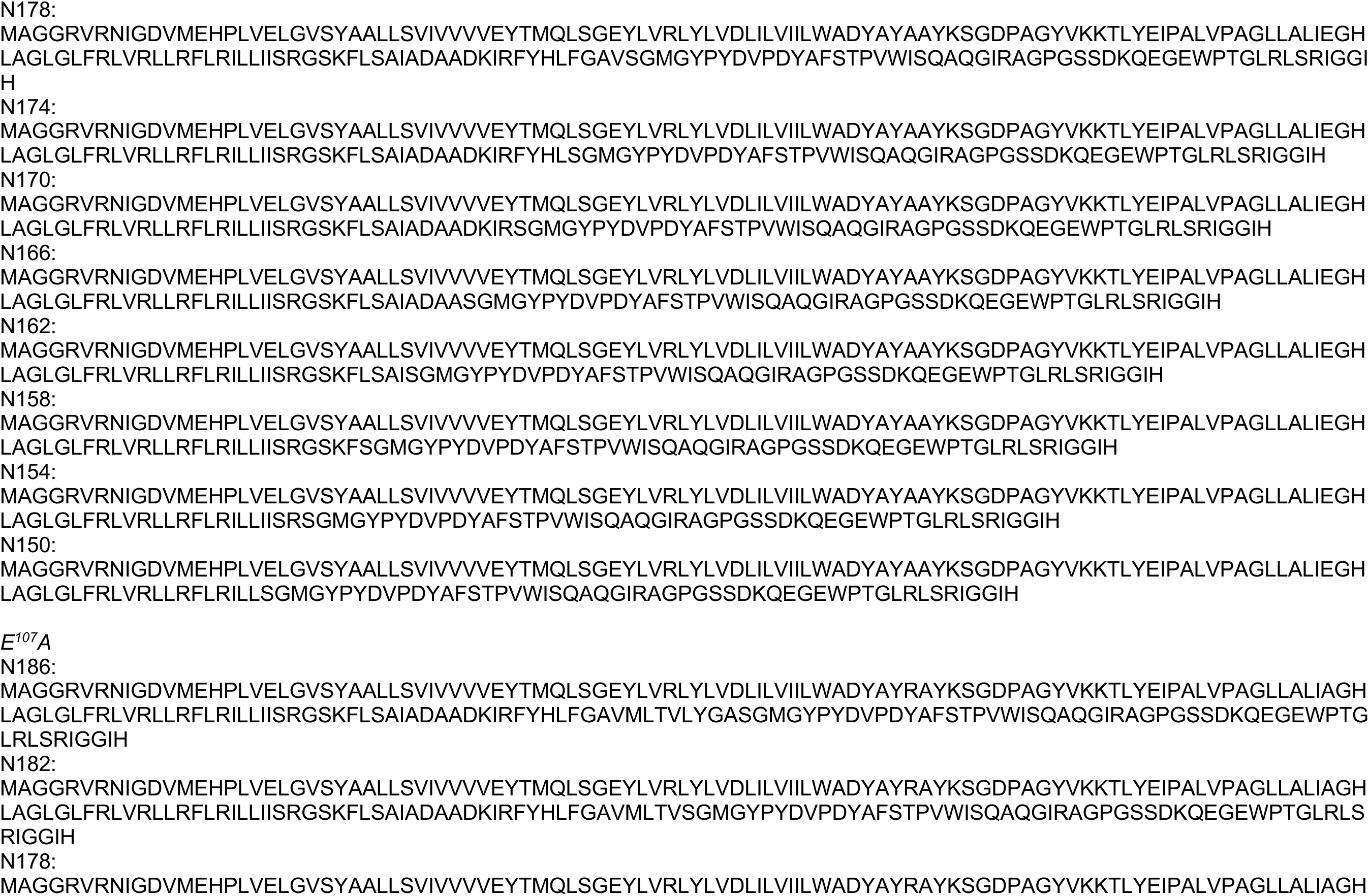
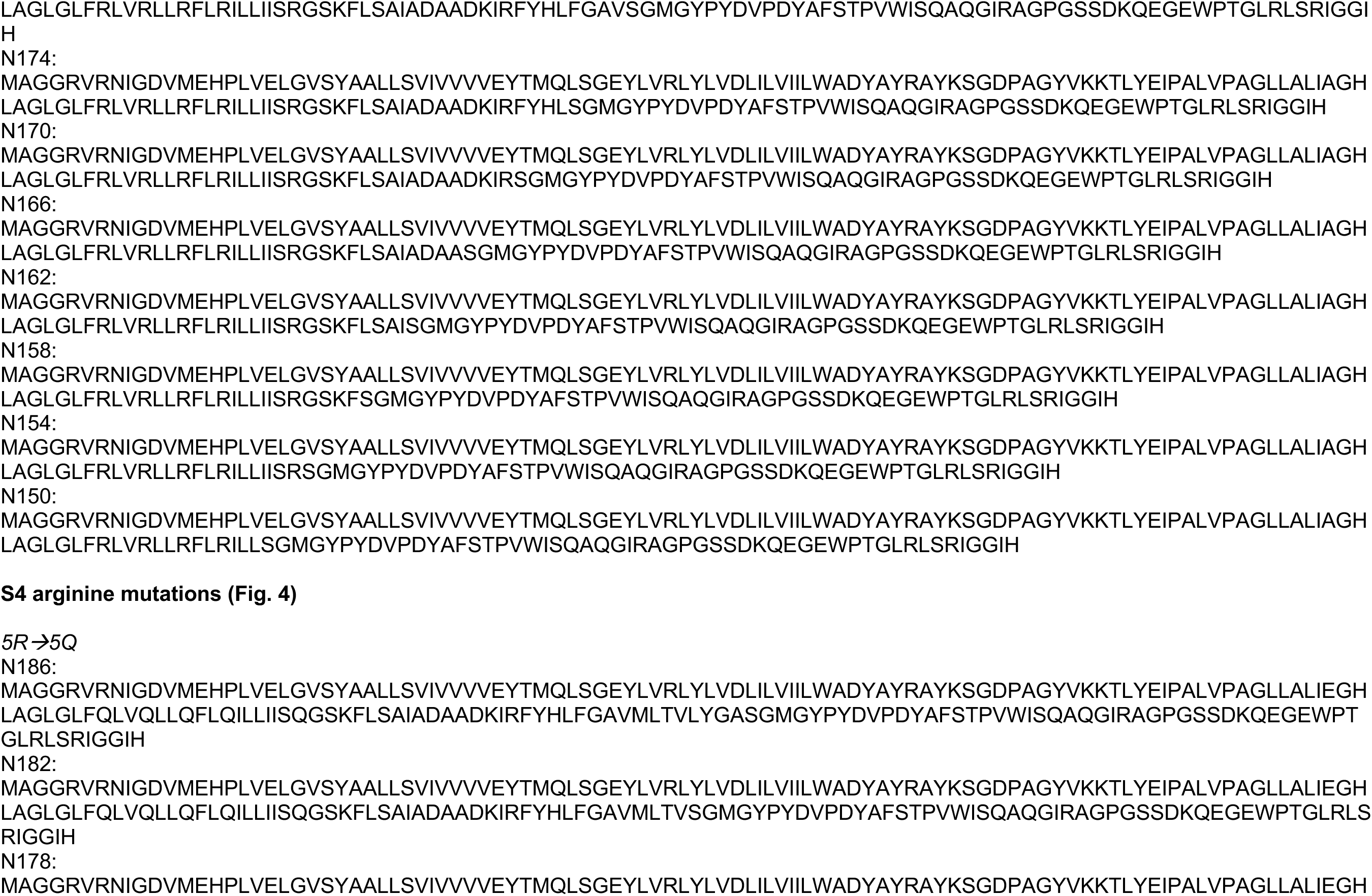
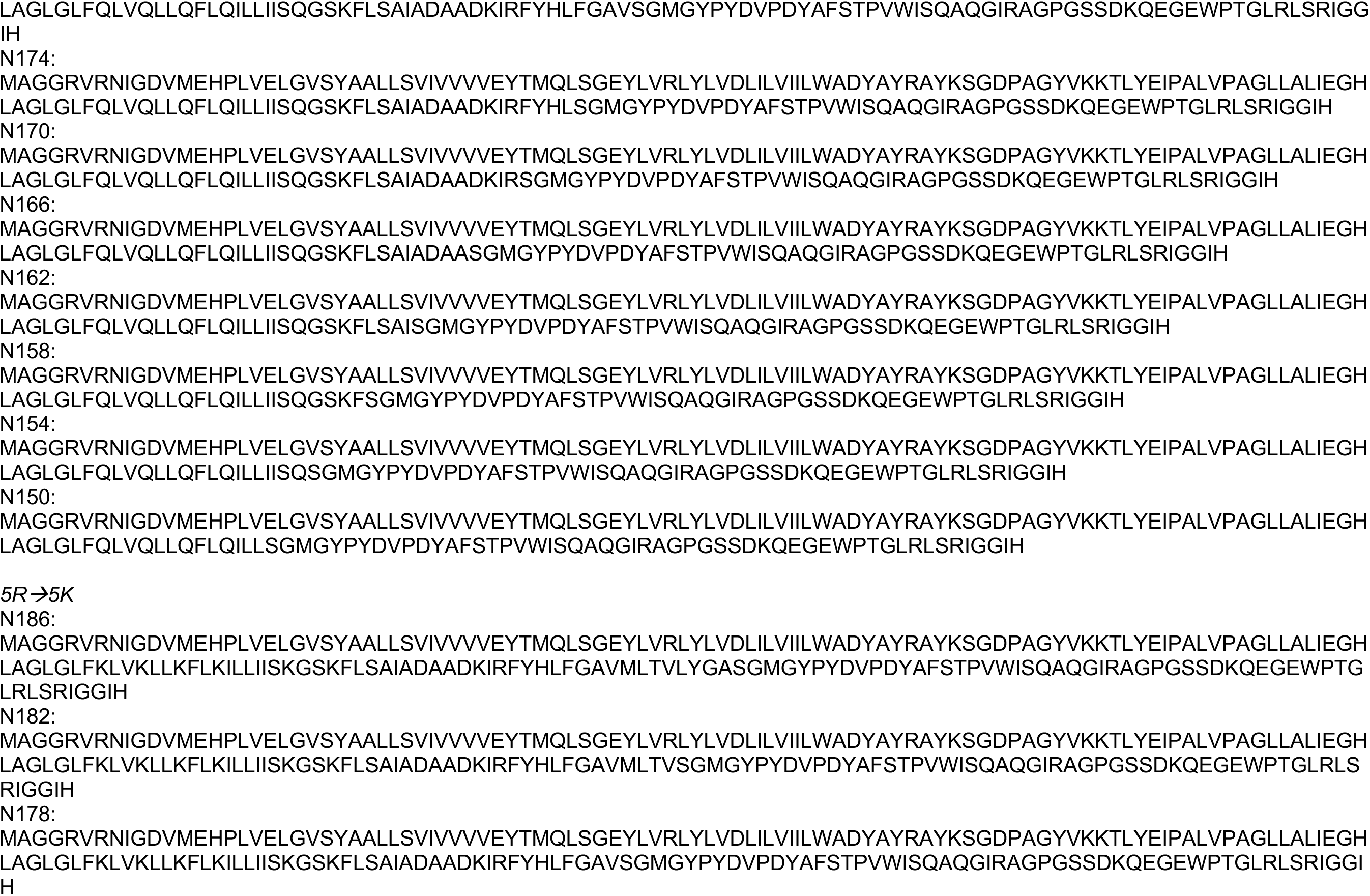
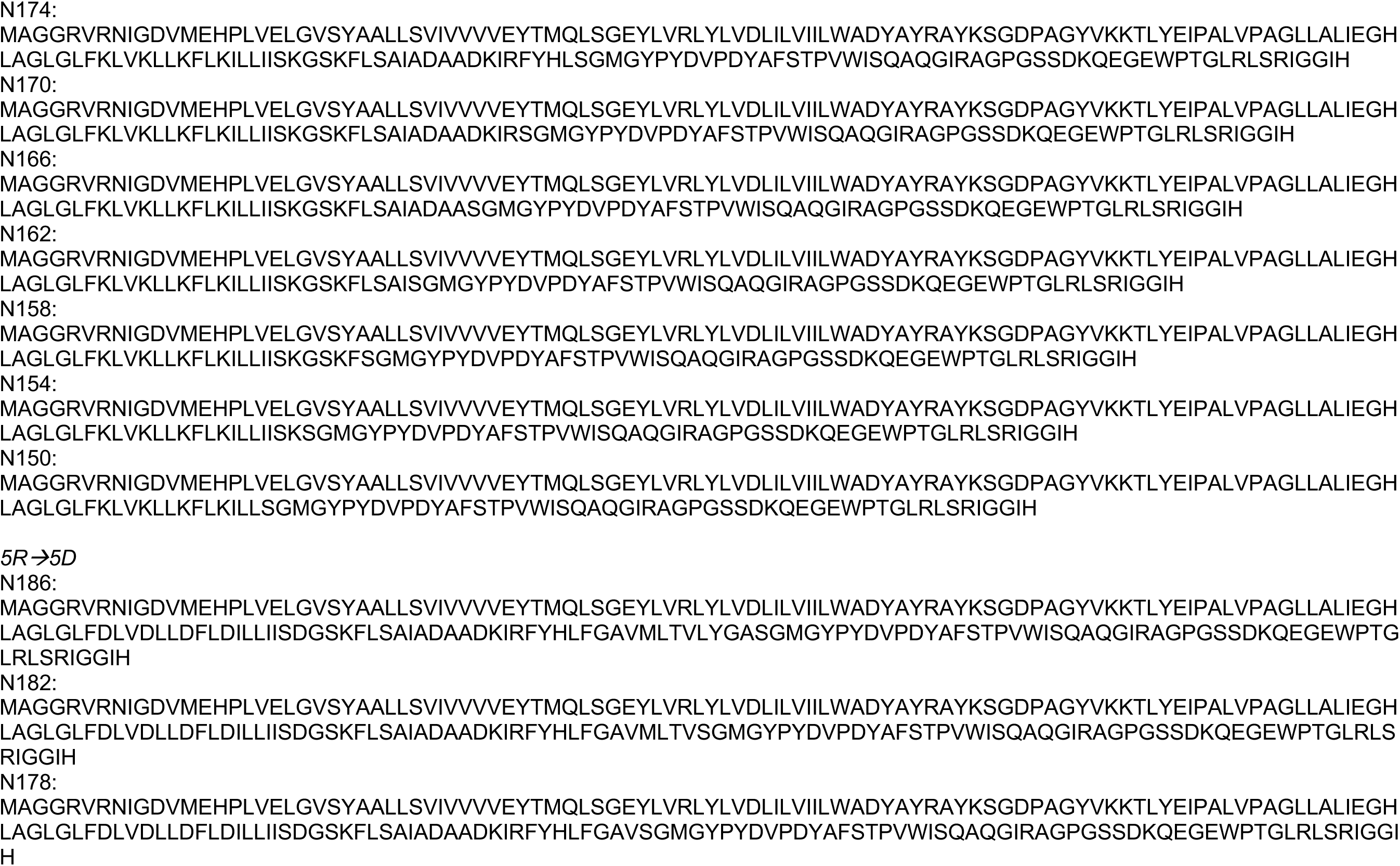
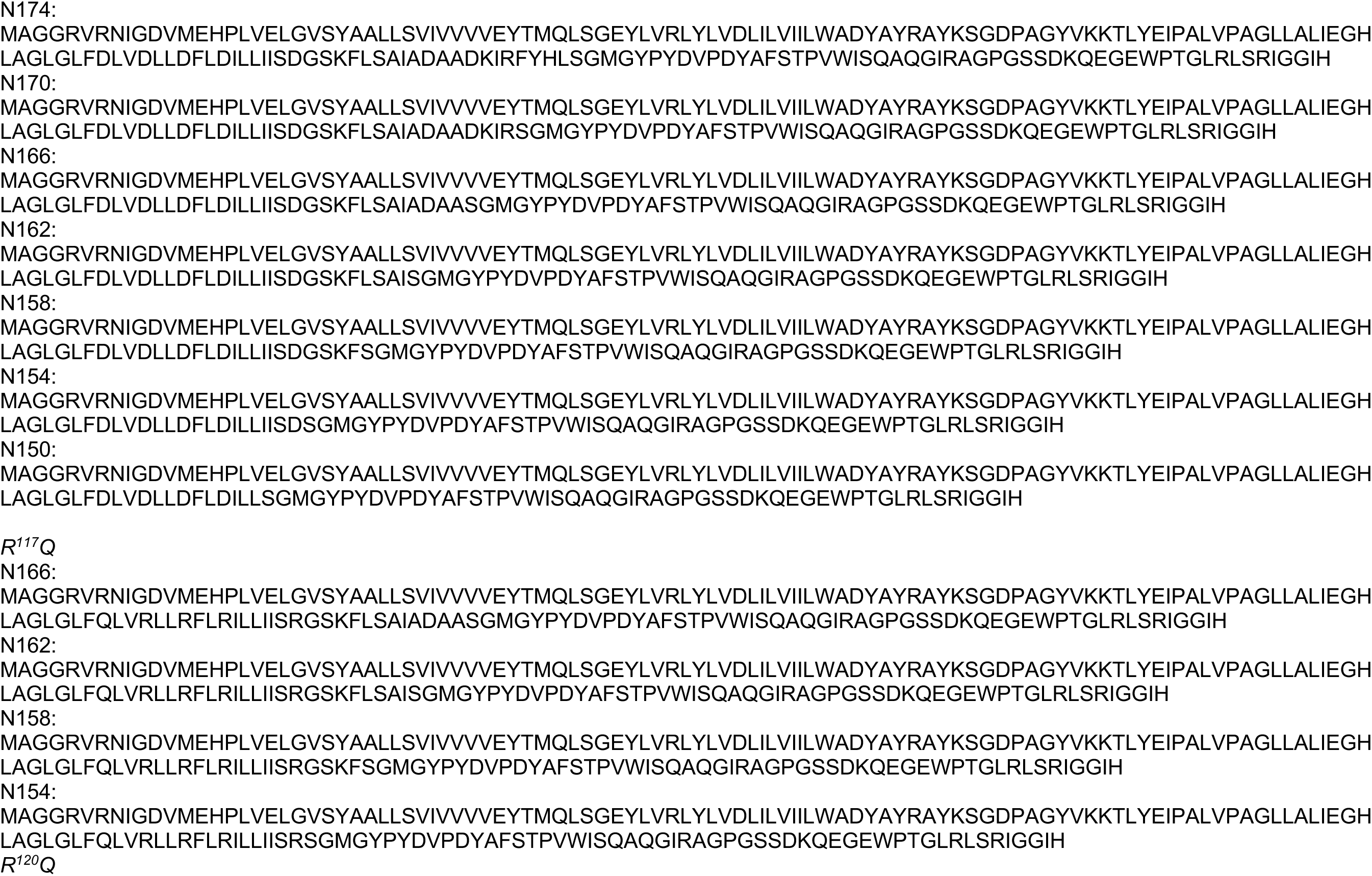
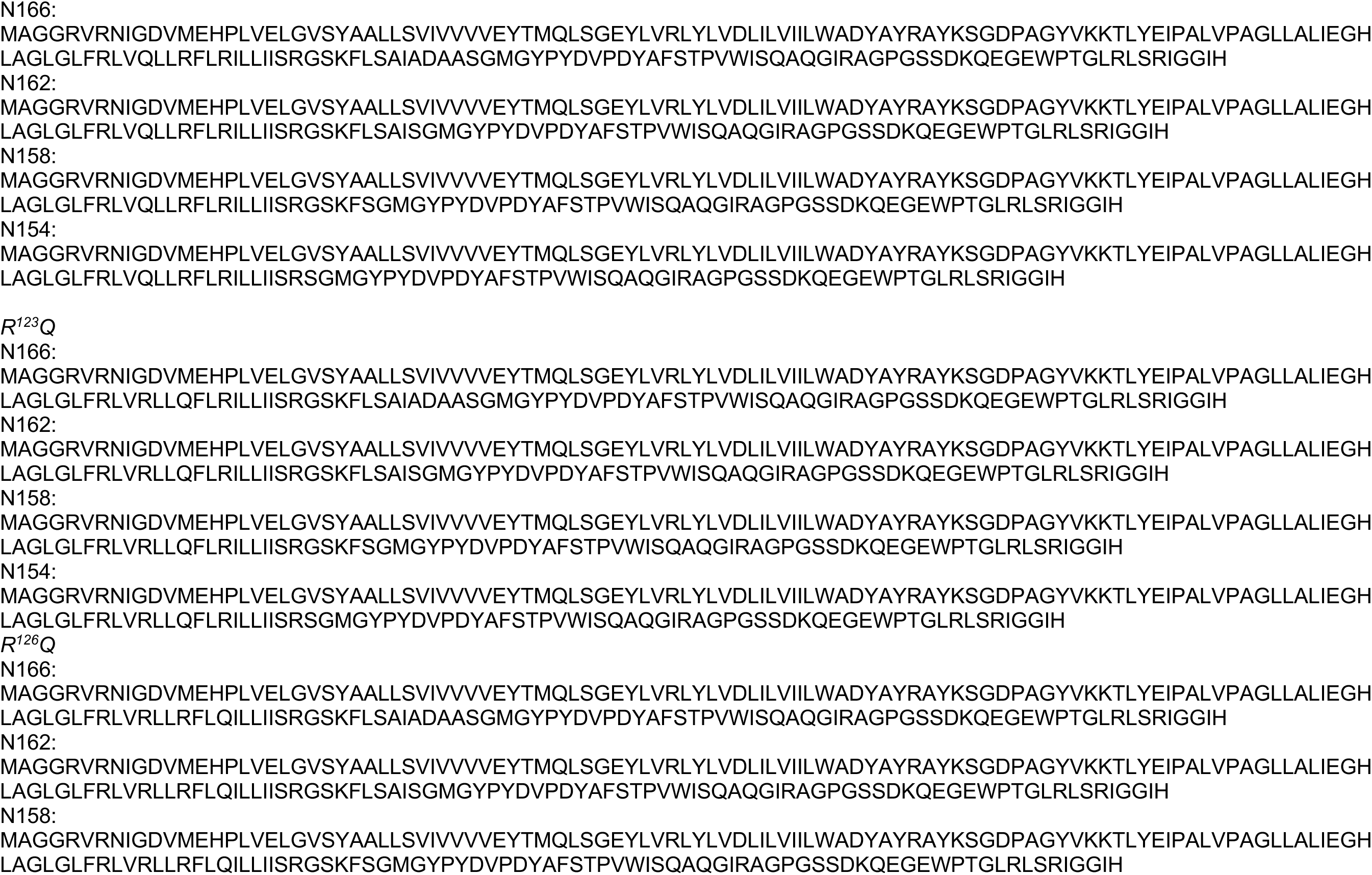
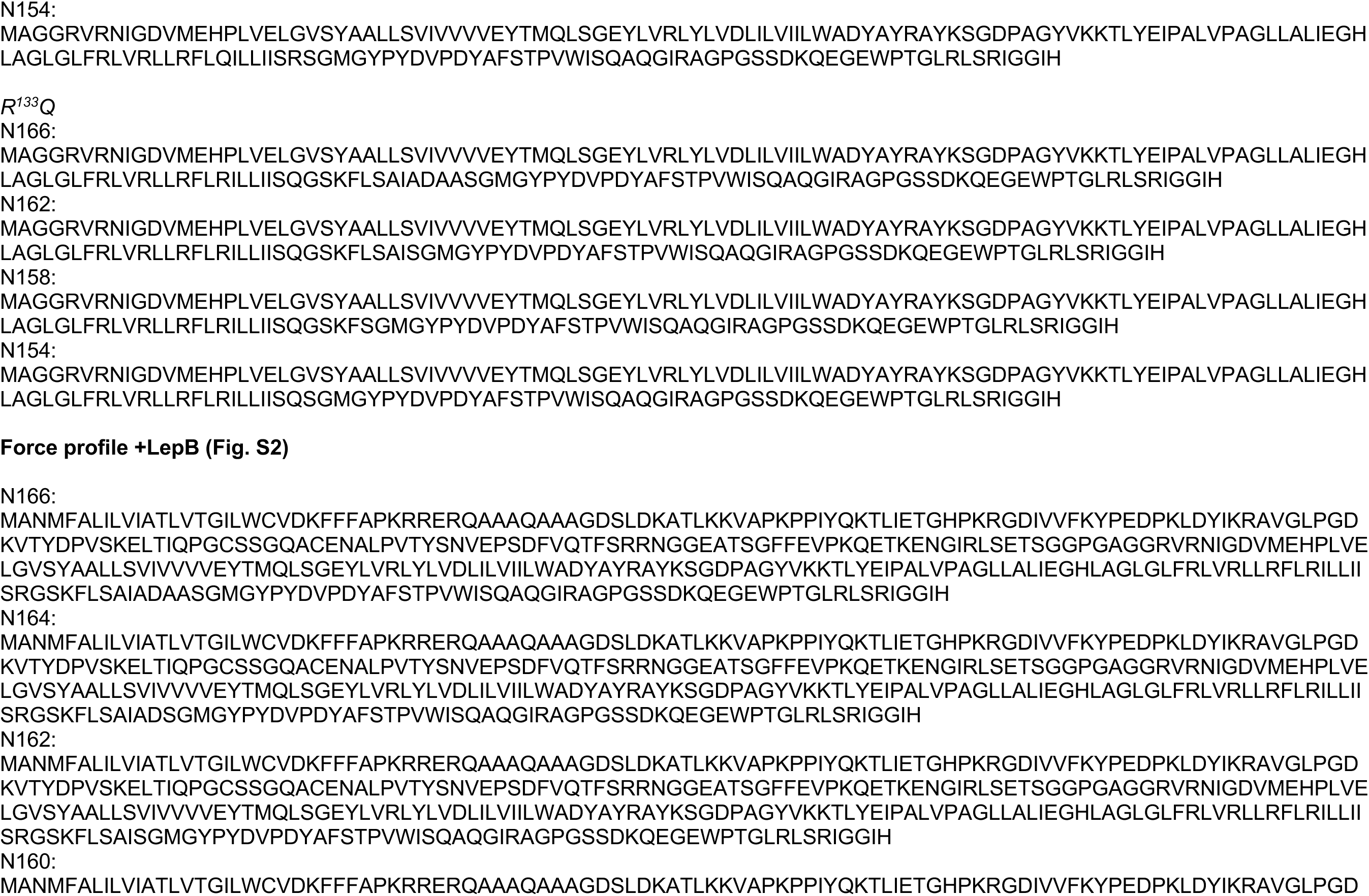
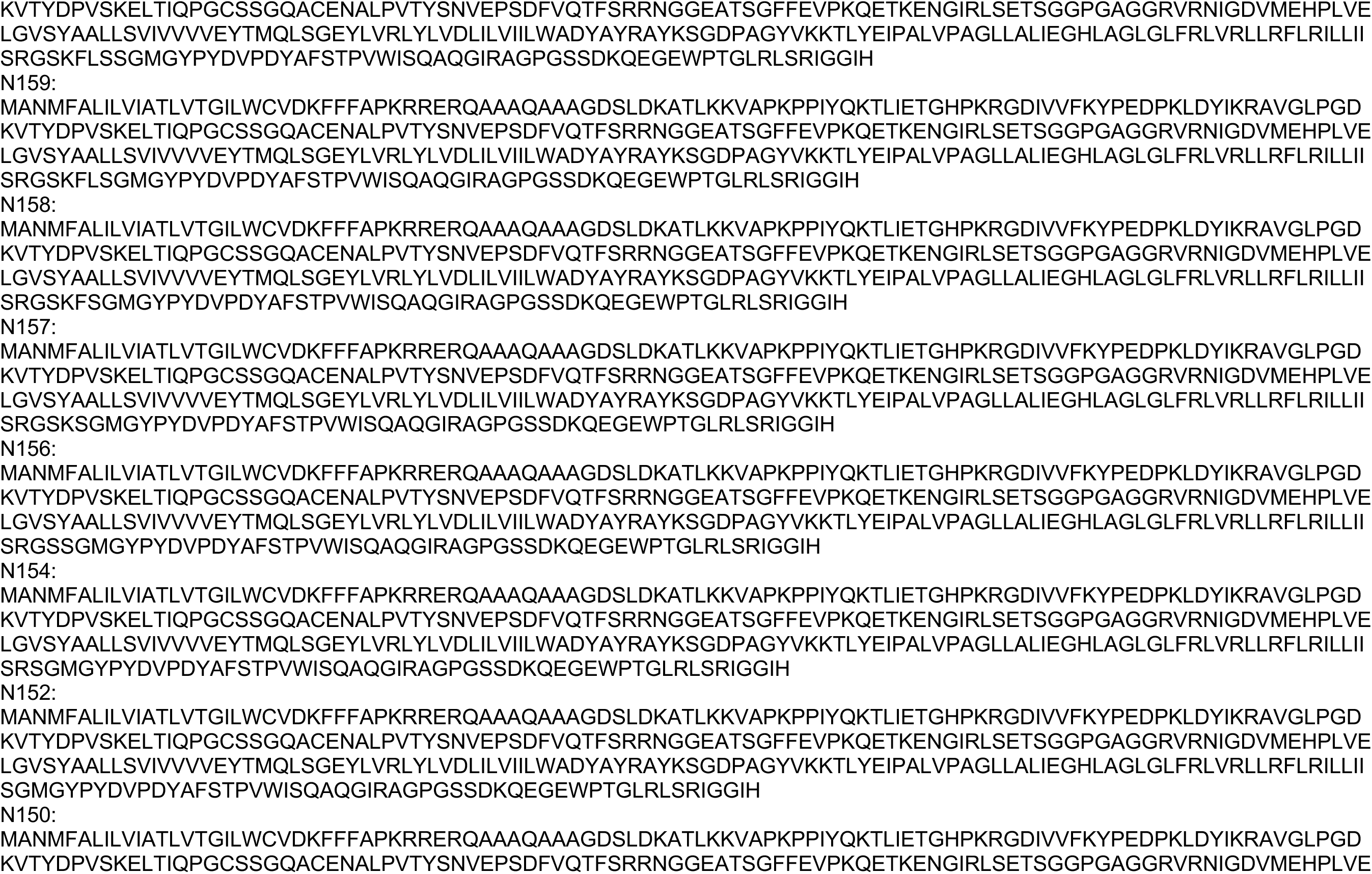
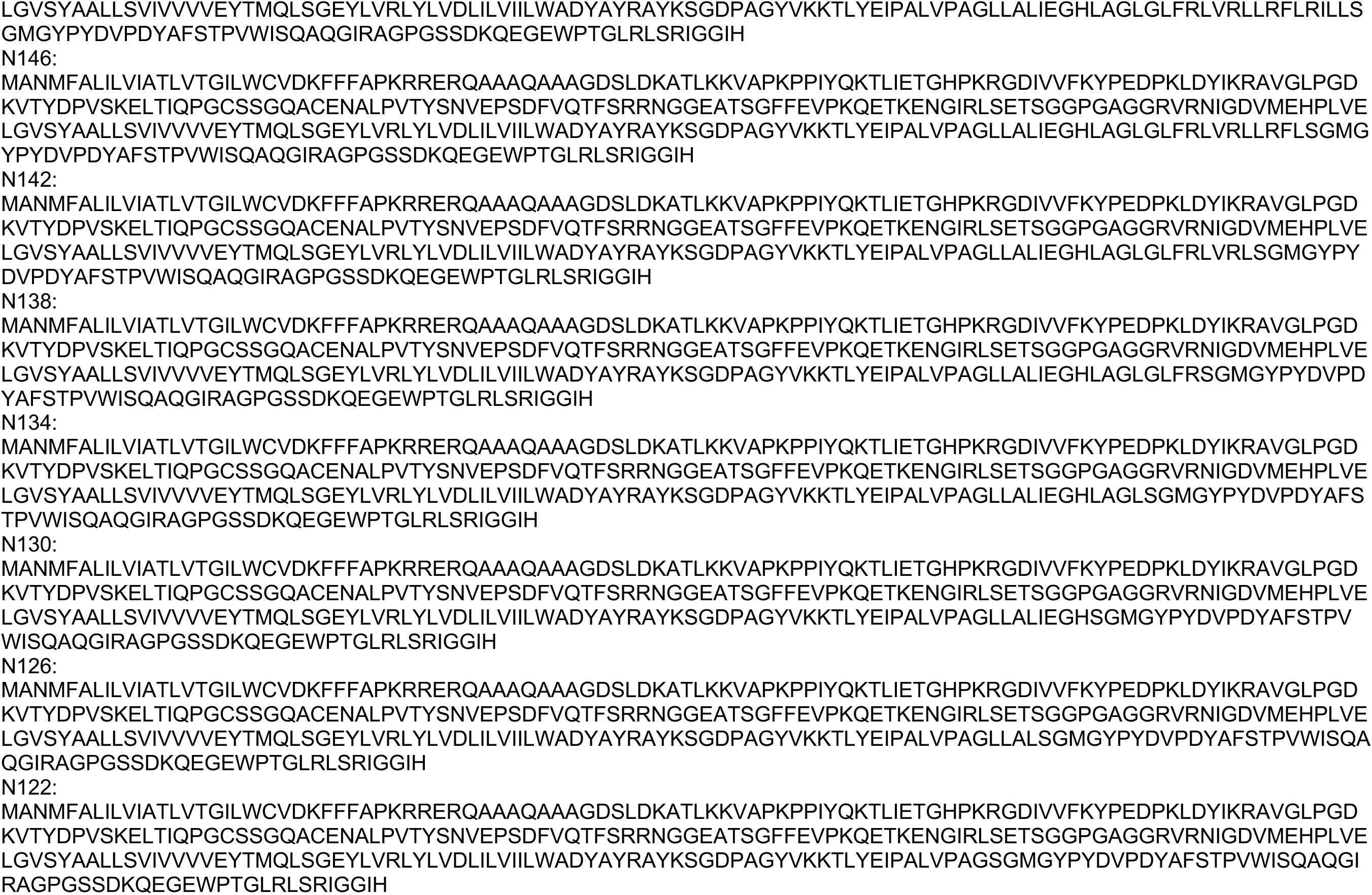
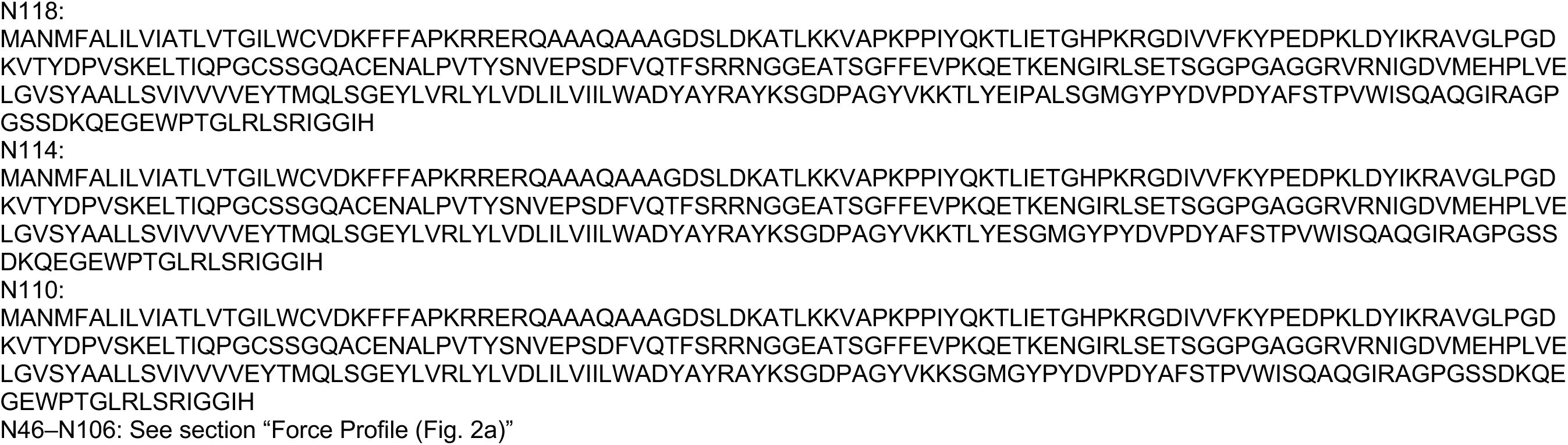
Amino acid sequences for all constructs.

